# Predicting Autopsy-Confirmed Neuropathology across Clinical, Neuroimaging, and CSF Biomarkers using Machine Learning

**DOI:** 10.64898/2026.05.20.726332

**Authors:** Christopher Patterson, Tamoghna Chattopadhyay, Sophia I. Thomopoulos, Andrew J. Saykin, Christos Davatzikos, Elizabeth C. Mormino, Duygu Tosun, Gary W. Beecham, Sarah A. Biber, Walter A. Kukull, Shannon L. Risacher, Thomas J. Montine, Sterling C. Johnson, Li Shen, Heng Huang, Guray Erus, Gyungah R. Jun, Shubhabrata Mukherjee, Paul K. Crane, Michael L. Cuccaro, Derek B. Archer, Bennett A. Landman, Arthur W. Toga, Timothy J. Hohman, Paul M. Thompson

## Abstract

Accurate in vivo prediction of neuropathology is critical for advancing diagnosis and treatment of Alzheimer’s disease and related dementias (ADRDs). As many individuals with ADRDs have mixed pathologies (β-amyloid, pathologic tau, cerebrovascular disease, vascular brain injury, pathologic TDP-43, hippocampal sclerosis, Lewy bodies), there is interest in determining how accurately we can infer these pathologic changes from clinical data, biofluid assays (e.g., CSF), and neuroimaging. Here we evaluated automated machine learning models trained on data curated by the AD Sequencing Project Phenotype Harmonization Consortium (N=7,894 individuals), to predict 26 autopsy-confirmed neuropathological outcomes. Predictors included in vivo clinical and cognitive composite scores, brain measures from 3D structural MRI and diffusion tensor imaging, image-derived measures of white matter hyperintensities (WMH), and CSF biomarkers. Predictive models were trained using ensemble learning with stratified cross-validation. We assessed performance using Spearman’s rank correlation and Matthews correlation coefficient, to accommodate co-occurring pathologic changes. The added value of neuroimaging and CSF versus clinical features alone was quantified. Braak stage was among the most consistently predicted outcomes. CSF biomarkers best predicted β-amyloid and tau pathology, but diffusion MRI metrics best captured vascular brain injury and white matter injury, and outperformed clinical and cognitive measures and anatomical MRI in predicting Lewy body disease. Anatomical measures from structural MRI outperformed standard clinical assessments in assessing neurodegeneration and hippocampal sclerosis, and WMH complemented cognitive measures in predicting TDP-43 pathology. These results establish a baseline for comparing modalities for inferring neuropathology.

## 1. Introduction

Alzheimer’s disease (AD) and the commonly co-morbid AD related dementias (ADRD) represent one of the greatest global health challenges of our time, and their prevalence is projected to triple by 2050 [Jellinger, 2022; Nichols et al., 2022; Alzheimer’s Association, 2023]. Biological AD is now formally defined by the abnormal accumulation of β-amyloid and tau pathology [Jack et al., 2018; Jack et al., 2024], but more broadly, ADRD encompasses a heterogeneous spectrum of prevalent neurodegenerative and cerebrovascular conditions that frequently co-occur in the same individuals [Schneider et al., 2007; Wilson et al., 2013; Boyle et al, 2018]. In many patients, AD neuropathologic change is accompanied by Lewy body disease (LBD), TDP-43 proteinopathy, hippocampal sclerosis, cerebrovascular disease (CVD), and vascular brain injury (VBI), in various combinations and possibly proceeding at different rates [Weller et al., 2009; Nelson et al., 2011; Thomas et al., 2020; Rabin et al., 2022; Woodworth et al., 2024].

If the underlying mixture of neuropathologic changes is not determined, patients may receive treatments that fail to address all of the primary biological drivers of their symptoms. Anti-amyloid immunotherapies, for example, may provide limited clinical benefit when symptoms are predominantly driven by ADRDs [van Dyck et al., 2023; Sperling et al., 2023; Sims et al., 2023; Tosun et al., 2024; Kapasi et al., 2025]. The high prevalence of mixed pathology in AD/ADRD may also have led to the failure of many late-stage clinical trials that target single molecular mechanisms [Cummings et al., 2014; Mehta et al., 2019; Duara et al., 2022; Wang et al., 2024]. Stratifying participants by neuropathological subtype may boost statistical power in drug trials by reducing biological heterogeneity within treatment groups. Even post hoc stratification may help to uncover therapeutic effects that are masked or diluted in mixed cohorts. Likewise, genomic and proteomic analyses within biologically defined subgroups may identify disease mechanisms more efficiently and improve power to detect meaningful associations.

Autopsy is still the gold standard to define and quantify neuropathologic features. Although the direct assessments used at autopsy cannot stratify patients during life, an increasing number of research cohorts are now being assessed with both ante mortem biomarker data and post mortem neuropathology. These datasets have stimulated efforts that attempt to predict neuropathologic features from data available in living patients. Positron emission tomography (PET) and cerebrospinal fluid (CSF) assays can detect β-amyloid and tau pathology during life and are now incorporated into recommended disease categorization frameworks for research studies of AD/ADRD [Jack et al., 2010; Jack et al., 2018; Jack et al., 2024]. Even so, PET and CSF assays are costly and invasive, limiting their use in large-scale screening and longitudinal monitoring. They also cannot capture the full range of co-pathologies observed at autopsy. Validated in vivo biomarkers are not yet available for TDP-43 pathology, Lewy body disease, and many forms of VBI that cause neurodegeneration [Dichgans et al., 2012; Thompson & Vinters, 2012; Wardlaw et al., 2013; McKeith et al., 2017; Nelson et al., 2019; Espay et al., 2025]. Although blood-based biomarkers such as ptau217 are now accurate for identifying AD [Hansson et al., 2022; Hannson et al., 2023; Ashton et al., 2024; Palmqvist et al., 2024; Therriault et al., 2024], and proteomic assays are rapidly improving, we still lack widely available methods to assess the presence and severity of ADRDs [Foulds et al., 2008; McMillan et al., 2023; Dark et al., 2024; Vrillion et al., 2024]. Because of this, there is great interest in developing machine learning methods that combine complementary, widely available data, including structural and diffusion MRI, clinical measures, and fluid biomarkers, to infer the presence and severity of multiple co-occurring neuropathologies [Tosun et al., 2024; Wang et al., 2025; Chattopadhyay et al., 2025a].

Among non-invasive imaging methods, structural MRI can detect cortical thinning and regional atrophy, and diffusion MRI can assess white matter degeneration. These methods lack molecular specificity, but the patterns they capture reflect the cumulative effects of neurodegeneration, CVD and VBI, and they may encode, indirectly, subtle information on the pattern and propagation of amyloid and tau pathology in the brain [Dickerson et al., 2009; Acosta-Cabronero et al., 2017; Chandio et al., 2024; Xu et al., 2026]

Machine learning and deep learning have increasingly been applied to multimodal MRI and PET data to infer underlying neuropathology. These approaches have been used to estimate spatial distributions of amyloid β and pathologic tau deposition [Chen et al., 2019; Jin et al., 2023] and to develop classifiers that distinguish AD, vascular dementia, and LBD [Makkinejad et al., 2021; Chattopadhyay et al., 2022; Dhinagar et al., 2023a; Dhinagar et al., 2023b; Chattopadhyay et al., 2024; Kapasi et al., 2025; Wang et al., 2025]. Tosun et al. developed an MRI-based multilabel classifier that detected co-occurring pathologic features of ADRDs, including pathologic TDP-43, LBD, and one form of CVD, cerebral amyloid angiopathy, with up to 93% accuracy in an autopsy-confirmed cohort [Tosun et al., 2024]. In Chattopadhyay et al. [2025a], we trained computer vision models on structural brain MRI along with routinely collected clinical data to predict multiple co-occurring neuropathologic features. More recently, deep learning analysis of pathology-confirmed datasets suggests that structural MRI captures characteristic, but somewhat overlapping, spatial signatures associated with AD, VBI, and LBD. Imaging-derived indices predicted neuropathology with fair accuracy (AUC ≈ 0.75–0.90), suggesting that T1-weighted MRI may help distinguish mixed pathologies [Wang et al., 2025; Kumar et al., 2025], although combining multimodal neuroimaging with other biomarkers and cognitive composite scores may be even better.

A key gap in the literature is the lack of large-scale, harmonized benchmarks that evaluate how well we can predict a broad range of AD/ADRD neuropathologic features from a combination of imaging, clinical, and biofluid data, in an effort to assess the added value of each type of data. Many prior machine learning studies focus on a single task - for example, predicting amyloid positivity [Chattopadhyay et al., 2024; Chattopadhyay et al., 2023a; Chattopadhyay et al., 2023b] - but relatively few have been validated across multi-cohort datasets with in vivo biomarkers and neuropathological ground truth. A range of algorithms, from classical machine learning to deep learning and computer vision methods [Chattopadhyay et al., 2025a], can be applied to this problem. Automated machine learning (AutoML) offers an approach to integrate multimodal data by systematically combining multiple algorithms while minimizing the need for manual model selection and tuning [Erickson et al., 2020; He et al., 2021; Gijsbers et al., 2024].

Given the ongoing interest in identifying multiple co-occurring neuropathologies that contribute to cognitive decline, in this study, we set out to evaluate how well automated machine learning could predict 26 post-mortem neuropathological features. To train the models, we benefited from the availability of carefully curated, harmonized ante mortem data from the AD Sequencing Project Phenotype Harmonization Consortium (ADSP-PHC) [Hohman et al., 2023]. We examined 12 predictor sets ranging from routinely collected clinical and imaging variables to non-invasive structural MRI measures (sMRI, dMRI) to protein-specific (yet invasive and expensive) CSF measures

Machine learning models were trained using a publicly available method, AutoGluon-Tabular [Erickson et al., 2020], in a stratified cross-validation framework. We evaluated predictive performance using the Matthews correlation coefficient (MCC) for binary outcomes and Spearman’s rank correlation for ordinal outcomes. We tested the added value of imaging and biomarker features beyond standard clinical information. This benchmark across 26 neuropathological features was intended to determine which types of commonly collected data would be most useful for inferring mixed pathology, with implications for improving diagnosis, clinical trial enrichment, and prioritization of future biomarker development for AD and ADRDs.

## 2. Methods

### 2.1 Datasets & Preprocessing

Study Cohorts. Data analyzed for this study are publicly available from the AD Sequencing Project (ADSP) Phenotype Harmonization Consortium (PHC) (release NIAGADS NG00067.v15: 2024/12/16). The ADSP-PHC (U24 AG074855) is an NIH-funded harmonization consortium, initiated in 2021, to harmonize phenotypic and biomarker data across NIH-funded research cohorts on ADRD with whole genome sequencing data [Hohman et al., 2023]. The ADSP-PHC harmonizes cognitive, imaging, neuropathological, and biofluid data across 40+ deeply phenotyped ADSP cohorts, and the harmonized data is available from a central repository [Beekly et al., 2007; Sonnen et al., 2009; Bennett et al., 2012; Barnes et al., 2012; Bennett et al., 2018; Hohman et al., 2023].

All component studies collected the original research study data with appropriate institutional ethics board approvals, using standard informed consent for participation in human subjects research.

For the current study, we focused on the subset of individuals with both postmortem and in vivo data (summarized in Table 1 and further detailed in Table 2).

**Table 1:** ADSP-PHC Cohorts analyzed in this study. This data is further detailed in Table 2. Note that the post-mortem neuropathology data from the ADNI cohort has not yet been harmonized with the rest of the PHC neuropathology dataset; its integration should boost sample sizes in future releases.

**Table 2:** Sample sizes and class structure for each neuropathological outcome across feature sets. Sample sizes (N) and class distributions for each neuropathological outcome and feature set combination. Variation in N reflects differences in data availability across modalities and outcomes. Class imbalance is quantified using normalized Shannon entropy (range 0–1), where lower values indicate greater imbalance. These factors influence model performance and should be considered when comparing predictive accuracy across outcomes.

Data used to predict neuropathology were restricted to the most recent time point prior to death, for participants who had at least one of the feature sets described below and who had a corresponding brain autopsy assessment. Sample sizes varied across feature sets and outcomes (Table 2), reflecting differences in the availability of imaging, biofluid, and cognitive measures across individuals and across cohorts. In later analyses, we performed sensitivity analyses to understand whether predictive performance differed across subsets defined by data availability.

### 2.2 Neuropathological Outcomes

Neuropathological outcomes were obtained from the ADSP PHC harmonized neuropathology dataset and, as detailed in Table 2, included 26 distinct measures organized across seven pathological domains: (1) amyloid pathology (Thal phase [Thal et al., 2002], A Score, CERAD score [Fillenbaum et al., 2008], and binary amyloid positivity); (2) tau pathology (AD Braak stage [Braak et al 2006], B Score); (3) cerebrovascular disease (CVD) (arteriolosclerosis [Montine et al., 2012], atherosclerosis, cerebral amyloid angiopathy (CAA) [Vonstattel et al., 1991; Olichney et al., 2000] ordinal and binary indicators, plus a composite CVD indicator); (4) hippocampal sclerosis (binary) [Nelson et al., 2013]; (5) LBD (Parkinson’s Disease (PD) Braak stage [Braak et al., 2003; McKeith et al., 2005; Braak et al., 2017]), L Score (ordinal), and binary Lewy body presence; (6) TDP-43 pathology (binary whole-brain presence, and 3- and 5-stage scales for assessing anatomic distribution of pathologic TDP-43 [Cairns et al., 2007; Josephs et al., 2014]; and (7) VBI (hemorrhages, infarcts, microinfarcts, composite vascular injury indicator [Vonstattel et al., 1991; Olichney et al., 2000]; white matter rarefaction (leukoaraiosis; WMR) [Alosco et al., 2019; Beach et al., 2023]. The neuropathology data dictionary defines binary WMR based on the presence of leukoencephalopathy; this can originate from VBI or Wallerian degeneration, but has been historically classified as cerebrovascular injury. Ordinal outcomes ranged from three to seven classes, depending on the staging system. The number of classes and class imbalance for each outcome and feature set are summarized in Table 2. The A Score, B Score, L Score, and pathologic TDP-43 (3-stage) are derived by condensing the number of ordinal classes from the Thal Phase, AD Braak Staging, PD Braak Staging, and pathologic TDP-43 (5-stage) scales, respectively.

#### 2.2.1 Class Imbalance

As noted in prior neuropathological studies [Wang et al 2025; Chattopadhyay et al 2025a], class imbalance is an important consideration; for rarer pathologies, we do not want to favor a classifier that simply predicts the majority class (no pathology). Because of this, measures of predictive accuracy need to account for class imbalance. Class (im)balance was quantified using the normalized Shannon Entropy:

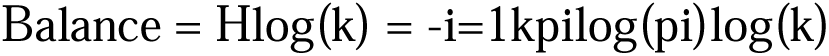

where H is the Shannon entropy, and pi is the proportion of the i-th class out of k total classes. This metric is scaled to the range [0, 1], where 1 indicates a perfectly balanced distribution (all classes equally represented) and 0 indicates complete imbalance (only one class represented).

### 2.3 Feature Sets

Twelve feature sets, individually and in combination, were used to predict neuropathological outcomes. These were grouped into four broad domains: (1) routinely collected clinical and demographic measures (“base clinical”), (2) 3D structural MRI (sMRI), white matter hyperintensities (WMH), and diffusion tensor imaging (DTI), (3) harmonized and co-calibrated cognitive domain scores, and (4) CSF biomarkers. The composition and counts for each feature set are summarized in Table 3. For feature sets with multiple time points available, we selected the last visit prior to their death as the cross-sectional basis for the analysis. In Table 3, we include the total number of features for each set as well as the effective dimensionality (ED) of that set and its outcomes, calculated by the spectral entropy of the eigenvectors [Cangelosi & Goriely 2007; Roy & Vetterli 2007] on the set of participants with complete data for both the features and outcomes. While each of the feature sets contained the same number of participants, the neuropathology outcomes did have missingness that led to excluded data points for different outcomes. The variation in sample sizes reported below reflects this data availability. The available sample size for each Feature Set - Outcome pairing is reported in Table 2.

**Table 3:** Feature set descriptions. ED denotes the effective dimensionality of each feature set (via spectral entropy), summarizing how many independent modes of variation are captured.

#### 2.3.1 Base clinical

The base clinical feature set included four variables: sex, age at death, diagnostic status at assessment (“normal”, “mild cognitive impairment (MCI)”, or “dementia”), and the ante-mortem interval between last assessment and death (ΔAge, also referred to as the “ante-mortem time”). For prospective use, this interval would correspond to the time between in vivo measurement and the point at which pathology is inferred. Clearly, this interval may influence class probabilities and prediction accuracy. The base clinical feature set had the largest sample size (N = 3,345-7,865; sex, age at death, cognitive status, and ΔAge were available for all cases) and served as the reference for comparison with imaging and biofluid models. All other feature sets were evaluated in combination with base clinical covariates, in proportions that reflect typical data availability.

#### 2.3.2 Structural MRI

Harmonized structural MRI features were derived from 3D volumetric T1-weighted brain MRI using the widely-used FreeSurfer software [Fischl and Dale, 2000; Fischl, 2012], harmonized by ComBat [Johnson et al., 2007; Fortin et al., 2018], and organized into four sub-types each tested separately: (1) Thickness - cortical thickness values for 69 regions of interest (ROIs; 73 features total); (2) Volumes - gray matter, subcortical white matter and CSF volumes for 111 ROIs (115 features); (3) rICV (ratio to intracranial volume) - all FreeSurfer volumetric measures divided by total intracranial volume (ICV) [Buckner et al., 2004], yielding 114 features; and (4) All sMRI Features - all three MRI sub-types combined into a single 302-feature set. All structural MRI feature sets had the same number of participants per outcome (N = 156 - 422; here the range covers the sample size for different neuropathological outcomes). Note that the volumetric ratios included each ventricle and a composite of all CSF, which are robust metrics of global atrophy. As such, they do not simply reflect tissue-occupying volume. We note that correlated ROI-level measures (e.g., thickness, volume, rICV) may introduce redundancy and potentially overweight certain regions. Even so, evaluating these feature types, separately and combined, allows us to assess any complementary and overlapping contributions. Ensemble learning methods are relatively robust to correlated inputs. Similarly, including both raw and ICV-adjusted volumes provides complementary representations, and the stronger performance of rICV features confirms normalization as the more informative signal.

#### 2.3.3 Cognitive composite scores

Cognitive composite scores consisted of 14 features including the base clinical covariates, the participant’s educational attainment in years [Stern et al, 2019], and measures derived from three harmonized cognitive domains: memory, executive function, and language [Mukherjee et al, 2023, Kang et al., 2025; Hohman et al., 2023]. Each domain was represented by a harmonized composite score, the standard error of that harmonization, and a binary precision filter indicating data quality. Cognitive composite scores had the second largest number of participants after the base clinical (N = 2,460 - 7,341). The harmonized cognitive dataset also included a visuospatial domain, but because one contributing cohort, NACC, did not include these composite scores until UDS3 [Besser et al 2018], most of the NACC cohort from UDS2 and earlier was missing the visuospatial scores, even if they had harmonized values in the other three cognitive domains. The availability of this test affected the N for all the outcomes by more than 3,000 participants.

#### 2.3.4 Cerebrospinal fluid (CSF) biomarkers

CSF biomarkers consisted of 14 features (10 biomarker features plus 4 base clinical covariates): raw and harmonized Z-scores for Aβ42, total tau, phosphorylated tau (pTau), the Symmetric Centipolar Normalized Score (SCeNS) for both Aβ42 and pTau, and a binary positivity indicator for Aβ42 and pTau [Timsina et al 2024]. CSF features were available in substantially smaller sub-cohorts (N = 39-281 per outcome; Table 2) than base clinical features. We included both raw and harmonized measures to preserve potentially informative biological variation while mitigating site effects; the model can selectively weight these representations, reducing the impact of redundancy.

#### 2.3.5 White Matter Hyperintensity

White Matter Hyperintensities (WMH) metrics included 10 variables harmonized from fluid-attenuated inversion recovery (FLAIR) MRI scans either directly measuring WMH or related to vascular disease: the signal-to-noise ratio (SNR) and contrast-to-noise ratio (CNR) for both gray and white matter, the contrast difference between gray and white matter, and the harmonized white matter lesion volume [Peter et al 2025].

#### 2.3.6 Diffusion tensor imaging (DTI)

DTI features included the four standard scalar metrics, each derived from the Johns Hopkins Eve Type 3 white matter atlas (95 ROIs) [Mori et al., 2008; Oishi et al., 2009] with ComBat harmonization applied [Fortin et al., 2017]: (1) fractional anisotropy (FA) - mean, standard deviation (SD), and median FA values per ROI (279 features); (2) axial diffusivity (AxD) - mean, SD, and median AxD values per ROI (265 features); (3) mean diffusivity (MD) - mean, SD, and median MD values per ROI (265 features); and (4) radial diffusivity (RD) - mean, SD, and median RD values per ROI (264 features). Note that the numbers of each feature differs slightly as some of the values may have been missing, and variables were removed with many missing entries. DTI features were not available for all participants, as DTI is not routinely collected by all cohorts, leading to the lowest counts among imaging measures (N = 49-220 per outcome; Table 2).

### 2.4 Machine Learning

As noted earlier, a wide range of ML methods could be tested for this task; in prior work, we have assessed deep learning and computer vision methods for related tasks. Due to its strong performance in our prior work predicting cognitive decline [Chattopadhyay et al 2025b; Chattopadhyay et al 2026], we used AutoGluon, a so-called “AutoML” method that searches for the best machine learning model for a given dataset, based on the amount and type of available data. This approach is also publicly available and easily handles tabular data, such as numerical scores.

Overall, we trained separate models for each neuropathological outcome, systematically evaluating different predictor feature sets (combined with base clinical covariates) within a unified cross-validation framework. Predictive models were trained using AutoGluon-Tabular (a version of AutoGluon designed to handle numeric data) with the “medium” quality presets [Erickson et al., 2020]; these are the default configuration and were chosen for their time efficiency - in our experiments, higher quality presets increased training time considerably, without any measurable improvement in predictive performance - as was also noted in the original AutoGluon paper [Erickson et al., 2020; Gijsbers et al., 2024]. AutoGluon trains an ensemble of machine learning model types, including gradient boosting machines, neural networks, and other tabular learners via multi-layer stack ensembling [Wolpert, 1992]. The evaluation metric used to guide model selection and training was the Matthews Correlation Coefficient (MCC) [Matthews, 1975; Chicco & Jurman, 2020], as it is suitable for classification problems with class imbalance. The MCC accounts for all four categories of the confusion matrix (true positives, false negatives, true negatives, and false positives) in proportion to the size of each class.

### 2.5 Cross-Validation

All models were evaluated using stratified 5-fold cross-validation. Stratification was applied jointly by “ΔAge” (the interval between the last antemortem assessment and death) and age at death, to ensure that both variables were proportionally represented across folds [Kohavi, 1995; Varoquaux, 2018]. Predicted class probabilities and predicted labels across all five held-out test folds were pooled to form a single set of out-of-sample predictions for each feature set-outcome combination, over which all performance metrics were subsequently calculated. While we focused on the last antemortem assessment, future work will examine longitudinal changes (e.g., first vs. last assessments), which may provide additional signal for modalities such as T1 and FLAIR.

### 2.6 Performance Metrics

Two correlation-based performance metrics were computed from the pooled cross-validation predictions for each feature set-outcome pairing. Reliable evaluation requires performance metrics that are robust to class imbalance, which is common in autopsy datasets. As noted above, the Matthews Correlation Coefficient (MCC) accounts for all cells of the confusion matrix and remains informative even when outcome classes are highly imbalanced - properties that make it increasingly recommended for biomedical classification problems [Gorodkin, 2004; Grandini et al., 2020; Chicco & Jurman 2020; Itaya et al., 2025].

#### 2.6.1 Matthews Correlation Coefficient (MCC)

MCC was the primary metric reported for binary outcomes. MCC confidence intervals and p-values were estimated using Fisher’s z-transform and the delta method, following the asymptotic framework introduced by Itaya et al. [2025].

#### 2.6.2 Spearman’s Rank Correlation Coefficient (Rs)

Rs was the primary metric for ordinal multi-class outcomes. It measures the degree to which predicted rank order agrees with true rank order [Spearman, 1904]. Rs confidence intervals and p-values were estimated by computing Pearson’s correlation coefficient on the mean rank values assigned to tied observations, with significance assessed via the standard Student’s t-distribution approximation [Conover 1999].

Both MCC and Rs were calculated for all feature set-outcome pairings and are detailed in the supplementary results tables. The supplementary results table also includes balanced accuracy [Broderson et al., 2010] and F1-Score [Grandini et al., 2020] for comparability with prior research.

P-values were adjusted for multiple comparisons using the Benjamini-Hochberg false discovery rate (FDR) procedure [Benjamini & Hochdorf, 1995], applied across all outcome-feature set combinations. Significance thresholds for adjusted p-values are denoted as follows: * padj < 0.05, ** padj < 0.01, *** padj < 0.001, **** padj < 0.0001. This asterisk notation is used in both the tables and graphs of results, to make them easier to interpret.

#### 2.6.3 Incremental Prediction Over Base Clinical Features

For every feature set except the “base clinical” data, an incremental analysis was conducted to determine whether inclusion of that feature set statistically improved prediction over the base clinical feature set alone. This analysis was restricted to the subset of participants for whom predictions were available under both the given feature set and base clinical data, ensuring a fair comparison.

For ΔMCC, the asymptotic variance of MCC was used to test whether MCC□ (the given feature set) was greater than MCC□ (base clinical data) via a one-tailed dependent difference test using the modified transform method described by Itaya et al. [2025]. The hypothesis tested was that MCC□ − MCC□ > 0.

For Rs, the dependent difference between two Spearman correlations was assessed using a bootstrapped procedure with 10,000 replicates. The p-value was estimated as the proportion of bootstrap samples in which Rs1 − Rs2 > 0, yielding a one-tailed test of whether the given feature set produced a higher rank correlation than the base clinical data alone for the same participants.

### 2.7 Class-Specific Ordinal Prediction Assessments

For each ordinal K-class outcome (where, e.g., A<B<C<D<E), we binarized the prediction and truth variable via binary decomposition into K-1 boundaries where, e.g. A|BCDE, AB|CDE, ABC|DE, and ABCD|E [Frank & Hall, 2001]. For binary outcomes, we report MCC, which summarizes confusion matrix performance and is robust to class imbalance; Spearman’s rank correlation was used for ordinal outcomes. As in 2.6.1, we calculated statistical significance (Ha: MCC != 0) for each FeatureSet:OutcomeClass pairing, using the z-transformed delta method and the asymptotic framework from Itaya et al. [2025], and adjusted for multiple comparisons [Benjamini & Hochdorf, 1995]. We also assessed whether the FeatureSet::OutcomeClass pairing improved the prediction over the prediction of the Clinical::OutcomeClass and reported the magnitude of that improvement and its significance (Ha: ΔMCC > 0; again adjusting for multiple comparisons [using the Benjamini & Hochberg method]; following the same procedures in 2.6.3). The results and discussion are summarized in the supplementary materials.

## 3. Results

To orient the reader to the results, we present below the findings for each major neuropathologic outcome, detailing which sets of features performed best in predicting each. After that, we examine the predictor sets in turn, identifying the predictive tasks for which each predictor added value. The text below guides the reader through a series of graphs. Each graph is organized in the same way: for a given neuropathologic outcome (e.g., Thal phase or CERAD score), we show the predictive accuracy of a model using baseline clinical data alone, or supplemented with neuroimaging, CSF, cognitive composite scores, and white matter hyperintensity metrics. The bars denote the accuracy of the models using different predictor sets. Higher values denote better performance.

### 3.1 Amyloid Measures (Figure 1, Table 4)

#### 3.1.1 Thal Phase (6-class ordinal, R⍰)

Thal phases (0–5) track the progression of amyloid-β accumulation through the brain, based on post-mortem histopathology. The top three predictive feature sets for Thal phase were CSF biomarkers (R⍰ = 0.666 ****), cognitive composite scores (Rs = 0.506 ****) and base clinical feature sets (R⍰ = 0.497 ****). This is in line with expectation, as elevated beta-amyloid is robustly present many years before symptoms and certainly before atrophy becomes apparent. However, CSF features did not outperform the base clinical data on the much smaller subset of matched participants (N = 169, ns), so ascertainment bias cannot be ruled out. Among the structural MRI metrics, rICV metrics performed best (R⍰ = 0.337 ****), followed by cortical thickness (R⍰ = 0.305 ***) and volumes (R⍰ = 0.209 *). WMH achieved a moderate R⍰ of 0.314 ****. Among DTI metrics, only axial diffusivity reached significance (R⍰ = 0.295 *). We note that diffusion metrics may reflect regionally heterogeneous effects (e.g., increases or decreases depending on fiber architecture such as crossing fibers), which may not be fully captured by global or summary diffusivity measures.

**Figure 1:**
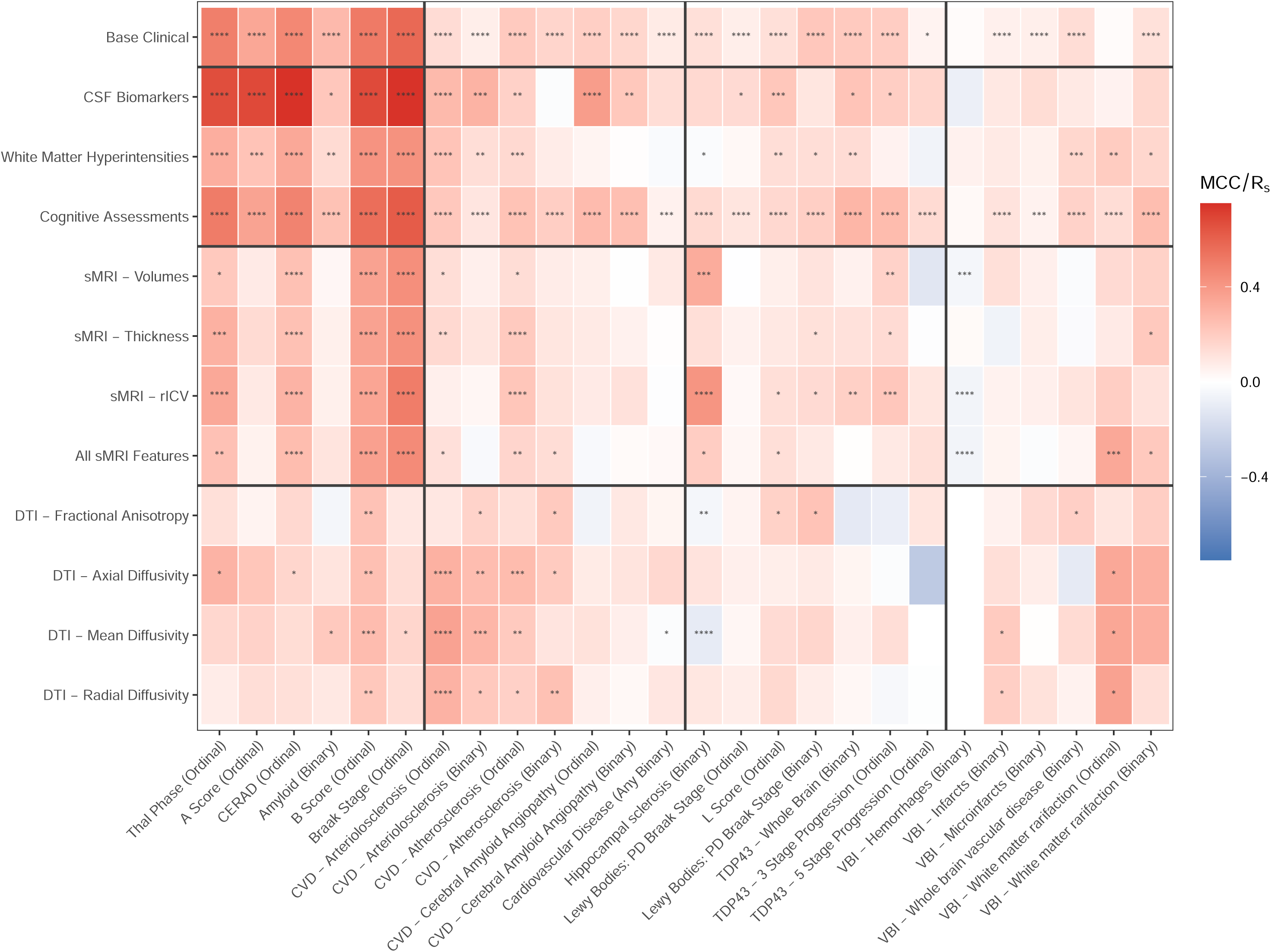
Prediction accuracy for Amyloid and Tau outcomes. Black stars indicate a statistically significant prediction better than random chance (padj != 0). Red stars indicate a statistical improvement of the participants in the given feature set over that of the predictions of the base clinical data alone.

**Table 4:** Amyloid & Tau Measure Prediction. The bolded value represents the best measure (MCC or Rs) for that given outcome. Stars reference the significance that the measure does equal 0 (See Methods). Values in red indicate that the measure for the given feature set gave a statistical improvement over matched Base Clinical predictions (when using the subset of participants from the feature set being studied).

#### 3.1.2 A Score (4-class ordinal, R⍰)

CSF biomarkers were the dominant predictor of A Score (R⍰ = 0.683 ****), significantly improving upon the base clinical predictions (ΔRs = 0.340, N = 169, *). Base clinical data produced a moderate but significant association (R⍰ = 0.343 ****). WMH and cognitive composite scores were the only other significant feature sets (R⍰ = 0.232 *** and 0.3575 ****; respectively). All sMRI- and DTI subtypes were non-significant.

#### 3.1.3 CERAD Score (4-class ordinal, R⍰)

CERAD score was the best-predicted beta-amyloid outcome overall, with CSF biomarkers achieving R⍰ = 0.746 **** - the single highest R⍰ value observed across the entire study - and this was a significant improvement on base clinical data (ΔRs = 0.191, N = 281 *). Base clinical data performed strongly (R⍰ = 0.463 ****), as did cognitive composite scores (R⍰ = 0.479 ****). The cognitive composite scores predictions significantly improved upon the performance of the base clinical (ΔRs = 0.020, N = 7,364 *). All sMRI feature sets reached significance (R⍰ range: 0.242-0.295 ****), and WMH achieved R⍰ = 0.339 ****. DTI axial diffusivity was the DTI metric that best predicted the CERAD score (R⍰ = 0.160 *).

#### 3.1.4 Amyloid Positivity (binary, MCC)

Binary beta-amyloid classification showed a notably different pattern from the ordinal beta-amyloid outcomes. Base clinical data achieved the highest MCC (0.272 ****), with CSF biomarkers yielding a lower MCC despite being the top-ranked predictor for ordinal beta-amyloid outcomes (MCC = 0.217 *). Cognitive composite scores were also significant (MCC = 0.239 ****). Mean diffusivity was the only significant DTI predictor (MCC = 0.210 *). All other sMRI feature sets and remaining DTI metrics were non-significant for binary amyloid classification. WMH reached marginal significance (MCC = 0.140 *). This pattern suggests that binary beta-amyloid classification, as defined here, was less well predicted by CSF than the ordinal staging schemes. CSF analytes may therefore be more sensitive to the graded pathological burden than to the binary presence/absence threshold.

### 3.2 Pathologic Tau Measures (Figure 1; Table 4)

#### 3.2.1 Braak Stage (7-class ordinal, R⍰)

Braak stage was among the most consistently and strongly predicted outcomes across all feature sets. CSF biomarkers led with R⍰ = 0.742 ****, showing a significant incremental improvement over base clinical (ΔRs = 0.159, N = 280; *). Cognitive composite scores were the second-strongest predictor (R⍰ = 0.620 ****), also improving on the base clinical features (ΔRs = 0.047, N = 731; ****), which were the third-best predictor (R⍰ = 0.579 ****). All sMRI sub-types reached significance (R⍰ range: 0.429-0.502 ****), with rICV the strongest sMRI predictor (R⍰ = 0.502 ****). WMH achieved R⍰ = 0.424 ****. Of the DTI measures, only mean diffusivity reached significance (R⍰ = 0.161 *); FA, AD, and RD were non-significant.

#### 3.2.2 B Score (4-class ordinal, R⍰)

B Score was also well predicted across feature sets, with CSF biomarkers leading (R⍰ = 0.678 ****). Base clinical (R⍰ = 0.515 ****), cognitive composite scores (R⍰ = 0.559 ****), and all sMRI features (R⍰ range: 0.350–0.370 ****) showed significant associations. WMH achieved R⍰ = 0.419 ****. Importantly, B Score was the best-predicted outcome for DTI features: all four DTI metrics reached significance (FA R⍰ = 0.236 **, AD R⍰ = 0.246 **, MD R⍰ = 0.259 ***, RD R⍰ = 0.216 **), marking B Score as uniquely recoverable from white matter microstructure. Only the cognitive assessment feature set significantly improved predictions over the base clinical data (ΔRs = 0.0498, N = 7,310 ****).

### 3.3 CVD (Figure 2, Table 5)

#### 3.3.1 Arteriolosclerosis (4-class ordinal, R⍰)

DTI diffusivity metrics were the strongest predictors of ordinal arteriolosclerosis. MD achieved the highest R⍰ across all feature sets for this outcome (R⍰ = 0.360 ****), followed by AD (R⍰ = 0.302 ****) and RD (R⍰ = 0.298 ****). FA was non-significant. CSF biomarkers also performed well (R⍰ = 0.269 ****). WMH showed a significant association (R⍰ = 0.228 ****). Predictions based on cognitive composite scores were also significant (R⍰ = 0.214 ****), and improved over the base clinical model on matched participants (ΔRs = 0.081, N = 6,818; ****). Base clinical prediction was significant but modest (R⍰ = 0.135 ****), as was sMRI thickness (R⍰ = 0.147 **), volumes (R⍰ = 0.127 *), and “all sMRI” (R⍰ = 0.119 *). rICV was non-significant.

**Figure 2:**
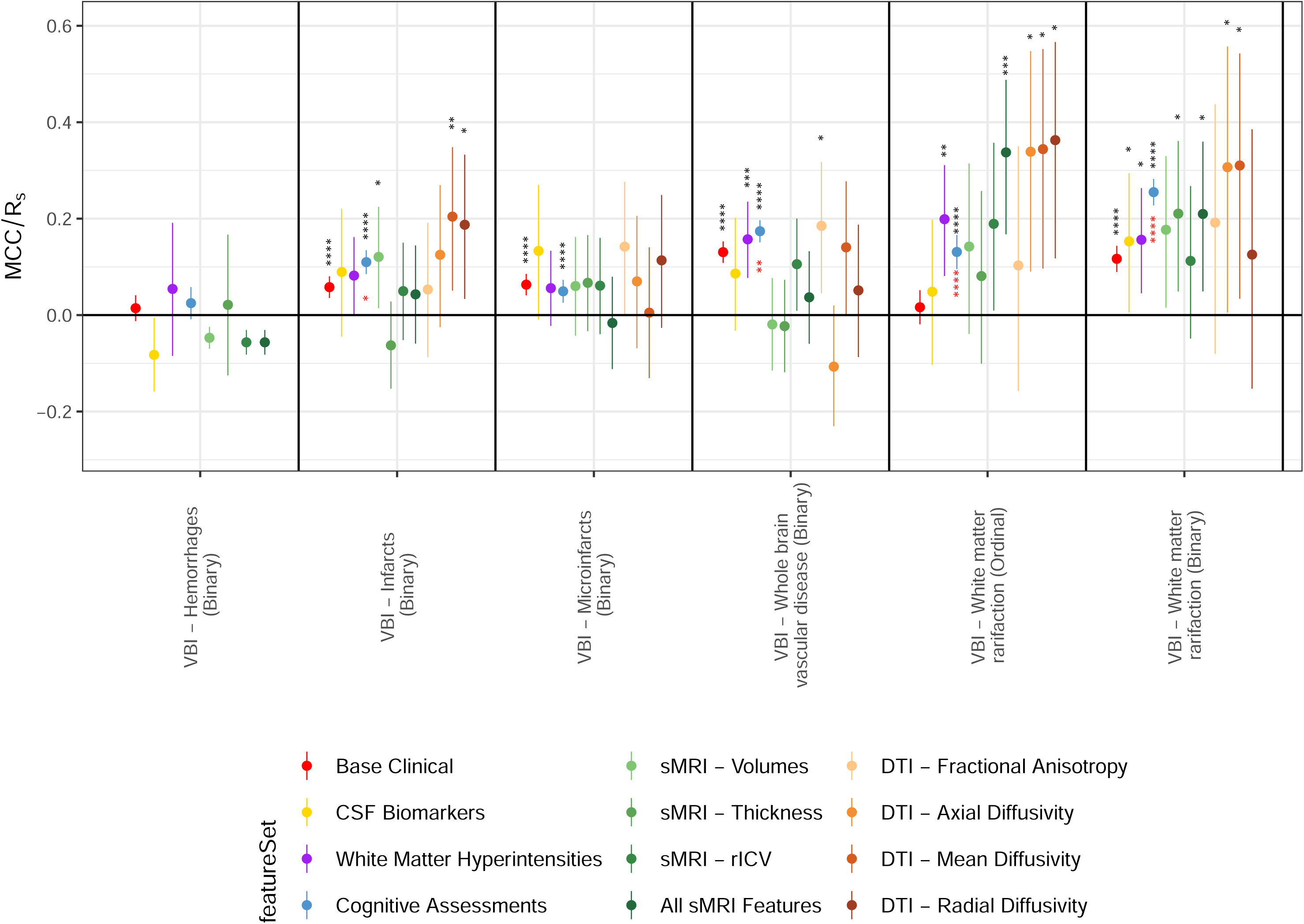
Prediction Accuracy for Cardiovascular Disease Measures. Black stars indicate a statistically significant prediction better than random chance (padj != 0). Red stars indicate a statistical improvement of the participants in the given feature set over that of the predictions of the base clinical data alone.

**Table 5:** Vascular Disease Measure Predictions. Amyloid & Tau Measure Prediction. The bolded value represents the best measure (MCC or Rs) for that given outcome. Stars indicate the significance level at which the measure differs from zero (see Methods). Values in red indicate that the measure for the given feature set yielded a statistical improvement over matched Base Clinical predictions (when using the subset of participants from the feature set being studied).

#### 3.3.2 Arteriolosclerosis (binary, MCC)

For binary classification, CSF biomarkers were the top predictor (MCC = 0.296 ***) with a statistically significant incremental improvement over the base clinical model (ΔMCC = 0.307, N = 281, *). DTI diffusivity metrics remained strongly predictive (MD MCC = 0.284 ***, AD MCC = 0.255 ***, RD MCC = 0.211 *), and DTI FA reached significance (MCC = 0.172 *). Base clinical was modestly significant (MCC = 0.067 ****). Cognitive composite scores were significant (MCC = 0.099 ****). All sMRI subtypes were non-significant.

#### 3.3.3 Atherosclerosis (4-class ordinal, R⍰)

DTI axial diffusivity was the strongest predictor of ordinal atherosclerosis (R⍰ = 0.262 ***), comparable to or exceeding sMRI thickness (R⍰ = 0.207 ****) and rICV (R⍰ = 0.223 ****). Base clinical data achieved R⍰ = 0.206 ****. MD (R⍰ = 0.208 **) and RD (R⍰ = 0.181 *) also reached significance. CSF biomarkers were significant but modest (R⍰ = 0.178 **). Cognitive composite scores (R⍰ = 0.2351 ****), WMH (R⍰ = 0.147 ***), sMRI volumes (R⍰ = 0.132 *), and “all-sMRI” (R⍰ = 0.162 **) also achieved significance. FA alone was non-significant. The cognitive composite scores feature set alone significantly improved on matched base clinical predictions (ΔRs = 0.036, N = 7,341, *).

#### 3.3.4 Atherosclerosis (binary, MCC)

DTI RD was the strongest single predictor for binary atherosclerosis classification (MCC = 0.243 **). DTI FA (MCC = 0.208 *), AD (MCC = 0.204 *), cognitive composite scores (MCC = 0.191 ****), and base clinical data (MCC = 0.165 ****) reached significance; Of the sMRI features, only the composite “all-sMRI” features reached significance (MCC = 0.130 *). CSF biomarkers, DTI MD, and WMH were non-significant.

#### 3.3.5 CAA (4-class ordinal, R⍰)

CSF biomarkers were by far the strongest predictor for ordinal CAA staging (R⍰ = 0.382 ****). Cognitive composite scores achieved the second highest R⍰ (0.264 ****) and improved over matched base clinical predictions (ΔRs = 0.078, N = 7,260; ****). The base clinical model was significant but modest (R⍰ = 0.186 ****). All sMRI features, WMH, and all DTI sub-types were non-significant for ordinal CAA, highlighting the specificity of CSF amyloid-tau analytes to this outcome relative to structural or diffusion imaging.

#### 3.3.6 CAA (binary, MCC)

Binary CAA was uniquely distinguished by strong performance from cognitive composite scores (MCC = 0.249 ****), improving on the base clinical predictions (ΔMCC = 0.096, N = 7,260 ****). CSF biomarkers (MCC = 0.216 **) and base clinical data (MCC = 0.156 ****) were also significant. sMRI rICV was marginal (MCC = 0.115, ns). Most imaging feature sets were non-significant.

#### 3.3.7 CVD — Any (binary, MCC)

The binary omnibus CVD indicator was the most difficult CVD outcome to predict, partly due to its extremely low class balance (truthBalance = 0.222). Base clinical and cognitive composite scores were the only feature sets reaching significance (MCC = 0.079 ****; MCC = 0.055 ****, respectively). All other feature sets, including CSF, sMRI sub-types, DTI sub-types, and WMH, were non-significant. This outcome likely captures only the most severe, clinically surfaced CVD cases and may carry little discriminative signal beyond age and diagnosis.

### 3.4 Neurodegeneration (Figure 3; Table 6)

#### 3.4.1 Hippocampal Sclerosis (binary, MCC)

Hippocampal sclerosis was uniquely well-predicted by volumetric sMRI, with rICV achieving the highest MCC among sMRI predictors across all outcomes (MCC = 0.413 ****) and a statistically significant incremental improvement over base clinical matched predictions (ΔMCC = 0.363, N = 301 *). sMRI volumes also performed strongly (MCC = 0.324 ***), as expected, hippocampal volume is almost a direct measure of hippocampal sclerosis. Base clinical data was relatively weak (MCC = 0.120 ****), and cognitive composite scores were modest (MCC = 0.144 ****), without significant incremental gain over base clinical data. CSF biomarkers were non-significant (MCC = 0.149, ns). Several DTI feature sets yielded small but statistically significant negative MCC values (DTI-MD: MCC =-0.100 ****, DTI-FA: MCC =-0.046 **), indicating that these features did not contain a stable disease-related signal for hippocampal sclerosis. In cross-validation, the learned decision boundaries were weak and inconsistent across folds, resulting in predictions that were slightly anti-correlated with the true labels. This pattern suggests that hippocampal sclerosis is primarily captured by gray-matter structural measures rather than white-matter microstructure metrics from DTI.

**Figure 3:**
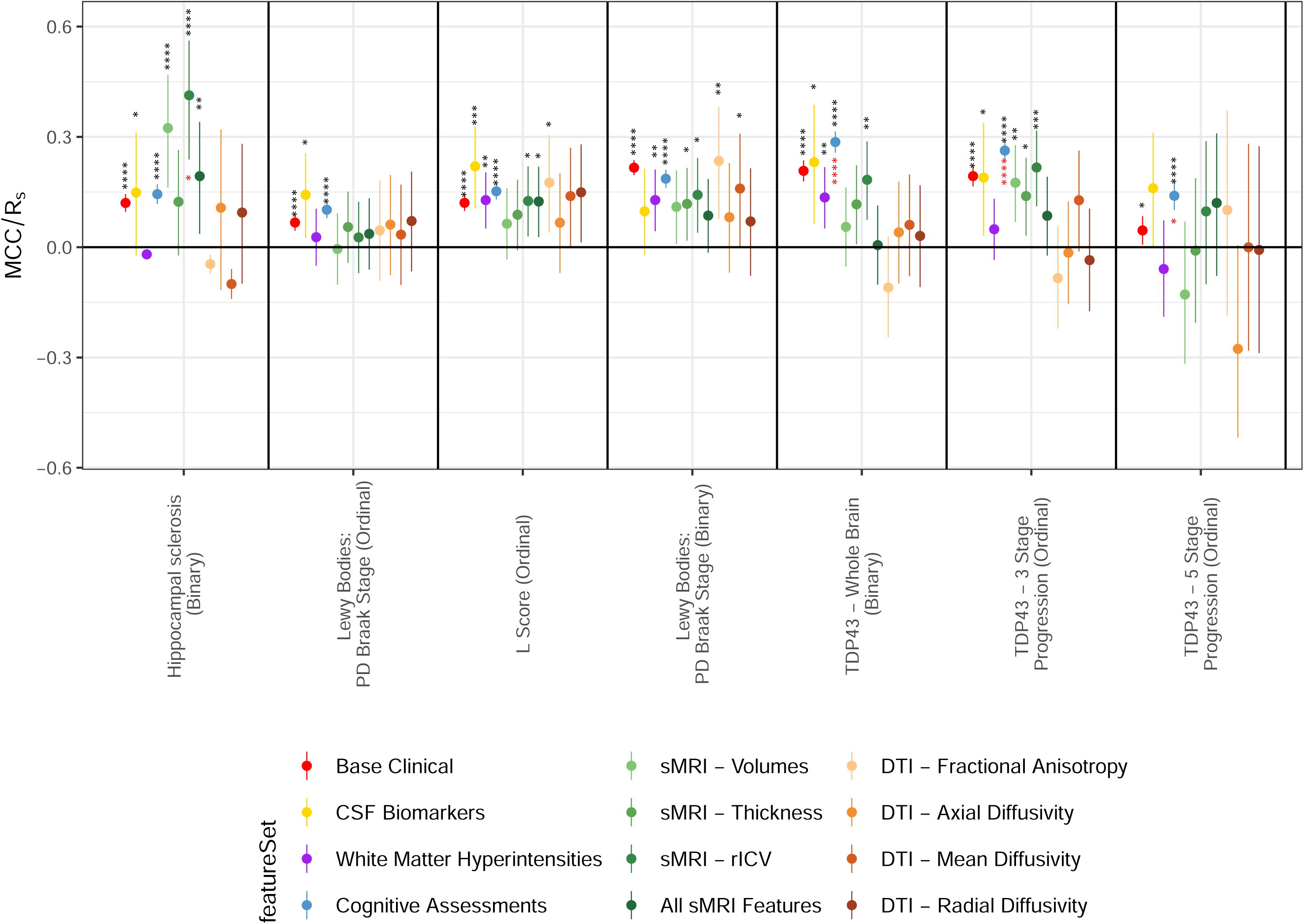
Prediction Accuracy for Hippocampal Sclerosis, Lewy bodies and TDP43. Black stars indicate a statistically significant prediction better than random chance (padj != 0). Red stars indicate a statistical improvement of the participants in the given feature set over that of the predictions of the base clinical data alone.

**Table 6:** Neurodegenerative, Lewy Bodies and TDP-43 outcomes. The bolded value represents the best measure (MCC or Rs) for that given outcome. Stars indicate the significance level at which the measure differs from zero (see Methods). Values in red indicate that the measure for the given feature set gave a statistical improvement over matched Base Clinical predictions (when using the subset of participants from the feature set being studied).

### 3.5 LBD (Figure 3; Table 6)

#### 3.5.1 PD Braak Stage (5-class ordinal, R⍰)

Ordinal PD Braak staging was poorly predicted across all feature sets. CSF biomarkers achieved the highest R⍰ (0.142 *), followed by cognitive composite scores (R⍰ = 0.102 ****) and Base Clinical (R⍰ = 0.067 ****). All sMRI subtypes, WMH, and DTI subtypes were non-significant. The very low overall R⍰ values indicate that fine-grained 5-class LBD staging is largely unpredictable from any of the antemortem feature sets studied here.

#### 3.5.2 L Score (3-class ordinal, R⍰)

The 3-class L Score showed broader, modestly significant associations. CSF biomarkers led (R⍰ = 0.220 ***), followed by cognitive composite scores (Rs = 0.152 ****), WMH (R⍰ = 0.128 **), and sMRI rICV (R⍰ = 0.126 *), All sMRI Features (R⍰ = 0.124 *), the base clinical model (R⍰ = 0.121 ****), and DTI FA reached significance (R⍰ = 0.176 *)-the only DTI metric to do so for this outcome.

#### 3.5.3 LBD Presence (binary, MCC)

Binary LBD classification was most strongly predicted by DTI FA (MCC = 0.235 *), with base clinical (MCC = 0.216 ****), and cognitive composite scores also significant (MCC = 0.186 ****). Among imaging feature sets, DTI FA was the strongest (MCC = 0.235 *), followed by sMRI rICV (MCC = 0.142 *), thickness (MCC = 0.118 *), and WMH (MCC = 0.128 *). CSF biomarkers were non-significant (MCC = 0.098, ns). The binary classification was substantially better predicted than the 5-class ordinal PD Braak stage across all feature sets, despite the expectation of greater precision from fine-grained LBD staging systems.

### 3.6 Pathologic TDP-43 (Figure 3; Table 6)

#### 3.6.1 Pathologic TDP-43 Whole Brain Binary (binary, MCC)

Cognitive composite scores were the strongest predictor of binary pathologic TDP-43 presence (MCC = 0.286 ****), narrowly exceeding CSF biomarkers (MCC = 0.231 *) and Base Clinical (MCC = 0.208 ****). sMRI rICV was also significant (MCC = 0.183 **), as was WMH (MCC = 0.135 **). All DTI subtypes were non-significant. Cognitive composite scores significantly improved upon matched base clinical predictions (ΔMCC = 0.079, N = 4,504 ***). The uniquely strong performance of cognitive composite scores - not replicated by any imaging feature set - is consistent with evidence that neuropathologic changes of limbic-predominant age-related TDP-43 encephalopathy (LATE-NC) leads to a neuropsychological signature detectable from domain-specific cognitive profiles.

#### 3.6.2 Pathologic TDP-43 3 Stage (4-class ordinal, R⍰)

Cognitive composite scores were again the top predictor of pathologic TDP-43 staging (R⍰ = 0.263 ****), a significant incremental improvement over the base clinical model (ΔRs = 0.070, N = 4,504 ****). Base clinical achieved R⍰ = 0.193 ****, with sMRI rICV (R⍰ = 0.217 ***), volumes (R⍰ = 0.175 **) and thickness (R⍰ = 0.139 *) all reaching significance. CSF biomarkers were marginally significant (R⍰ = 0.189 *). WMH and all DTI subtypes were non-significant. The pattern of cognitive and sMRI associations for pathologic TDP-43 staging, in the absence of meaningful DTI signal, distinguishes this outcome from CVD and WM outcomes, where DTI features are informative.

#### 3.6.3 Pathologic TDP-43 5 Stage (6-class ordinal, R⍰)

The 6-class pathologic TDP-5 staging outcome showed only significant associations with cognitive composite scores (R⍰ = 0.1398 ****), which improved on matched base clinical predictions (ΔRs = 0.094; N = 2,460 *). The base clinical model reached only marginal significance (R⍰ = 0.046 *). This near-universal failure reflects both the high granularity of the 6-class staging scheme and the small subsample sizes available for this outcome (N = 49–223 across most feature sets), which hamper the detection of any signal.

### 3.7 VBI (Figure 4; Table 7)

#### 3.7.1 Hemorrhages (binary, MCC)

Hemorrhages was the only outcome to remain entirely non-significant across all 12 feature sets tested. The lack of any predictive signal suggests that this pathologic outcome represents a categorically distinct prediction problem: although discrete hemorrhagic lesions may have profound clinical and neuroimaging consequences, they leave no consistent antemortem structural, microstructural, or biofluid fingerprint detectable from the feature sets evaluated here.

**Figure 4:**
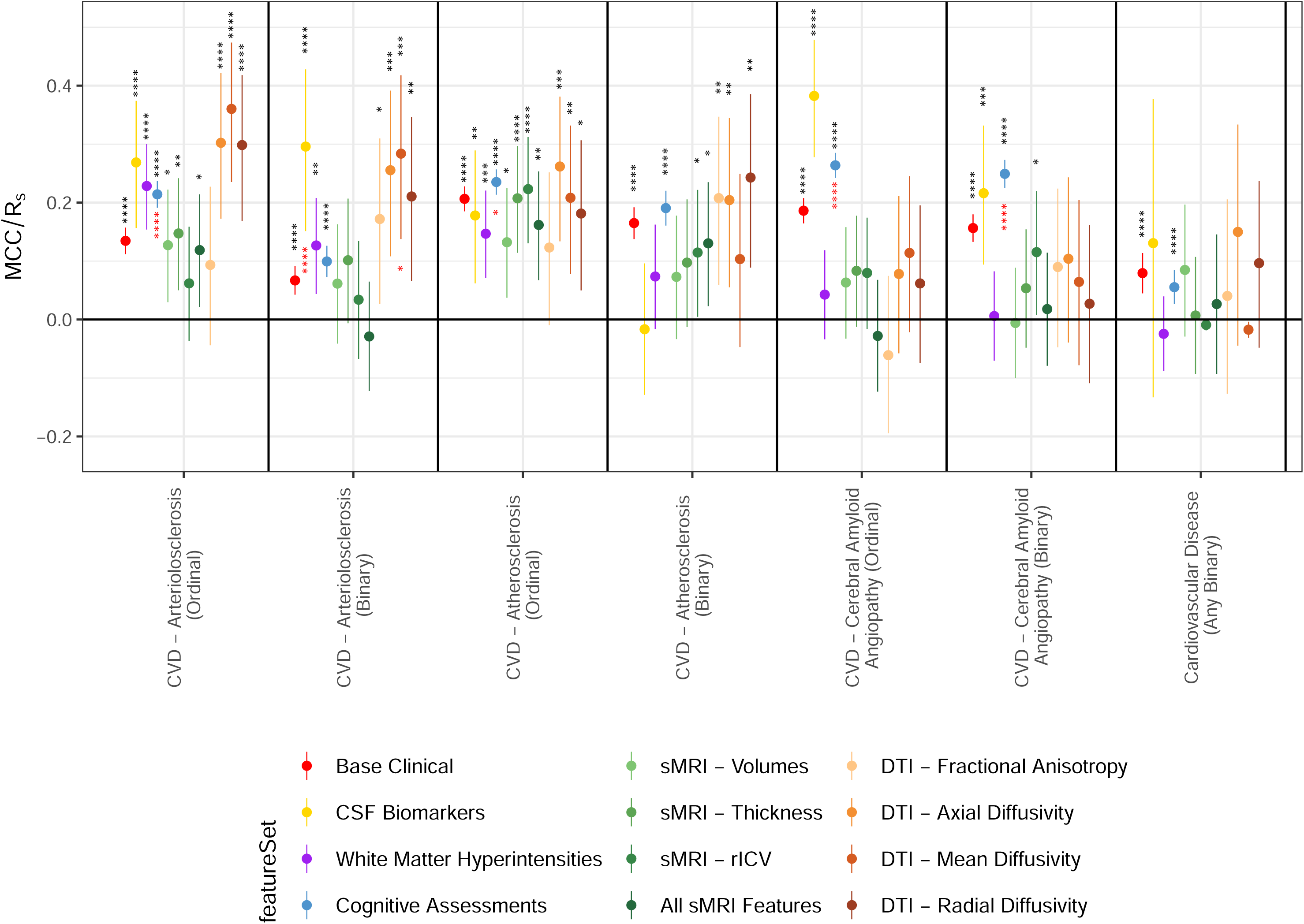
Prediction Accuracy for Vascular Brain Injury metrics. Black stars indicate a statistically significant prediction better than random chance (padj != 0). Red stars indicate a statistical improvement of the participants in the given feature set over that of the predictions of the base clinical data alone.

**Figure 5:**
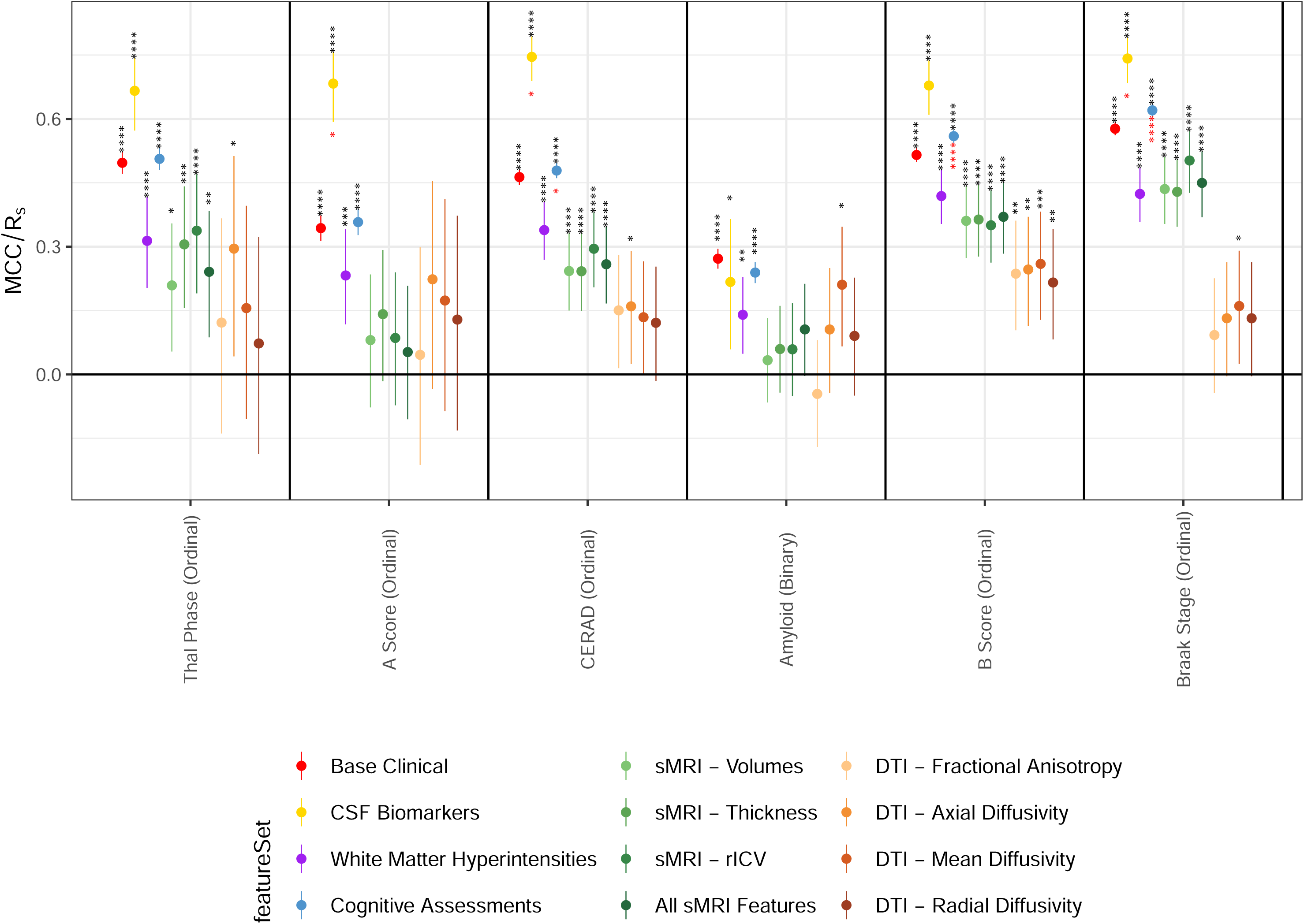
Predictor Performance of Neuropathology Classification. Heatmap showing the pooled 5-fold cross-validation prediction performance using the AutoGluon ‘Medium” presets. Warmer colors (red) indicate stronger predictive performance (higher MCC/Rs), highlighting the biomarker domains most informative for neuropathology prediction, while cooler colors (blue) indicate weaker or inverse predictive associations. Rows correspond to the Feature Sets described in this paper. Black stars indicate a statistically significant prediction better than random chance (padj != 0)

**Table 7:** Vascular Brain Injury Measures. The bolded value represents the best measure (MCC or Rs) for that given outcome. Stars indicate the significance level at which the measure differs from zero (see Methods). Values in red indicate that the measure for the given feature set gave a statistical improvement over matched Base Clinical predictions (when using the subset of participants from the feature set being studied).

#### 3.7.2 Infarcts (binary, MCC)

While strategically located brain infarcts may severely impact different cognitive domains, including memory, this global metric of infarcts showed only small but significant associations in several feature sets. DTI mean diffusivity was the strongest predictor (MCC = 0.204 **), followed by DTI radial diffusivity (MCC = 0.187 *). Base Clinical (MCC = 0.058 ****) and cognitive composite scores (MCC = 0.110 ****) were significant but modest. Cognitive composite scores improved on their respective matched base clinical predictions (ΔMCC = 0.050, N = 7,332 *). sMRI Volumes (MCC = 0.121 *) also reached significance. CSF biomarkers, WMH, DTI FA, and DTI AD were non-significant.

#### 3.7.3 Microinfarcts (binary, MCC)

Microinfarcts showed similarly small but significant associations. Base clinical data was significant (MCC = 0.063 ****), as were cognitive composite scores (MCC = 0.049 ***). DTI FA (MCC = 0.142, ns) and DTI RD (MCC = 0.114, ns) did not survive FDR correction. CSF biomarkers, WMH, and most sMRI subtypes were non-significant. The low MCC values throughout reflect the difficulty of detecting diffuse microscopic lesions from macro-level imaging features.

#### 3.7.4 Whole Brain Vascular Disease (binary, MCC)

Whole brain vascular disease (binary) showed small but significant associations with cognitive composite scores (MCC = 0.174 ****), Base clinical (MCC = 0.131 ****), WMH (MCC = 0.157 ***), and DTI FA (MCC = 0.185 *). CSF biomarkers, most sMRI subtypes, and other DTI subtypes were non-significant.

#### 3.7.5 White Matter Rarefaction (4-class ordinal, R⍰)

White matter rarefaction (4-class) was not significantly predicted by Base clinical data (R⍰ = 0.016, ns), in contrast to most other outcomes. DTI diffusivity metrics were the strongest predictors: DTI RD (R⍰ = 0.363 **) and AD (R⍰ = 0.339 *) both reached significance, as did “All sMRI” Features (R⍰ = 0.337 ***) and WMH (R⍰ = 0.199 **). Cognitive composite scores were the most significant predictor (R⍰ = 0.131 ****) and significantly improved on matched base clinical predictions (ΔRs = 0.111, N = 2,967 ****). CSF biomarkers were non-significant (R⍰ = 0.049, ns). The clear benefit of neuroimaging in this case aligns with the known sensitivity of white matter microstructure and WMH measures to the perilesional changes associated with white matter rarefaction.

#### 3.7.6 White Matter Rarefaction (binary, MCC)

For the binary version of this outcome, base clinical data reached significance (MCC = 0.117 ****). Cognitive composite scores had the largest significant MCC (MCC = 0.255 ****), which also improved on matched base clinical predictions (ΔMCC = 0.154, N = 4,950 ****). Cortical thickness (MCC = 0.210 *) and “All sMRI” features (MCC = 0.210 *) also reached significance, as did WMH (MCC = 0.156 *).

### 3.8 Results by Feature Set (See Figfure 5)

#### 3.8.1 Base Clinical

With only four features (sex, age at death, time since last assessment, and cognitive status), the base clinical data achieved significant associations for 24 of 26 outcomes, the second widest coverage of any feature set. Years of education was included with the cognitive composite features as a covariate, reflecting its role as a confounder of cognitive performance rather than a standalone clinical predictor. The two non-significant outcomes were Hemorrhages and White Matter Rarefaction (4-class ordinal). The strongest associations were for Braak stage (R⍰ = 0.577 ****), B Score (R⍰ = 0.515 ****), Thal phase (R⍰ = 0.497 ****), CERAD (R⍰ = 0.463 ****), and LBD binary (MCC = 0.216 ****). The base clinical dataset served as the reference benchmark against which all other feature sets were evaluated. Its large sample sizes (N = 3,345-7,865 per outcome) and consistently broad coverage reflect the shared demographic and diagnostic correlates of late-life neuropathology.

#### 3.8.2 CSF Biomarkers

CSF biomarkers yielded the highest absolute metric values observed across the study for dementia-related outcomes: CERAD (R⍰ = 0.746 ****), Braak stage (R⍰ = 0.742 ****), A Score (R⍰ = 0.683 ****), B Score (R⍰ = 0.678 ****), and Thal phase (R⍰ = 0.666 ****). CSF also provided significant incremental improvement over base clinical data for A Score, CERAD, AD Braak Stage, and binary arteriolosclerosis. Performance dropped sharply for LBD outcomes (binary MCC = 0.098, ns), pathologic TDP-43 (binary MCC = 0.231 *), and most VBI outcomes. These results confirm the high specificity of CSF amyloid and tau analytes for AD neuropathological staging, while quantifying the diagnostic gap for non-AD proteinopathies. In four of 26 outcomes, the inclusion of the CSF biomarker data significantly improved the predictive accuracy over the base clinical features alone in matched participants. The sample sizes for CSF biomarkers were much smaller (N = 39-281 per outcome) than for base clinical data, which limits the power to detect incremental improvements. As CSF data were available in a relatively small subset, the sample may be subject to selection bias, as individuals who undergo lumbar puncture may not be representative of the broader cohort; this should be considered when interpreting the strong performance of CSF-based models.

#### 3.8.3 Cognitive Domain Scores

Broadly, cognitive domain scores were the most successful predictors of outcomes, reaching statistical significance on 25 of 26 outcomes, and significantly improving upon the predictions of the base clinical features in 13 of 26 outcomes. We note that there was a larger sample size available for the cognitive domain score feature set than for the other feature sets examined. Also, the cognitive feature set partially overlaps with base clinical variables (e.g., diagnostic status), which may contribute to its strong performance and should be considered when interpreting incremental gains.

Cognitive domain scores produced broad, significant associations across 25 of 26 outcomes. The only outcome that this feature set was unable to significantly predict was Hemorrhages (R⍰ = 0.0248 ns) (which no other feature set was able to successfully predict either). Braak stage (R⍰ = 0.620 ****), B Score (R⍰ = 0.559 ****), and CERAD (Rs = 0.479 ****) were the findings best predicted by cognitive domain scores, with statistically significant incremental improvements over Base Clinical for both. A distinctive feature of this set was its leading performance for pathologic TDP-43 outcomes: 5 stage (Rs = 0.140 ****), 3 stage (R⍰ = 0.263 ****), and binary (MCC = 0.286 ****) were each the largest significant values of any feature set for those outcomes, with all three outcomes showing a significant incremental improvement over Base Clinical. All CVD outcomes were significantly predicted, with each of the ordinal outcomes in this set also significantly improved upon the base clinical feature predictions. Cognitive domain scores also led binary CAA (MCC = 0.234 ****) with a significant incremental improvement over Base Clinical. Thal Phase (R⍰ = 0.506 ****), A Score (R⍰ = 0.358 ****), and binary beta-amyloid (MCC = 0.239 ****) were also significant. These pathologic TDP-43 and CAA patterns were not replicated by any imaging-only feature set, suggesting that domain-specific neuropsychological profiles detect latent pathological signals not accessible from structural or diffusion MRI alone.

#### 3.8.4 Structural MRI

All sMRI subtypes showed significant associations for pathologic tau and beta-amyloid outcomes; Braak stage was consistently the most strongly-predicted outcome (rICV R⍰ = 0.502 ****). The rICV feature set was generally the strongest sMRI predictor, outperforming raw volumes and thickness for most outcomes and providing the only significant incremental improvement over Base Clinical data among sMRI sub-types - specifically for hippocampal sclerosis (ΔMCC = 0.363, N = 301, *). This was the only outcome for which one of the sMRI features improved predictions over the base clinical feature set. Hippocampal sclerosis was the “standout” sMRI outcome, where rICV achieved the highest MCC of any sMRI prediction across all outcomes (MCC = 0.413 ****), markedly exceeding Base Clinical data (MCC = 0.120 ****). Pathologic TDP-3 staging was also significantly predicted by rICV (R⍰ = 0.217 ***) and Volumes (R⍰ = 0.175 **). sMRI features showed little predictive value for VBI outcomes, CVD omnibus, or fine-grained LBD staging.

#### 3.8.5 White Matter Hyperintensities

WMH features were significant for predicting 15 of 26 neuropathological outcomes, but failed to statistically improve upon the predictions of base clinical features for any of the outcomes. Performance was strongest for pathologic tau outcomes (Braak R⍰ = 0.424 ****, B Score R⍰ = 0.419 ****) and broadly comparable to sMRI features for beta-amyloid outcomes (Thal phase R⍰ = 0.314 ****, CERAD R⍰ = 0.339 ****). This is of interest as WMH is expected to be more strongly associated with CVD and VBI, although there is some evidence of an added value of WMH in predicting pathologic tau and beta-amyloid, which may reflect a common cause. As expected, WMH also showed significant associations for arteriolosclerosis (ordinal R⍰ = 0.228 ****, binary MCC = 0.127 *), Whole Brain CVD (MCC = 0.157 ***), WM Rarefaction binary (MCC = 0.156 *), and WM Rarefaction ordinal (R⍰ = 0.199 **). WMH features were non-significant for CAA (ordinal and binary), TDP-43 outcomes, or hemorrhages.

#### 3.8.6 DTI Feature Sets

DTI features showed a distinctive outcome profile diverging markedly from clinical, CSF, and sMRI patterns. Diffusivity metrics (MD, RD, AD) were the strongest predictors of arteriolosclerosis across all tested feature sets and the strongest imaging predictors of WM rarefaction, in both ordinal and binary forms. Specifically:

DTI – Mean Diffusivity achieved the highest R⍰ for arteriolosclerosis ordinal (R⍰ = 0.360 ****) and was significant for infarcts binary (MCC = 0.204 **) and WM rarefaction (ordinal R⍰ = 0.344 *, binary MCC = 0.310 *). Total significant outcomes: 10.

DTI – Axial Diffusivity was strongest for atherosclerosis ordinal (R⍰ = 0.262 ***) and WM rarefaction ordinal (R⍰ = 0.339 *). Total significant outcomes: 9.

DTI – Radial Diffusivity achieved the highest observed R⍰ among all DTI sub-types for WM rarefaction ordinal (R⍰ = 0.363 *) and atherosclerosis binary (MCC = 0.243 **). Total significant outcomes: 7.

DTI – Fractional Anisotropy was the weakest DTI metric overall (6/25 outcomes significant), with its best predictive value being for binary LBD presence (MCC = 0.235 *). FA was non-significant for all beta-amyloid and most pathologic tau outcomes. Although FA is commonly used in prior studies, our findings are consistent with work showing that diffusivity metrics (e.g., MD, RD) are often more sensitive to microstructural degeneration in aging and vascular pathology.

No DTI metric was significant for predicting beta-amyloid positivity (binary) or A-Score ordinal, and DTI was non-significant for pathologic TDP-43 outcomes. DTI features consistently failed to provide statistically significant incremental improvement over Base Clinical by the dependent difference test for any outcome, perhaps reflecting the small DTI sub-cohort sizes (N = 49-220).

### 3.9 Outcomes Resistant to Prediction

Hemorrhages was the only outcome that was non-significantly predicted across all 12 feature sets (max MCC = 0.054; WMH, ns). Pathologic TDP-43 5 Stage and CVD Any Binary were only significantly predicted by the Base Clinical (R⍰ = 0.046 *, R⍰ = 0.079 ****, respectively) and Cognitive composite scores (R⍰ = 0.140 ****, R⍰ = 0.055 ***, respectively). These outcomes represent a predictive frontier where no current antemortem feature set provides discriminative signals, with the models we assessed. For hemorrhages and the omnibus CVD indicator, the low class balance, spatially discrete lesion distributions, and the lack of molecular surrogates likely explain the near-null performance across all modalities.

## 4. Discussion

Across 26 post-mortem neuropathological outcomes, commonly collected clinical, cognitive, MRI, diffusion MRI, and CSF measures showed synergistic ability to predict underlying pathologic changes during life. As expected, CSF biomarkers were the strongest predictors of AD neuropathologic changes, achieving the highest accuracy for beta-amyloid and pathologic tau staging, confirming their value when diagnostic specificity for AD is required. Structural MRI measures were the best predictors of hippocampal sclerosis and pathologic tau-related atrophy, reinforcing their central role in assessing neurodegeneration. Diffusion MRI provided complementary information for vascular and white matter injury, including arteriolosclerosis and white matter rarefaction, highlighting its value when vascular contributions to cognitive impairment are suspected. Cognitive domain scores were broadly predictive across many outcomes and were especially informative for TDP-43 pathology, which was harder to predict from imaging or biofluids. Together, these findings support a targeted multimodal approach in which CSF biomarkers, structural and diffusion MRI, and cognitive testing provide complementary information on disease mechanisms. More broadly, the results clarify when each modality is most informative, providing guidance for clinicians and researchers designing biomarker panels for diagnosis, prognosis, and clinical trials. We note that differences in predictive performance across outcomes and feature sets are influenced not only by biological signal but also by variation in sample size (N) and data availability. Feature sets such as CSF and DTI were available in smaller subsets, reducing statistical power and perhaps underestimating their true predictive value relative to more widely available modalities. Even so, structural and diffusion MRI can often be acquired within a single imaging session, and their complementary strengths suggest that multimodal MRI may improve prediction without substantial added cost over standard structural MRI.

### 4.1 CSF Biomarkers and AD Neuropathologic Changes

The strongest associations across all feature sets were achieved by CSF biomarkers for AD outcomes. Spearman correlations for CERAD score (R⍰ = 0.746 ****), Braak stage (R⍰ = 0.742 ****), A Score (R□ = 0.683 ****), B Score (R□ = 0.678 ****), and Thal phase (R□ = 0.666 ****) were all moderate to high. CSF produced significant incremental improvements over base clinical data for CERAD (ΔR□ = 0.191; N = 281 *), A Score (ΔR□ = 0.340; N = 169 *), and Braak Stage (ΔR□ = 0.159; N = 280 *). CSF also produced the best predictions across feature sets for binary Arteriosclerosis (MCC = 0.296 ***; ΔMCC = 0.307; N = 281 *). CSF analytes - Aβ42, phosphorylated tau, and total tau - reflect the core molecular pathology of AD with high fidelity [Shaw et al., 2009; Blennow et al., 2010]. Since the 2024 revision of the NIA-AA diagnostic criteria [Jack et al., 2018; Jack et al., 2024], these CSF measures are now central to biological staging of AD [Jack et al., 2024]. Even so, due to the high invasiveness and added cost of lumbar puncture, CSF is not routinely used in clinical practice, even in cases where amyloid PET, APOE genotyping, and MRI are used as part of the diagnostic work-up. The costs for routine use of CSF in the AT(N) research framework are also significant [Mattsson et al., 2017; Jack et al., 2018; Burke et al., 2021]. CSF biomarker associations were far weaker for LBD binary classification and pathologic TDP-43 whole brain binary classification (MCC = 0.231 *), and were not significant for hemorrhages and infarcts. Standard CSF panels dominated by Aβ42, p-tau181, and t-tau have no direct molecular surrogate for pathologic α-synuclein or pathologic TDP-43 inclusions. Alpha-synuclein seed amplification assays (αSyn-SAA) are emerging as promising diagnostics for synucleinopathies, demonstrating high sensitivity (86%) and specificity (92%) [Siderowf et al., 2023; Zheng et al., 2023; Espay et al., 2025], but were not available in our current cohorts.

### 4.2 Structural MRI

Across the sMRI features, associations were broadly significant for pathologic tau and beta-amyloid outcomes but generally modest. The rICV-normalized volumes yielded R□ = 0.502 **** for Braak stage and R□ = 0.350 **** for B Score, outperforming raw volumes and thickness for most outcomes. This may reflect the greater sensitivity of ICV-normalized volumetrics for detecting disease-related atrophy independent of head size variation. For every pathologic tau and beta-amyloid outcome where both CSF and sMRI were tested, CSF outperformed all sMRI metrics - for example, CSF R□ = 0.742 versus rICV R□ = 0.502 for Braak stage. This margin is consistent with the model proposed by Jack and colleagues that positions structural MRI as one of the last biomarkers to show detectable changes in the AD cascade [Jack et al., 2010; Jack et al., 2013].

The strongest sMRI association observed was for hippocampal sclerosis, where sMRI volumes achieved MCC = 0.324 **** and rICV reached MCC = 0.413 **** - the highest MCC of any sMRI predictor across all outcomes - with a significant incremental improvement over Base Clinical (ΔMCC = 0.363; N = 301 *). Regional brain volumes carry differential diagnostic information for hippocampal sclerosis relative to the global atrophy pattern captured by clinical staging alone [Nelson et al., 2011; Neltner et al., 2016; Risacher et al., 2017].

### 4.3 DTI as a Vascular and Axonal Injury Marker

The profile of DTI associations diverged markedly from clinical, CSF, and conventional sMRI. Diffusivity metrics - particularly mean and radial diffusivity - were the strongest imaging predictors of arteriolosclerosis across all tested feature sets. DTI-MD achieved R□ = 0.360 **** for the 4-class ordinal scale, and MCC = 0.284 *** for the binary outcome, surpassing Base Clinical (R□ = 0.135 ****) and all sMRI feature sets (all non-significant for binary). For white matter rarefaction, DTI-RD reached the highest R□ of any imaging feature set (R□ = 0.363 *), with DTI-MD (R□ = 0.344 *) and DTI-AD (R□ = 0.339 *) also significant, consistent with the sensitivity of diffusivity to periarteriolar white matter injury underlying these conditions [Pasi et al., 2015; Chen et al., 2023]. Diffusivity metrics are more robust indicators of white matter pathology in aging populations than anisotropy measures, which may be less sensitive to demyelination and axonal loss in the context of heterogeneous white matter lesion distributions [Pasi et al., 2015].

### 4.4 Cognitive Domain Scores

For AD-related outcomes, cognitive domain scores achieved strong associations with Braak stage (R□ = 0.620 ****; ΔR□ = 0.047, N = 7310 ****) and B Score (R□ = 0.559 ****; ΔR□ = 0.050, N = 7310 ****), with significant incremental improvements over Base Clinical for both. CERAD score was also strongly predicted (R□ = 0.479 ****) with significant incremental gain over base clinical (ΔR□ = 0.020; N = 7364 ****). These results align with clinical research linking neurofibrillary and amyloid burden to domain-specific cognitive decline [Jack et al., 2013; Hohman et al., 2023; Mukherjee et al., 2023; Kang et al., 2025].

Cognitive composite scores significantly predicted all seven CVD outcomes, including atherosclerosis binary (MCC = 0.191 ****), atherosclerosis ordinal (R□ = 0.235 ****), and the omnibus CVD binary indicator (MCC = 0.055 ***). Cognitive composite scores significantly improve upon matched Base Clinical predictions for both atherosclerosis ordinal (R□ = 0.235; ΔR□ = 0.036, N = 7341 *) and CAA ordinal (R□ = 0.264 ****, ΔR□ = 0.078; N = 7260 ****) and binary (MCC = 0.249 ****, ΔMCC = 0.096; N = 7260 ****). This pattern is consistent with growing epidemiological and neuropsychological evidence that vascular cognitive impairment - even from subclinical large-vessel atherosclerosis and small-vessel arteriolar disease - produces a characteristic cognitive profile. This may include executive dysfunction and processing speed slowing that precedes clinical diagnosis [Gorelick et al., 2011; Sorond et al., 2015; Ihle et al., 2017]. The CAA results are also notable: the cognitive profile of CAA often includes an amnestic presentation alongside attentional and executive deficits linked to cortical microinfarcts and white matter injury. The strong performance of cognitive composite scores for binary CAA (MCC = 0.249 ****) may reflect this distinctive neuropsychological effect [Charidimou et al., 2017; Greenberg et al., 2018; Charidimou et al., 2022].

Neuropsychological profiles may also help for fine-grained staging of LATE-NC. Recent longitudinal work shows that domain-specific cognitive decline trajectories - particularly in episodic memory and language - diverge in individuals with LATE-NC years before autopsy. These trajectories are partly dissociable from concurrent AD [Boyle et al., 2017; Hiya et al., 2024; Sajjadi et al., 2025; Butler-Pagnotti et al., 2023]. Pathologic TDP-43 proteinopathy lacks validated in vivo biomarkers analogous to those for beta-amyloid and pathologic tau [Nelson et al., 2019; Wolk et al., 2025], making cognitive profiling a valuable interim surrogate for LATE-NC risk stratification before targeted molecular assays are developed and widely available.

For LBD outcomes, cognitive composite scores show significant associations across all three staging schemes: binary LBD presence (MCC = 0.186 ****), L Score (R□ = 0.152 ****), and PD Braak Staging (R□ = 0.102 ****). LBD produces a characteristic neuropsychological profile - featuring visuospatial deficits, attentional fluctuations, and relative preservation of memory relative to AD - that may be partially captured by the harmonized cognitive composites used here [McKeith et al., 2017; Ferman et al., 2013]. Even so, the absence of a visuospatial domain in the current feature set (excluded due to missing data in NACC) may have suppressed performance for these outcomes. These results motivate continued investment in the harmonization of cognitive composite scores across AD/ADRD cohorts [Kang et al., 2025; Mukherjee et al., 2023], including the computation of domain-specific decline measures that may further improve the sensitivity of cognitive data for inferring neuropathologic features.

### 4.5 LBD Staging

Binary LBD classification (Base Clinical MCC = 0.216 ****; Cognitive composite scores MCC = 0.186 ****; DTI-FA MCC = 0.235 * as best imaging predictor) was substantially better predicted than the 5-class ordinal PD Braak stage (R□ = 0.067 **** for Base Clinical; R□ = 0.142 * for CSF as best predictor), and this gap persisted across all feature sets. All sMRI sub-types, WMH, and most DTI sub-types were non-significant for ordinal PD Braak staging. This mirrors a broader challenge in the field: the multiple competing staging systems for LBD show only moderate inter-rater reliability and are not always congruent in categorizing the same cases. The BrainNet Europe consortium found that applying the Braak protocol to α-synuclein pathology yielded inter-observer agreement of only 65% [Alafuzoff et al., 2009], and a subsequent multi-center consensus effort demonstrated that Braak and Beach staging systems showed lower inter-rater reliability (Krippendorff’s α ≈ 0.4) than newly proposed LBD consensus criteria (α ≈ 0.6) [Beach et al., 2009; Attems et al., 2021].

### 4.6 Implications for the Field

CSF biomarkers remain the gold standard in vivo proxy for AD molecular pathology [Jack et al., 2024], with R□ values for CERAD and Braak stage (0.746 and 0.742, respectively) that exceed all imaging-based feature sets; that DTI microstructural imaging provides unique signal for vascular and white matter pathology - outperforming Base Clinical and all sMRI sub-types for arteriolosclerosis ordinal (DTI-MD R□ = 0.360 ****) and white matter rarefaction (DTI-RD R□ = 0.363 *), Cognitive domain scores currently provide the strongest antemortem signal for pathologic TDP-43 outcomes (binary MCC = 0.286 ****), the broadest coverage across CVD outcomes (now significant for all seven, including the omnibus binary indicator at MCC = 0.055 ***), the strongest single predictor for binary white matter rarefaction (MCC = 0.255 ****), and meaningful associations with LBD staging at all levels of granularity. None of these patterns were replicated by any imaging-only or CSF feature set, underscoring the value of neuropsychological profiling in capturing the cognitive sequelae of diseases that currently lack in vivo molecular biomarkers.

From a translational standpoint, the validation of αSyn-SAA and pathologic TDP-43-specific biomarkers remains the highest-yield priority for expanding in vivo neuropathological prediction beyond the AD domain [Nelson et al., 2019; Irwin et al., 2023; Bauer et al., 2023; Espay et al., 2025]. The clear value of DTI measures for CVD supports their broader inclusion in studies of mixed-pathology dementia, where vascular contributions are routinely under-ascertained [Pasi et al., 2015]. Longitudinal domain-specific neuropsychological testing may substantially improve prediction of pathologic TDP-43 staging, LBD presence, and vascular injury outcomes, and should be a priority for future modeling work [Wilson et al., 2020; Dodge et al., 2025; Wolk et al., 2025]. Fourth, the visuospatial cognitive domain - excluded here due to missing data in the NACC sub-cohort - should be prioritized for harmonization in future PHC releases, as it is among the most diagnostically sensitive features for LBD [McKeith et al., 2005; Ferman et al., 2006; McKeith et al., 2017].

### 4.7 Limitations

Several limitations should be considered when interpreting these findings. One fundamental limitation is sample size. Paired ante-mortem imaging and post-mortem neuropathology datasets remain rare, and the ADSP-PHC cohort, while among the largest harmonized resources of this kind, yields subgroups with fewer than 50 participants for some DTI-outcome pairings to several hundred for structural MRI outcomes. The limited training data imposes practical constraints on model capacity and statistical power, particularly for detecting incremental improvements over base clinical features in smaller sub-cohorts. Notably, the neuropathology data from the ADNI cohort has not yet been harmonized with the rest of the PHC neuropathology dataset. Its integration should boost sample sizes in future releases.

Absence of key biomarkers. Several biomarker types that carry strong neuropathological signals were not available in sufficient numbers to be included in the current analysis. Amyloid and pathologic tau PET imaging did not sufficiently overlap with neuropathology outcomes in the current PHC release. Similarly, emerging blood-based biomarkers - including plasma phospho tau species and pathologic alpha-synuclein, which have demonstrated high sensitivity and specificity for synucleinopathies [Espay et al., 2025; Zheng et al., 2023] - and proteomics panels were unavailable at meaningful scale. The absence of these modalities likely explains part of the performance gap observed for ADRD proteinopathies, particularly LBD outcomes, where no validated antemortem molecular surrogate was available.

Finally, feature importance is necessary to shed light into the black box that is machine learning, but at the current sample size, the feature importance used in Autogluon was not robust across folds, suggesting that more data is needed to stabilize these estimates [Strobl et al., 2007].

### 4.8 Future Directions

A near-term priority is the inclusion of ADNI neuropathology and corresponding antemortem feature sets into the ADSP-PHC harmonized framework, which will boost sample sizes and enable more powerful tests of incremental prediction. Future work could also take advantage of longitudinal data, with cognitive and imaging trajectories as predictors. This requires careful time-series modeling within the AutoML framework.

A major goal of future work will be to integrate the mutual information across modalities. For example, if CSF indicates beta-amyloid, but MRI atrophy is disproportionate to what is expected for a given level of beta-amyloid pathology, this may indirectly point to the presence of comorbidity. Incorporating profiles across the modalities may convey useful information in the future, but is challenging as many participants lack at least some of the modalities [Suk et al., 2014; Cheerla & Gevaert, 2019; Lyu et al 2025; Brown et al 2025].

The structural MRI and DTI features used in this study are derived summaries from established parcellation pipelines, not the raw imaging data. Replacing volumetric measures with 3D T1-weighted images, DTI scalar maps with diffusion-weighted images, and white matter hyperintensity features with FLAIR images would allow computer vision architectures - including convolutional neural networks and vision transformers - to learn spatial patterns not captured by region-level summaries [Chattopadhyay et al., 2024; Dhinagar et al., 2023]. Similarly, the development of whole-slide image-based machine learning for automated, quantitative neuropathological scoring [Nguyen et al., 2022; Oliveira et al., 2025] may ultimately provide more reliable and granular outcome labels than current expert rating systems.

Clinical translation and validation. The ultimate translational goal of this work is probabilistic inference of neuropathological burden in living patients, to support clinical trial stratification and individualized care. Prospective validation of the models reported here - in cohorts not included in the PHC, and in samples enriched for less common diseases such as LBD and LATE-NC - will be necessary to establish generalizability.

## Declaration of generative AI and AI-assisted technologies in the manuscript preparation process

During the preparation of this work the authors used Claude (claude-sonnet-4-6, Anthropic, PBC) in order to improve the clarity and readability of the text, assist with structuring the manuscript, and detect errors, inconsistencies, and typos. After using this tool, the authors reviewed and edited the content as needed and take full responsibility for the content of the published article.

## Supporting information

Sfigure 13

SFigure 12

SFigure 11

SFigure 10

SFigure 9

SFigure 8

Sffigure 7

SFigure 6

SFigure 5

SFigure 4

SFigure 3

SFigure 2

SFigure 1

## Acknowledgments

This work was supported by NIH NIA grants U01 AG068057 (‘AI4AD’), U01AG082350 (‘CLARiTI’) and U24AG074855 (‘PHC’). Research reported in this publication was supported by NIH S10OD032285, 5R01EB017230 (Dr. Landman), R01AG082730 (Dr. Mukherjee), U24AG0748502 (Dr. Mukherjee), R01AG091657 (Dr. Tosun), R01AG062695 (Dr. Beecham) and K01-073584 (Dr. Archer). Dr. Saykin acknowledges support from multiple NIH grants (P30 AG010133, P30 AG072976, R01 AG019771, R01 AG057739, U19 AG024904, R01 LM013463, R01 AG068193, R01 AG092591, T32 AG071444, U01 AG068057, U01 AG072177, U19 AG074879, as well as U24 AG074855). The ADSP Phenotype Harmonization Consortium (ADSP-PHC) is funded by NIA (U24 AG074855, U01 AG068057 and R01 AG059716). Cohorts for this study included the Adult Changes in Thought study (ACT), U01 AG006781, U19 AG066567; National Alzheimer’s Coordinating Center (NACC): The NACC database is funded by NIA/NIH Grant U24 AG072122. SCAN is a multi-institutional project that was funded as a U24 grant (AG067418) by the National Institute on Aging in May 2020. Data collected by SCAN and shared by NACC are contributed by the NIA-funded ADRCs as follows: P30 AG062429 (PI James Brewer, MD, PhD), P30 AG066468 (PI Oscar Lopez, MD), P30 AG062421 (PI Bradley Hyman, MD, PhD), P30 AG066509 (PI Thomas Grabowski, MD), P30 AG066514 (PI Mary Sano, PhD), P30 AG066530 (PI Helena Chui, MD), P30 AG066507 (PI Marilyn Albert, PhD), P30 AG066444 (PI John Morris, MD), P30 AG066518 (PI Jeffrey Kaye, MD), P30 AG066512 (PI Thomas Wisniewski, MD), P30 AG066462 (PI Scott Small, MD), P30 AG072979 (PI David Wolk, MD), P30 AG072972 (PI Charles DeCarli, MD), P30 AG072976 (PI Andrew Saykin, PsyD), P30 AG072975 (PI David Bennett, MD), P30 AG072978 (PI Neil Kowall, MD), P30 AG072977 (PI Robert Vassar, PhD), P30 AG066519 (PI Frank LaFerla, PhD), P30 AG062677 (PI Ronald Petersen, MD, PhD), P30 AG079280 (PI Eric Reiman, MD), P30 AG062422 (PI Gil Rabinovici, MD), P30 AG066511 (PI Allan Levey, MD, PhD), P30 AG072946 (PI Linda Van Eldik, PhD), P30 AG062715 (PI Sanjay Asthana, MD, FRCP), P30 AG072973 (PI Russell Swerdlow, MD), P30 AG066506 (PI Todd Golde, MD, PhD), P30 AG066508 (PI Stephen Strittmatter, MD, PhD), P30 AG066515 (PI Victor Henderson, MD, MS), P30 AG072947 (PI Suzanne Craft, PhD), P30 AG072931 (PI Henry Paulson, MD, PhD), P30 AG066546 (PI Sudha Seshadri, MD), P20 AG068024 (PI Erik Roberson, MD, PhD), P20 AG068053 (PI Justin Miller, PhD), P20 AG068077 (PI Gary Rosenberg, MD), P20 AG068082 (PI Angela Jefferson, PhD), P30 AG072958 (PI Heather Whitson, MD), P30 AG072959 (PI James Leverenz, MD); National Institute on Aging Alzheimer’s Disease Family Based Study (NIA-AD FBS): U24 AG056270; Religious Orders Study (ROS): P30AG10161,R01AG15819, R01AG42210; Memory and Aging Project (MAP - Rush): R01AG017917, R01AG42210; Minority Aging Research Study (MARS): R01AG22018, R01AG42210. Additional acknowledgments include the National Institute on Aging Genetics of Alzheimer’s Disease Data Storage Site (NIAGADS, U24AG041689) at the University of Pennsylvania, funded by NIA.

## References

1. Jellinger KA. Recent update on the heterogeneity of the Alzheimer’s disease spectrum. J Neural Transm. 2022;129(1):1–24.

2. Nichols E, Steinmetz JD, Vollset SE, Fukutaki K, Chalek J, Abd-Allah F, et al. Estimation of the global prevalence of dementia in 2019 and forecasted prevalence in 2050: an analysis for the Global Burden of Disease Study 2019. Lancet Public Health. 2022 Feb;7(2):e105–e125. doi: 10.1016/S2468-2667(21)00249-8. Epub 2022 Jan 6. PMID: 34998485; PMCID: PMC8810394.

3. 2023 Alzheimer’s disease facts and figures. Alzheimers Dement. 2023 Apr;19(4):1598-1695. doi: 10.1002/alz.13016. Epub 2023 Mar 14. PMID: 36918389.

4. Jack CR, Bennett DA, Blennow K, Carrillo MC, Dunn B, Haeberlein AB, et al. NIAA-AA Research Framework: Toward a biological definition of Alzheimer’s disease. Alzheimers Dement. 2018 Apr;14(4):535–62. doi: 10.1016/j.jalz.2018.02.018.

5. Jack CR, Andrews JS, Beach TG, et al. Revised criteria for diagnosis and staging of Alzheimer’s disease: Alzheimer’s Association Workgroup. Alzheimer’s Dement. 2024;20:5143–5169. 10.1002/alz.13859

6. Schneider JA, Arvanitakis Z, Bang W, Bennett DA. Mixed brain pathologies account for most dementia cases in community-dwelling older persons. Neurology. 2007 Dec 11;69(24):2197–204. doi: 10.1212/01.wnl.0000271090.28148.24. Epub 2007 Jun 13. PMID: 17568013.

7. Wilson RS, Yu L, Trojanowski JQ, Chen EY, Boyle PA, Bennett DA, et al. TDP-43 pathology, cognitive decline, and dementia in old age. JAMA Neurol. 2013;70(11):1418–24.

8. Boyle PA, Yu L, Wilson RS, Leurgans SE, Schneider JA, Bennett DA. Person-specific contribution of neuropathologies to cognitive loss in old age. Ann Neurol. 2018 Jan;83(1):74–83. doi: 10.1002/ana.25123. Epub 2018 Jan 14. PMID: 29244218; PMCID: PMC5876116.

9. Weller RO, Boche D, Nicoll JA. Microvasculature changes and cerebral amyloid angiopathy in Alzheimer’s disease and their potential impact on therapy. Acta Neuropathol. 2009;118(1):87–102

10. Nelson PT, Schmitt FA, Lin Y, Abner EL, Jicha GA, Patel E, Thomason PC, Neltner JH, Smith CD, Santacruz KS, Sonnen JA, Poon LW, Gearing M, Green RC, Woodard JL, Van Eldik LJ, Kryscio RJ. Hippocampal sclerosis in advanced age: clinical and pathological features. Brain. 2011a May;134(Pt 5):1506–18. doi: 10.1093/brain/awr053. PMID: 21596774; PMCID: PMC3097889.

11. Thomas DX, Bajaj S, McRae-McKee K, Hadjichrysanthou C, Anderson RM, Collinge J. Association of TDP-43 proteinopathy, cerebral amyloid angiopathy, and Lewy bodies with cognitive impairment in individuals with or without Alzheimer’s disease neuropathology. Sci Rep. 2020;10(1):14579.

12. Rabin JS, Nichols E, LaJoie R, Casaletto KB, Palta P, Dams-O’Connor K, et al. Cerebral amyloid angiopathy interacts with neuritic amyloid plaques to promote tau and cognitive decline. Brain. 2022;145(8):2823–33.

13. Woodworth DC, Nguyen KM, Sordo L, Scambray KA, Head E, Kawas CH, et al. Comprehensive assessment of TDP-43 neuropathology data in the National Alzheimer’s Coordinating Center database. Acta Neuropathol. 2024;147(1):103. Epub 20240619. doi: 10.1007/s00401-024-02728-8. PMCID: PMC11186885.

14. van Dyck CH, Swanson CJ, Aisen P, Bateman RJ, Chen C, Gee M, et al. Lecanemab in Early Alzheimer’s Disease. N Engl J Med. 2023 Jan 5;388(1):9–21. doi: 10.1056/NEJMoa2212948. Epub 2022 Nov 29. PMID: 36449413.

15. Sperling RA, Donohue MC, Raman R, Rafii MS, Johnson K, Masters CL, et al. Trial of Solanezumab in Preclinical Alzheimer’s Disease. N Engl J Med. 2023 Sep 21;389(12):1096–1107. doi: 10.1056/NEJMoa2305032. Epub 2023 Jul 17. PMID: 37458272; PMCID: PMC10559996.

16. Sims JR, Zimmer JA, Evans CD, Lu M, Ardayfio P, Sparks J, Wessels AM, Shcherbinin S, Wang H, Nery ES, Collins EC, Solomon P, Salloway S, Apostolova LG, Hansson O, Ritchie C, Brooks DA, Mintun MA, Skovronsky DM; TRAILBLAZER-ALZ 2 Investigators. Donanemab in early symptomatic Alzheimer disease: the TRAILBLAZER-ALZ 2 randomized clinical trial. JAMA. 2023 Aug 8;330(6):512–527. doi: 10.1001/jama.2023.13239. PMID: 37459141; PMCID: PMC10352931.

17. Tosun D, Yardibi O, Benzinger TLS, Kukull WA, Masters CL, Perrin RJ, et al. Identifying individuals with non-Alzheimer’s disease co-pathologies: a precision medicine approach to clinical trials in sporadic Alzheimer’s disease. Alzheimers Dement. 2024;20(1):421–36.

18 Kapasi A, James BD, Yu L, Sood A, Arvanitakis Z, Bennett DA, et al. Mixed Pathologies and Cognitive Outcomes in Persons Considered for Anti-Amyloid Treatment Eligibility Assessment: A Community-Based Study. Neurology. 2025;105(5):e214004. Epub 20250818. doi: 10.1212/WNL.0000000000214004. PMCID: PMC12367424.

19. Cummings JL, Morstorf T, Zhong K. Alzheimer’s disease drug-development pipeline: few candidates, frequent failures. Alzheimers Res Ther. 2014 Jul 3;6(4):37. doi: 10.1186/alzrt269.

20. Mehta SJ, Volpp KG, Troxel AB, Day SC, Lim R, Marcus N, Norton L, Anderson S, Asch DA. Electronic Pill Bottles or Bidirectional Text Messaging to Improve Hypertension Medication Adherence (Way 2 Text): a Randomized Clinical Trial. J Gen Intern Med. 2019 Nov;34(11):2397–2404. doi: 10.1007/s11606-019-05241-x. Epub 2019 Aug 8. PMID: 31396815; PMCID: PMC6848522.

21. Duara R, Barker W. Heterogeneity in Alzheimer’s disease diagnosis and progression rates: implications for therapeutic trials. Neurotherapeutics. 2022;19(1):8–25.

22. Wang X, Ye T, Jiang D, Zhou W, Zhang J. Characterizing the clinical heterogeneity of early symptomatic Alzheimer’s disease: a data-driven machine learning approach. Front Aging Neurosci. 2024;16:1410544.

23. Jack CR, Knopman DS, Jagust WJ, Shaw LM, Aisen PS, Weiner MW, et al. Hypothetical model of dynamic biomarkers of the Alzheimer’s pathological cascade. Lancet Neurol. 2010;9(1):119–28.

24. Dichgans M, Zietemann V. Prevention of vascular cognitive impairment. Stroke. 2012 Nov;43(11):3137–46. doi: 10.1161/STROKEAHA.112.651778. Epub 2012 Aug 30. PMID: 22935401.

25. Thompson PM, Vinters HV. Pathologic lesions in neurodegenerative diseases. Prog Mol Biol Transl Sci. 2012;107:1–40.

26. Wardlaw JM, Smith C, Dichgans M. Mechanisms of sporadic cerebral small vessel disease: insights from neuroimaging. Lancet Neurol. 2013 May;12(5):483–97. doi: 10.1016/S1474-4422(13)70060-7. Erratum in: Lancet Neurol. 2013 Jun;12(6):532. PMID: 23602162; PMCID: PMC3836247.

27. McKeith IG, Boeve BF, Dickson DW, Halliday G, Taylor JP, Weintraub D, et al. Diagnosis and management of dementia with Lewy bodies: Fourth consensus report of the DLB Consortium. Neurology. 2017 Jul 4;89(1):88–100. doi: 10.1212/WNL.0000000000004058. Epub 2017 Jun 7. PMID: 28592453; PMCID: PMC5496518.

28. Nelson PT, Dickson DW, Trojanowski JQ, Jack CR, Boyle PA, Arfanakis K, et al. Limbic-predominant age-related TDP-43 encephalopathy (LATE): consensus working group report. Brain. 2019;142(6):1503–27.

29. Espay AJ, Lees AJ, Cardoso F, Frucht SJ, Erskine D, Sandoval IM, et al. The α-synuclein seed amplification assay: interpreting a test of Parkinson’s pathology. Parkinsonism Relat Disord. 2025;131:107256.

30. Hansson O, Edelmayer RM, Boxer AL, Carrillo MC, Mielke MM, Rabinovici GD, et al. The Alzheimer’s Association appropriate use recommendations for blood biomarkers in Alzheimer’s disease. Alzheimers Dement. 2022 Dec;18(12):2669–2686. doi: 10.1002/alz.12756. Epub 2022 Jul 31. PMID: 35908251; PMCID: PMC10087669.

31. Hansson O, Blennow K, Zetterberg H, Dage J. Blood biomarkers for Alzheimer’s disease in clinical practice and trials. JAMA Neurol. Published online 2023.

32. Ashton NJ, Brum WS, Di Molfetta G, Benedet AL, Arslan B, Jonaitis E, et al. Diagnostic Accuracy of a Plasma Phosphorylated Tau 217 Immunoassay for Alzheimer Disease Pathology. JAMA Neurol. 2024 Mar 1;81(3):255–263. doi: 10.1001/jamaneurol.2023.5319. PMID: 38252443; PMCID: PMC10804282.

33. Palmqvist S, Tideman P, Mattsson-Carlgren N, Schindler SE, Smith R, Ossenkoppele R, et al. Blood Biomarkers to Detect Alzheimer Disease in Primary Care and Secondary Care. JAMA. 2024 Oct 15;332(15):1245–1257. doi: 10.1001/jama.2024.13855. PMID: 39068545; PMCID: PMC11284636.

34. Therriault J, Janelidze S, Benedet AL, Ashton NJ, Arranz Martinez J, Gonzalez-Escalante A, et al. Diagnosis of Alzheimer’s disease using plasma biomarkers adjusted to clinical probability. Nat Aging. 2024;4(11):1529–37. Epub 20241112. doi: 10.1038/s43587-024-00731-y. PMCID: PMC11564087.

35. Foulds PG, Walker RM, Farrell M, O’Dowd S, Alladi S, Bak TH, et al. TAR DNA-binding protein-43 in amyotrophic lateral sclerosis and frontotemporal lobar degeneration. Acta Neuropathol. 2008 Jun;116(2):135–40. doi: 10.1007/s00401-008-0405-z

36. McMillan M, Gomez N, Hsieh C, Bekier M, Li X, Miguez R, Tank EMH, Barmada SJ. RNA methylation influences TDP43 binding and disease pathogenesis in models of amyotrophic lateral sclerosis and frontotemporal dementia. Mol Cell. 2023 Jan 19;83(2):219–236.e7. doi: 10.1016/j.molcel.2022.12.019. Epub 2023 Jan 11. PMID: 36634675; PMCID: PMC9899051.

37. Dark HE, Duggan MR, Walker KA. Plasma biomarkers for Alzheimer’s and related dementias: a review and outlook for clinical neuropsychology. Arch Clin Neuropsychol. 2024;39(3):313–24.

38. Vrillon A, Bousiges O, Götze K, Demuynck C, Muller C, Ravier A, et al. Plasma biomarkers of amyloid, tau, axonal, and neuroinflammation pathologies in dementia with Lewy bodies. Alzheimers Res Ther. 2024;16:146.

39. Wang D, Honnorat N, Toledo JB, Li K, Charisis S, Rashid T, et al. Deep learning reveals pathology-confirmed neuroimaging signatures in Alzheimer’s, vascular and Lewy body dementias. Brain. 2025;148(6):1963–77.

40. Chattopadhyay T, Kush R, Senthilkumar P, Patterson C, Owens-Walton C, Gleave EJ, et al. Evaluation of deep learning algorithms to predict multiple dementia-related neuropathologies from brain MRI, clinical and genetic data. In: Proceedings of the 21st International Symposium on Biomedical Image Processing and Analysis (SIPAIM); 2025; Pasto, Colombia. IEEE; 2025a. p. 1–4.

41. Dickerson BC, Bakkour A, Salat DH, Feczko E, Pacheco J, Greve DN, et al. The cortical signature of Alzheimer’s disease: regionally specific cortical thinning relates to symptom severity in very mild to mild AD dementia and is detectable in asymptomatic amyloid-positive individuals. Cereb Cortex. 2009 Mar;19(3):497–510. doi: 10.1093/cercor/bhn113.

42. Acosta-Cabronero J, Cardenas-Blanco A, Betts MJ, Booth T, Martínez-Herraez J, Nestor PJ; Alzheimer’s Disease Neuroimaging Initiative. The whole-brain pattern of magnetic susceptibility perturbations in Parkinson’s disease. Brain. 2017 Jan;140(1):118–131. doi: 10.1093/brain/aww278. Epub 2016 Nov 11

43. Chandio BQ, Villalon-Reina JE, Nir TM, Thomopoulos SI, Feng Y, Benavidez S, et al. Amyloid, Tau, and APOE in Alzheimer’s Disease: Impact on White Matter Tracts. bioRxiv [Preprint]. 2024 Aug 6:2024.08.05.606560. [Version 2] doi: 10.1101/2024.08.05.606560

44. Xu FH, Duong-Tran D, Huang H, Saykin AJ, Thompson PM, Davatzikos C, et al. Gene-modulated network diffusion models for improved modeling of amyloid-β pathophysiological spread in Alzheimer’s disease. IEEE Trans Comput Biol Bioinform. 2026. Submitted.

45. Chen KT, Gong E, de Carvalho Macruz FB, Xu J, Boumis A, Khalighi M, et al. Ultra-low-dose 18F-florbetaben amyloid PET imaging using deep learning with multi-contrast MRI inputs. Radiology. 2019;290(3):649–56.

46. Jin Y, DuBois J, Zhao C, Zhan L, Gabelle A, Jahanshad N, et al. Brain MRI to PET synthesis and amyloid estimation in Alzheimer’s disease via 3D multimodal contrastive GAN. In: Machine Learning in Medical Imaging. Cham: Springer Nature Switzerland; 2023. p. 94–103.

47. Makkinejad N, Evia AM, Tamhane AA, Javierre-Petit C, Leurgans SE, Lamar M, Barnes LL, et al. ARTS: A novel In-vivo classifier of arteriolosclerosis for the older adult brain. Neuroimage Clin. 2021;31:102768. doi: 10.1016/j.nicl.2021.102768. Epub 2021 Jul 24. PMID: 34330087; PMCID: PMC8329541.

48. Chattopadhyay T, Thomopoulos SI, Thompson PM. Predicting amyloid positivity from hippocampal and entorhinal cortex volume and APOE genotype. Alzheimers Dement. 2022;18:e067902.

49. Dhinagar NJ, Thomopoulos SI, Laltoo E, Thompson PM. Efficiently training vision transformers on structural MRI scans for Alzheimer’s Disease detection. In: Proceedings of the 45th Annual International Conference of the IEEE Engineering in Medicine and Biology Society; 2023a; Sydney, Australia. IEEE; 2023. p. 1–6.

50. Dhinagar NJ, Thomopoulos SI, Rajagopalan P, Stripelis D, Ambite JL, Ver Steeg G, et al. Evaluation of transfer learning methods for detecting Alzheimer’s disease with brain MRI. Proc SPIE. 2023b;12567:504–13.

51. Chattopadhyay T, Ozarkar SS, Buwa K, Joshy NA, Komandur D, Naik J, et al. Comparison of deep learning architectures for predicting amyloid positivity in Alzheimer’s disease, mild cognitive impairment, and healthy aging, from T1-weighted brain structural MRI. Front Neurosci. 2024;18:1387196.

52. Kapasi A, Wagner M, Evia AM, Fleischman DA, Boyle P, Marquez D, et al. Association of an in vivo classifier for ARTerioloSclerosis (ARTS) with cortical thickness and cognition in older adults. Neurobiol Aging. 2025 Dec;156:143–149. doi: 10.1016/j.neurobiolaging.2025.09.006. Epub 2025 Sep 17. PMID: 40974878.

53. Kumar R, Rashid T, Charisis S, Ho NH, Brandigampala SR, Toga AW, et al. Bridging neuroimaging and pathology in dementia: a multi-cohort investigation of MRI-derived brain volumes. Alzheimers Dement. 2025;21:e105854.

54. Chattopadhyay T, Ozarkar SS, Buwa K, Thomopoulos SI, Thompson PM; Alzheimer’s Disease Neuroimaging Initiative. Predicting Brain Amyloid Positivity from T1 weighted brain MRI and MRI-derived Gray Matter, White Matter and CSF maps using Transfer Learning on 3D CNNs. bioRxiv [Preprint]. 2023a Feb 16:2023.02.15.528705. doi: 10.1101/2023.02.15.528705. PMID: 36824826; PMCID: PMC9949045.

55. Chattopadhyay T, Komandur D, Naik J, Buwa K, Ozarkar SS, Thompoulos SI, et al. “Can Structural MRIs be used to Detect Amyloid Positivity using Deep Learning?,” 2023 19th International Symposium on Medical Information Processing and Analysis (SIPAIM), Mexico City, Mexico, 2023b, pp. 1-4, doi: 10.1109/SIPAIM56729.2023.10373510.

56. Erickson N, Mueller J, Shirkov A, Zhang H, Larson P, Li M, et al. AutoGluon-Tabular: robust and accurate AutoML for structured data. arXiv preprint arXiv:2003.06505. 2020.

57. He X, Zhao K, Chu X. AutoML: A survey of the state-of-the-art. Knowl Based Syst. 2021 Jan 15;212:106622. doi: 10.1016/j.knosys.2020.106622

58. Gijsbers P, Bueno MLP, Coors S, LeDell E, Poirier S, Thomas J, et al. AMLB: an AutoML Benchmark. J Mach Learn Res. 2024;25(101):1–65. 10.48550/arXiv.2207.12560

59. Hohman TJ, Cuccaro ML, Toga AW, et al. ADSP Phenotype Harmonization Consortium. Alzheimers Dement. 2023;19(S14):e077713.

60. Sonnen JA, Larson EB, Haneuse S, Woltjer R, Li G, Crane PK, Craft S, Montine TJ. Neuropathology in the adult changes in thought study: a review. J Alzheimers Dis. 2009;18(3):703–11. doi: 10.3233/JAD-2009-1180. PMID: 19661627; PMCID: PMC4008877.

61. Bennett DA, Buchman AS, Boyle PA, Barnes LL, Wilson RS, Schneider JA. Religious Orders Study and Rush Memory and Aging Project. J Alzheimers Dis. 2018;64(s1):S161-S189. doi: 10.3233/JAD-179939. PMID: 29865057; PMCID: PMC6380522.

62. Bennett DA, Schneider JA, Buchman AS, Barnes LL, Boyle PA, Wilson RS. Overview and findings from the Rush Memory and Aging Project. Curr Alzheimer Res. 2012 Jul;9(6):646–63. doi: 10.2174/156720512801322663. PMID: 22471867; PMCID: PMC3439198.

63. Barnes LL, Shah RC, Aggarwal NT, Bennett DA, Schneider JA. The Minority Aging Research Study: ongoing efforts to obtain brain donation in African Americans without dementia. Curr Alzheimer Res. 2012 Jul;9(6):734–45. doi: 10.2174/156720512801322627. PMID: 22471868; PMCID: PMC3409294.

64. Beekly, D.L., et. al. (2007). The National Alzheimer’s Coordinating Center (NACC) database: the Uniform Data Set. Alzheimer Disease and Associated Disorders, 21(3), 249–258.

65. Thal DR, Rüb U, Orantes M, & Braak H. Phases of A beta-deposition in the human brain and its relevance for the development of AD. Neuropathology and Applied Neurobiology. 2002;28(4):335–43. DOI: 10.1046/j.1365-2990.2002.00398.x

66. Fillenbaum GG, van Belle G, Morris JC, Mohs RC, Mirra SS, Davis PC, et al. CERAD (Consortium to Establish a Registry for Alzheimer’s Disease) The first 20 years. Alzheimers Dement. 2008 Mar;4(2):96–109. doi: 10.1016/j.jalz.2007.08.005

67. Braak H, Alafuzoff I, Arzberger T, Kretzschmar H, Tredici KD. Staging of Alzheimer disease-associated neurofibrillary pathology using paraffin sections and immunocytochemistry. Acta Neuropathol. 2006 Aug 12;112(4):389–404. doi: 10.1007/s00401-006-0127-z

68. Montine TJ, Phelps CH, Beach TG, Bigio EH, Cairns NJ, Dickson DW, et al. National Institute on Aging-Alzheimer’s Association guidelines for the neuropathologic assessment of Alzheimer’s disease: a practical approach. Acta Neuropathol. 2012 Jan;123(1):1–11. doi: 10.1007/s00401-011-0910-3. Epub 2011 Nov 20. PMID: 22101365; PMCID: PMC3268003.

69. Vonsattel JP, Myers RH, Hedley-Whyte ET, Ropper AH, Bird ED, Richardson EP. Cerebral amyloid angiopathy without and with cerebral hemorrhages: a comparative histological study. Ann Neurol. Nov 1991;30(5):637–649.

70. Olichney JM, Hansen LA, Hofstetter CR, Lee JH, Katzman R, Thal LJ. Association between severe cerebral amyloid angiopathy and cerebrovascular lesions in Alzheimer disease is not a spurious one attributable to apolipoprotein E4. Arch Neurol. Jun 2000;57(6):869–874.

71. Nelson PT, Smith CD, Abner EL, Wilfred BJ, Wang WX, Neltner JH, Baker M, Fardo DW, Kryscio RJ, Scheff SW, Jicha GA, Jellinger KA, Van Eldik LJ, Schmitt FA. Hippocampal sclerosis of aging, a prevalent and high-morbidity brain disease. Acta Neuropathol. 2013 Aug;126(2):161–177. doi: 10.1007/s00401-013-1154-1. PMID: 23864344; PMCID: PMC3744371.

72. Braak H, Tredici KD, Rüb U, de Vos RAI, Jansen-Steur ENH, Braak E. Staging of brain pathology related to sporadic Parkinson’s disease. Neurobiol Aging. 2003 Mar-Apr;24(2):197-211. doi: 10.1016/s0197-4580(02)00065-9.

73. McKeith IG, Dickson DW, Lowe J, Emre M, O’Brien JT, Feldman H, et al; Consortium on DLB. Diagnosis and management of dementia with Lewy bodies: third report of the DLB Consortium. Neurology. 2005 Dec 27;65(12):1863–72. doi: 10.1212/01.wnl.0000187889.17253.b1. Epub 2005 Oct 19. Erratum in: Neurology. 2005 Dec 27;65(12):1992. PMID: 16237129.

74. Braak H, Del Tredici K. Neuropathological Staging of Brain Pathology in Sporadic Parkinson’s disease: Separating the Wheat from the Chaff. J Parkinsons Dis. 2017;7(s1):S71-S85. doi: 10.3233/JPD-179001. PMID: 28282810; PMCID: PMC5345633.

75. Cairns NJ, Neumann M, Bigio EH, Holm IE, Troost D, Hatanpaa KJ, et al. TDP-43 in familial and sporadic frontotemporal lobar degeneration with ubiquitin inclusions. Am J Pathol. 2007 Jul;171(1):227–40. doi: 10.2353/ajpath.2007.070182. PMID: 17591968; PMCID: PMC1941578.

76. Josephs KA, Murray ME, Whitwell JL, Parisi JE, Petrucelli L, Jack CR, Petersen RC, Dickson DW. Staging TDP-43 pathology in Alzheimer’s disease. Acta Neuropathol. 2014 Mar;127(3):441–50. doi: 10.1007/s00401-013-1211-9. Epub 2013 Nov 16. PMID: 24240737; PMCID: PMC3944799.

77. Alosco ML, Stein TD, Tripodis Y, Chua AS, Kowall NW, Huber BR, et al. Association of White Matter Rarefaction, Arteriolosclerosis, and Tau With Dementia in Chronic Traumatic Encephalopathy. JAMA Neurol. 2019 Nov 1;76(11):1298–1308. doi: 10.1001/jamaneurol.2019.2244. PMID: 31380975; PMCID: PMC6686769.

78. Beach TG, Sue LI, Scott S, Intorcia AJ, Walker JE, Arce RA, et al. Cerebral white matter rarefaction has both neurodegenerative and vascular causes and may primarily be a distal axonopathy. J Neuropathol Exp Neurol. 2023 May 25;82(6):457–466. doi: 10.1093/jnen/nlad026. PMID: 37071794; PMCID: PMC10209646.

79. Roy O, Vetterli M. The effective rank: A measure of effective dimensionality. In: 2007 15th European Signal Processing Conference. IEEE; 2007. p. 606

80. Cangelosi R, Goriely A. Component retention in principal component analysis with application to cDNA microarray data. Biol Direct. 2007 Jan 17;2:2. doi: 10.1186/1745-6150-2-2. PMID: 17229320; PMCID: PMC1797006.

81. Fischl B. FreeSurfer. NeuroImage. 2012;62(2):774–81.

82. Fischl B, Dale AM. Measuring the thickness of the human cerebral cortex from magnetic resonance images. Proc Natl Acad Sci U S A. 2000;97(20):11050–5.

83. Johnson WE, Li C, & Rabinovic A. Adjusting batch effects in microarray expression data using empirical Bayes methods. Biostatistics. 2007;8(1), 118–127. DOI: 10.1093/biostatistics/kxj037

84. Fortin JP, Parker D, Tunç B, Watanabe T, Elliott MA, Ruparel K, et al. Harmonization of multi-site diffusion tensor imaging data. Neuroimage. 2017 Nov 1;161:149–170. doi: 10.1016/j.neuroimage.2017.08.047. Epub 2017 Aug 18. PMID: 28826946; PMCID: PMC5736019.

85. Buckner RL, Head D, Parker J, Fotenos AF, Marcus D, Morris JC, Snyder AZ. A unified approach for morphometric and functional data analysis in young, old, and demented adults using automated atlas-based head size normalization: reliability and validation against manual measurement of total intracranial volume. Neuroimage. 2004 Oct;23(2):724–38. doi: 10.1016/j.neuroimage.2004.06.018. PMID: 15488422.

86. Stern Y, Barnes CA, Grady C, Jones RN, Raz N. Brain reserve, cognitive reserve, compensation, and maintenance: operationalization, validity, and mechanisms of cognitive resilience. Neurobiol Aging. 2019 Nov;83:124–129. doi: 10.1016/j.neurobiolaging.2019.03.022. PMID: 31732015; PMCID: PMC6859943.

87. Mukherjee S, Choi SE, Lee ML, Scollard P, Trittschuh EH, Mez J, et al. Cognitive domain harmonization and cocalibration in studies of older adults. Neuropsychology. 2023;37(4):409–23

88. Kang K, Zhang P, Dumitrescu L, Mukherjee S, Lee ML, Choi SE, et al. The dynamics of cognitive decline toward Alzheimer’s disease progression: results from ADSP-PHC’s harmonized cognitive composites. Alzheimers Dement. 2025;21(6):e70335.

89. Besser L, Kukull W, Knopman DS, Chui H, Galasko D, Weintraub S, et al. Version 3 of the National Alzheimer’s Coordinating Center’s Uniform Data Set. Alzheimer Dis Assoc Disord. 2018;32(4):351–8. doi: 10.1097/WAD.0000000000000279. PMID: 30376508; PMCID: PMC6249084.

90. Timsina J, Ali M, Do A, Wang L, Western D, Sung YJ, Cruchaga C. Harmonization of CSF and imaging biomarkers in Alzheimer’s disease: Need and practical applications for genetics studies and preclinical classification. Neurobiol Dis. 2024 Jan;190:106373. doi: 10.1016/j.nbd.2023.106373. Epub 2023 Dec 9. PMID: 38072165; PMCID: PMC12661535.

91. Peter C, Sathe A, Shashikumar N, Pechman KR, Workmeister AW, Jackson TB, et al. White Matter Abnormalities and Cognition in Aging and Alzheimer Disease. JAMA Neurol. 2025;82(8):825–836. doi:10.1001/jamaneurol.2025.1601

92. Mori S, Oishi K, Jiang H, Jiang L, Li X, Akhter K, et al. Stereotaxic white matter atlas based on diffusion tensor imaging in an ICBM template. NeuroImage. 2008;40(2):570–82.

93. Oishi K, Faria A, Jiang H, Li X, Akhter K, Zhang J, et al. Atlas-based whole brain white matter analysis using large deformation diffeomorphic metric mapping: application to normal elderly and Alzheimer’s disease participants. NeuroImage. 2009;46(2):486–99.

94. Fortin JP, Cullen N, Sheline YI, Taylor WD, Aselcioglu I, Cook PA, et al. Harmonization of cortical thickness measurements across scanners and sites. Neuroimage. 2018 Feb 15;167:104–120. doi: 10.1016/j.neuroimage.2017.11.024. Epub 2017 Nov 17. PMID: 29155184; PMCID: PMC5845848.

95. Chattopadhyay T, Senthilkumar P, Ankarath RH, Patterson C, Gleave EJ, Thomopoulos SI, et al. Predicting Longitudinal Cognitive Decline via Multimodal Deep Learning Fusing Brain MRI and Clinical Data. In 2025 21st International Symposium on Biomedical Image Processing and Analysis (SIPAIM). IEEE. 2025b Nov:1-4.

96. Chattopadhyay T, Senthilkumar P, Ankarath RH, Patterson C, Gleave EJ, Thomopoulos SI, Huang H, Shen L, You L, Zhi D and Thompson PM (2026) Deep learning to predict future cognitive decline: a multimodal approach using brain MRI and clinical data. Front. Neuroimaging. 2026;5:1726037. doi: 10.3389/fnimg.2026.1726037

97. Gijsbers P, Bueno MLP, Coors S, LeDell E, Poirier S, Thomas J, et al. AMLB: an AutoML Benchmark. J Mach Learn Res. 2024;25(1):4978–5042. Article 101. Available from: https://www.jmlr.org/papers/volume25/22-0493/22-0493.pdf.

98. Wolpert D. Stacked generalization. Neural Netw. 1992;5(2):241–259.

99. Matthews BW. Comparison of the predicted and observed secondary structure of T4 phage lysozyme. Biochim Biophys Acta. 1975;405(2):442–51.

100. Chicco D, Jurman G. The advantages of the Matthews correlation coefficient (MCC) over F1 score and accuracy in binary classification evaluation. BMC Genomics. 2020;21(1):6.

101. Kohavi R. A study of cross-validation and bootstrap for accuracy estimation and model selection. In: Proceedings of the 14th International Joint Conference on Artificial Intelligence (IJCAI’95); 1995 Aug 20-25; Montreal, Quebec, Canada. San Francisco (CA): Morgan Kaufmann; 1995. p. 1137–1143.

102. Varoquaux G. Cross-validation failure: Small sample sizes lead to large error bars. Neuroimage. 2018 Oct 15;180(Pt A):68–77. doi: 10.1016/j.neuroimage.2017.06.061. Epub 2017 Jun 24. PMID: 28655633.

103. Gorodkin J. Comparing two K-category assignments by a K-category correlation coefficient. Comput Biol Chem. 2004;28(5–6):367–74.

104. Grandini M, Bagli E, Visani G. Metrics for multi-class classification: an overview. arXiv preprint arXiv:2008.05756. 2020.

105. Itaya Y, Tamura J, Hayashi K, Yamamoto K. Asymptotic properties of Matthews correlation coefficient. Stat Med. 2025;44(1–2):e10303.

106. Spearman C. The proof and measurement of association between two things. Am J Psychol. 1904;15(1):72–101.

107. Conover WJ. Practical Nonparametric Statistics. 3rd ed. New York: Wiley; 1999.

108. Brodersen KH, Ong CS, Stephan KE, Buhmann JM. The balanced accuracy and its posterior distribution. In: 2010 20th International Conference on Pattern Recognition; 2010 Aug 23-26; Istanbul, Turkey. Piscataway (NJ): IEEE; 2010. p. 3121–4.

109. Benjamini Y, Hochberg Y. Controlling the false discovery rate: a practical and powerful approach to multiple testing. J R Stat Soc Series B Stat Methodol. 1995;57(1):289–300.

110. Frank E, Hall M. A Simple Approach to Ordinal Classification. In: De Raedt, L., Flach, P. (eds) Machine Learning: ECML 2001. ECML 2001. Lecture Notes in Computer Science(), vol 2167. Springer, Berlin, Heidelberg. 10.1007/3-540-44795-4_13

111. Blennow K, Hampel H, Weiner M, Zetterberg H. Cerebrospinal fluid and plasma biomarkers in Alzheimer disease. Nat Rev Neurol. 2010 Mar;6(3):131–44. doi: 10.1038/nrneurol.2010.4. Epub 2010 Feb 16. PMID: 20157306.

112. Shaw LM, Vanderstichele H, Knapik-Czajka M, Clark CM, Aisen PS, Petersen RC, Blennow K, Soares H, Simon A, Lewczuk P, Dean R, Siemers E, Potter W, Lee VM, Trojanowski JQ; Alzheimer’s Disease Neuroimaging Initiative. Cerebrospinal fluid biomarker signature in Alzheimer’s disease neuroimaging initiative subjects. Ann Neurol. 2009 Apr;65(4):403–13. doi: 10.1002/ana.21610. PMID: 19296504; PMCID: PMC2696350.

113. Mattsson N, Andreasson U, Zetterberg H, Blennow K, for the Alzheimer’s Disease Neuroimaging Initiative. Association of Plasma Neurofilament Light With Neurodegeneration in Patients With Alzheimer Disease. JAMA Neurol. 2017;74(5):557–566. doi:10.1001/jamaneurol.2016.6117

114. Burke BT, Latimer C, Keene CD, Sonnen JA, McCormick W, Bowen JD, et al. Theoretical impact of the AT(N) framework on dementia using a community autopsy sample. Alzheimers Dement. 2021 Dec;17(12):1879–1891. doi: 10.1002/alz.12348. Epub 2021 Apr 26. PMID: 33900044; PMCID: PMC8875276.

115. Jack CR Jr, Knopman DS, Jagust WJ, Shaw LM, Aisen PS, Weiner MW, et al. Tracking pathophysiological processes in Alzheimer’s disease: an updated hypothetical model of dynamic biomarkers. Lancet Neurol. 2013;12(2):207–16.

116. Siderowf A, Concha-Marambio L, Lafontant DE, Farris CM, Ma Y, Urenia PA, et al. Assessment of heterogeneity among participants in the Parkinson’s Progression Markers Initiative cohort using α-synuclein seed amplification: a cross-sectional study. Lancet Neurol. 2023 May;22(5):407–417. doi: 10.1016/S1474-4422(23)00109-6. PMID: 37059509; PMCID: PMC10627170.

117. Zheng Y, Li S, Yang C, Yu Z, Jiang Y, Feng T. Comparison of biospecimens for α-synuclein seed amplification assays in Parkinson’s disease: a systematic review and network meta-analysis. Eur J Neurol. 2023;30(12):3949–67.

118. Neltner JH, Abner EL, Jicha GA, Schmitt FA, Patel E, Snowden T, Sherwood SN, Azam MT, Mog S, Kryscio RJ, Nelson PT. Brain pathologies in extreme old age. Neurobiol Aging. 2016 Apr;37:1–11. doi: 10.1016/j.neurobiolaging.2015.10.009. PMID: 26545630; PMCID: PMC4706522.

119. Risacher SL, Anderson WH, Charil A, Castelluccio PF, Shcherbinin S, Saykin AJ, et al. Alzheimer disease brain atrophy subtypes are associated with cognition and rate of decline. Neurology. 2017 Nov 21;89(21):2176–2186. doi: 10.1212/WNL.0000000000004670. Epub 2017 Oct 25. PMID: 29070667; PMCID: PMC5696639

120. Pasi M, van Uden IWM, Tuladhar AM, de Leeuw FE, Pantoni L. White matter microstructural damage on diffusion tensor imaging in cerebral small vessel disease. Stroke. 2015 Feb;46(2):572–577. doi: 10.1161/STROKEAHA.114.007445. PMID: 25523040.

121. Chen Y, Tozer D, Li R, Li H, Tuladhar A, De Leeuw FE, et al. Improved dementia prediction in cerebral small vessel disease using deep learning-derived diffusion scalar maps from T1. Stroke. 2023. doi:10.1161/STROKEAHA.124.047449.

122. Gorelick PB, Scuteri A, Black SE, Decarli C, Greenberg SM, Iadecola C, et al. Vascular contributions to cognitive impairment and dementia: a statement for healthcare professionals from the American Heart Association/American Stroke Association. Stroke. 2011 Sep;42(9):2672–713. doi: 10.1161/STR.0b013e3182299496. PMID: 21778438; PMCID: PMC3778669.

123. Sorond FA, Cruz-Almeida Y, Clark DJ, Viswanathan A, Scherzer CR, De Jager P, et al. Aging, the Central Nervous System, and Mobility in Older Adults: Neural Mechanisms of Mobility Impairment. J Gerontol A Biol Sci Med Sci. 2015 Dec;70(12):1526–32. doi: 10.1093/gerona/glv130. Epub 2015 Sep 18. PMID: 26386013; PMCID: PMC4643615.

124. Ihle A, Ghisletta P, Ballhausen N, Fagot D, Vallet F, Baeriswyl M, et al. The role of cognitive reserve accumulated in midlife for the relation between chronic diseases and cognitive decline in old age: A longitudinal follow-up across six years. Neuropsychologia. 2018 Dec 1;121:37–46.

125. Charidimou A, Boulouis G, Gurol ME, Ayata C, Bacskai BJ, Frosch MP, et al. Emerging concepts in sporadic cerebral amyloid angiopathy. Brain. 2017 Jul 1;140(7):1829–50. doi: 10.1093/brain/awx047. PMID: 28334869.

126. Greenberg SM, Charidimou A. Diagnosis of cerebral amyloid angiopathy: evolution of the Boston criteria. Stroke. 2018 Feb;49(2):491–7.

127. Charidimou A, Boulouis G, Frosch MP, Baron JC, Pasi M, Albucher JF, et al. The Boston criteria version 2.0 for cerebral amyloid angiopathy: a multicentre, retrospective, MRI-neuropathology diagnostic accuracy study. Lancet Neurol. 2022 Aug;21(8):714–725. doi: 10.1016/S1474-4422(22)00208-3.

128. Boyle PA, Yang J, Yu L, Leurgans SE, Capuano AW, Schneider JA, Wilson RS, Bennett DA. Varied effects of age-related neuropathologies on the trajectory of late life cognitive decline. Brain. 2017 Mar 1;140(3):804–812. doi: 10.1093/brain/aww341. PMID: 28053080; PMCID: PMC5278309.

129. Hiya S, Maldonado-Díaz C, Walker JM, Richardson TE. Cognitive symptoms progress with limbic-predominant age-related TDP-43 encephalopathy stage and co-occurrence with Alzheimer disease. J Neuropathol Exp Neurol. 2024;83(1):2–10.

130. Sajjadi SA, Khan Z, El-Khoury A, Bubbico G, Akhlaghipour G, Corrada MM, et al. Association of language markers with future cognitive impairment and presence of limbic predominant age related TDP-43 encephalopathy. Alzheimers Dement. 2025. doi:10.1002/alz.70168.

131. Butler-Pagnotti RM, Pudumjee SB, Cross CL, Miller JB. Cognitive and clinical characteristics of patients with limbic-predominant age-related TDP-43 encephalopathy. Neurology. 2023;100(16):e1682–93.

132. Wolk DA, Nelson PT, Apostolova L, Arfanakis K, Boyle PA, Carlsson CM, et al. Clinical criteria for limbic-predominant age-related TDP-43 encephalopathy. Alzheimers Dement. 2025;21:e14202.

133. Ferman TJ, Smith GE, Kantarci K, Boeve BF, Pankratz VS, Dickson DW, Graff-Radford NR, Wszolek Z, Van Gerpen J, Uitti R, Pedraza O, Murray ME, Aakre J, Parisi J, Knopman DS, Petersen RC. Nonamnestic mild cognitive impairment progresses to dementia with Lewy bodies. Neurology. 2013 Dec 3;81(23):2032–8. doi: 10.1212/01.wnl.0000436942.55281.47. Epub 2013 Nov 8. PMID: 24212390; PMCID: PMC3854825.

134. Alafuzoff I, Ince PG, Arzberger T, Al-Sarraj S, Bell J, Bodi I, et al. Staging/typing of Lewy body related α-synuclein pathology: a study of the BrainNet Europe Consortium. Acta Neuropathol. 2009;117(6):635–52.

135. Beach TG, Adler CH, Lue L, Sue LI, Bachalakuri J, Henry-Watson J, et al. Unified staging system for Lewy body disorders: correlation with nigrostriatal degeneration, cognitive impairment and motor dysfunction. Acta Neuropathol. 2009 Jun;117(6):613–634. doi: 10.1007/s00401-009-0538-8. PMID: 19399512; PMCID: PMC2689361.

136. Attems J, Toledo JB, Walker L, Gelpi E, Gentleman S, Halliday G, et al. Neuropathological consensus criteria for the evaluation of Lewy pathology in post-mortem brains: a multi-centre study. Acta Neuropathol. 2021;141(2):159–72.

137. Irwin KE, Jasin P, Braunstein KE, Sinha I, Bowden KD, Moghekar A, et al. A novel fluid biomarker for TDP-43 loss of function. Alzheimers Dement. 2023;19:e082829. doi: 10.1002/alz.082829. PMID: 37793833.

138. Bauer CE, Zachariou V, Sudduth TL, Van Eldik LJ, Jicha GA, Nelson PT, et al. Plasma TDP-43 levels are associated with neuroimaging measures of brain structure in limbic regions. Alzheimers Dement (Amst). 2023 May 31;15(2):e12437. doi: 10.1002/dad2.12437. PMID: 37266411; PMCID: PMC10231556.

139. Wilson RS, Yu L, Leurgans SE, Bennett DA, Boyle PA. Proportion of cognitive loss attributable to terminal decline. Neurology. 2020 Jan 7;94(1):e42–e50. doi: 10.1212/WNL.0000000000008671. PMID: 31792096.

140. Dodge S, Sharp M, Smith W, Massimo L. The Frontotemporal Degeneration (FTD) Clinical Research Learning Institute. Innov Aging. 2025 Dec 31;9(Suppl 2):igaf122.019. doi: 10.1093/geroni/igaf122.019. PMCID: PMC12759528.

141. McKeith IG, Dickson DW, Lowe J, Emre M, O’brien JT, Feldman H, et al. Diagnosis and management of dementia with Lewy bodies: third report of the DLB Consortium. Neurology. 2005 Dec 27;65(12):1863–72.

142. Ferman TJ, Smith GE, Boeve BF, Graff-Radford NR, Lucas JA, Knopman DS, et al. Neuropsychological differentiation of dementia with Lewy bodies from normal aging and Alzheimer’s disease. The Clinical Neuropsychologist. 2006 Dec 1;20(4):623–36.

143. Strobl C, Boulesteix AL, Zeileis A, Hothorn T. Bias in random forest variable importance measures: illustrations, sources and a solution. BMC Bioinformatics. 2007 Jan 25;8:25. doi: 10.1186/1471-2105-8-25. PMID: 17254353; PMCID: PMC1796903.

144. Suk HI, Lee SW, Shen D; Alzheimer’s Disease Neuroimaging Initiative. Hierarchical feature representation and multimodal fusion with deep learning for AD/MCI diagnosis. Neuroimage. 2014 Nov 1;101:569–82. doi: 10.1016/j.neuroimage.2014.06.077. Epub 2014 Jul 18. PMID: 25042445; PMCID: PMC4165842.

145. Cheerla A, Gevaert O. Deep learning with multimodal representation for pancancer prognosis prediction. Bioinformatics. 2019 Jul 15;35(14):i446–i454. doi: 10.1093/bioinformatics/btz342. PMID: 31510656; PMCID: PMC6612862.

146. Lyu X, Mundada NS, Brown CA, Sadeghpour N, McGrew E, Xie L, et al. Medial temporal lobe Tau-Neurodegeneration mismatch from MRI and plasma biomarkers identifies vulnerable and resilient phenotypes with AD. medRxiv. 2025. Epub 20250819. doi: 10.1101/2025.08.17.25333859. PMCID: PMC12393630.

147. Brown CA, Mundada NS, Cousins KAQ, Sadeghpour N, Lyu X, McGrew E, et al. Tau-Clinical Mismatch Identifies Individuals with Co-Pathology and Predicts Clinical Trajectory. medRxiv. 2025. Epub 20250725. doi: 10.1101/2025.07.25.25332195. PMCID: PMC12330401.

148. Nguyen AT, Kouri N, Labuzan SA, Przybelski SA, Lesnick TG, Raghavan S, et al. Neuropathologic scales of cerebrovascular disease associated with diffusion changes on MRI. Acta Neuropathol. 2022. doi:10.1007/s00401-022-02465-w.

149. Oliveira LC, Chauhan J, Chaudhari A, Cheung SCS, Patel V, Villablanca AC, et al. A machine learning approach to automate microinfarct and microhemorrhage screening in hematoxylin and eosin-stained human brain tissues. J Neuropathol Exp Neurol. 2025;84(2):114–24.

