## Supplemental Materials for "Predicting Autopsy-Confirmed Neuropathology across Clinical, Neuroimaging, and CSF Biomarkers using Machine Learning"

**5.1 Supplemental Results: Global Rank Correlation and Class-Specific Binary MCC at Ordinal Boundaries**

For each ordinal outcome, we report the global Spearman rank correlation (Rₛ) across all classes, followed by class-specific binary MCCs at each ordinal boundary (e.g., 0|1 separates classes ≤0 from ≥1), which quantify discrimination at clinically meaningful thresholds. Missing MCC values (NA) occur when no samples are assigned to one class at a boundary, typically reflecting extreme class imbalance in small sub-cohorts. Incremental improvement over matched Base Clinical predictions is reported as ΔMCC with significance. Each section concludes with a summary of the best-performing feature sets and the most informative boundaries.

**5.1.1 Thal Phase  (Supplementary Figure 1)**

Staging. The Thal Phase is a 6-class system (0–5) tracking the progressive spatial spread of β-amyloid deposition through the brain: 0 = no amyloid; 1 = neocortical deposits only; 2 = allocortical involvement; 3 = subcortical nuclei; 4 = brainstem; 5 = cerebellar involvement. The five class boundaries are: no amyloid vs. any (0|1); neocortical only vs. wider spread (1|2); up to allocortical vs. subcortical-or-beyond (2|3); up to subcortical vs. brainstem-or-cerebellar (3|4); and brainstem vs. maximal cerebellar involvement (4|5).

Global performance. Globally, CSF Biomarkers (Rₛ = 0.666 ****) and Cognitive Composite Scores (Rₛ = 0.506 ****) led all feature sets, with Base Clinical also strong (Rₛ = 0.497 ****). All four sMRI subtypes reached global significance - rICV (Rₛ = 0.337 ****), Thickness (Rₛ = 0.305 ***), All sMRI (Rₛ = 0.241 **), and Volumes (Rₛ = 0.209 *) - as did WMH (Rₛ = 0.314 ****). Among DTI metrics, only DTI-AD approached significance globally (Rₛ = 0.295 *). No feature set significantly improved upon matched Base Clinical predictions globally.

Base Clinical (Rₛ = 0.497 ****): Consistently significant across all boundaries, rising from MCC = 0.201 **** at 0|1 to MCC = 0.456 **** at 4|5. Provided the reference signal throughout the staging spectrum.

sMRI – Volumes (Rₛ = 0.209 *): Non-significant at all boundaries except the highest (4|5: MCC = 0.209 *, ΔMCC = −0.300). The global significance reflects a concentration of signal at the most advanced stage of amyloid deposition.

sMRI – Thickness (Rₛ = 0.305 ***): Significant at the 1|2 (MCC = 0.282 *), 2|3 (MCC = 0.255 *), 3|4 (MCC = 0.208 *), and 4|5 (MCC = 0.279 ***) boundaries. The broadest coverage of any sMRI subtype, though never significantly exceeding matched Base Clinical predictions at any boundary.

sMRI – rICV (Rₛ = 0.337 ****): Non-significant at 0|1 through 3|4, reaching significance only at the highest boundary (4|5: MCC = 0.309 ***, ΔMCC = −0.200). Its global significance is carried almost entirely by that one boundary.

All sMRI Features (Rₛ = 0.241 **): Non-significant at all boundaries except 4|5 (MCC = 0.234 **, ΔMCC = −0.275). Like rICV, the global rank signal aggregates a very threshold-specific effect.

CSF Biomarkers (Rₛ = 0.666 ****): Non-significant at the earliest boundary (0|1: MCC = 0.065), then strongly significant from 1|2 onward: 1|2 (MCC = 0.396 ***), 2|3 (MCC = 0.515 ****), 3|4 (MCC = 0.590 ****), and 4|5 (MCC = 0.575 ****). Despite these large MCCs, no individual boundary improvement over Base Clinical survived FDR correction (maximum ΔMCC = 0.210 at 2|3).

White Matter Hyperintensities (Rₛ = 0.314 ****): Non-significant at the three lowest boundaries (0|1 through 2|3), then significant at 3|4 (MCC = 0.235 ***) and 4|5 (MCC = 0.296 ****). Signal appears only at advanced amyloid stages, consistent with co-occurring vascular-amyloid burden.

Cognitive Composite Scores (Rₛ = 0.506 ****): The only non-CSF feature set significant at all five boundaries: 0|1 (MCC = 0.208 ****), 1|2 (MCC = 0.270 ****), 2|3 (MCC = 0.268 ****), 3|4 (MCC = 0.349 ****), 4|5 (MCC = 0.469 ****). No boundary significantly exceeded matched Base Clinical predictions (maximum ΔMCC = 0.017 at 1|2).

DTI – Fractional Anisotropy (Rₛ = 0.122): Significantly negative at the earliest boundary (0|1: MCC = −0.179 ***), then non-significant at all remaining boundaries. The negative prediction at 0|1 indicates anti-correlated classification in the small DTI sub-cohort at this threshold.

DTI – Axial Diffusivity (Rₛ = 0.295 *): Non-significant at all individual boundaries despite a nominally positive global Rₛ. The global signal does not resolve into reliable predictions at any single threshold.

DTI – Mean Diffusivity (Rₛ = 0.156): Significantly negative at 0|1 (MCC = −0.137 **), non-significant at all remaining boundaries. Like DTI-FA, performs below chance for the earliest amyloid boundary.

DTI – Radial Diffusivity (Rₛ = 0.073): Non-significant globally and at all boundaries.

Summary. “Cognitive Composite Scores” was the only feature set with consistent significant predictions across all five Thal Phase boundaries. CSF Biomarkers provided the highest MCCs from stage 1|2 onward but failed entirely at the earliest threshold. sMRI signals were concentrated at the highest boundary (4|5), suggesting that macrostructural atrophy correlates most strongly with near-maximal amyloid burden. DTI metrics were counterproductive at the earliest boundary and unreliable throughout. No feature set significantly improved upon Base Clinical predictions at any single boundary after FDR correction.

**5.1.2 A Score  (Supplementary Figure 2)**

Staging. The A Score is a 4-class condensed summary of amyloid deposition derived from Thal Phase: 0 = no amyloid (Thal 0); 1 = mild (Thal 1–2); 2 = moderate (Thal 3); 3 = severe (Thal 4–5). Boundaries: no amyloid vs. any (0|1); mild vs. moderate-to-severe (1|2); mild-to-moderate vs. severe (2|3).

Global performance. CSF Biomarkers dominated globally (Rₛ = 0.683 ****) and significantly improved upon matched Base Clinical predictions globally (ΔMCC = 0.340 *). Cognitive Composite Scores (Rₛ = 0.358 ****) and Base Clinical (Rₛ = 0.343 ****) were next. WMH reached global significance (Rₛ = 0.232 ***); all sMRI subtypes and all DTI metrics were non-significant globally.

Base Clinical (Rₛ = 0.343 ****): Significant at all three boundaries — 0|1 (MCC = 0.185 ****), 1|2 (MCC = 0.262 ****), 2|3 (MCC = 0.334 ****) — with rising performance toward severe amyloid.

sMRI – Volumes (Rₛ = 0.080): Non-significant globally and at all boundaries; slightly negative at 0|1 (MCC = −0.053).

sMRI – Thickness (Rₛ = 0.141): Non-significant globally and at all boundaries, with nominally positive but sub-threshold values throughout.

sMRI – rICV (Rₛ = 0.085): Non-significant globally and at all boundaries.

All sMRI Features (Rₛ = 0.052): Non-significant globally and at all boundaries, with the weakest sMRI performance for this outcome.

CSF Biomarkers (Rₛ = 0.683 ****, ΔMCC = 0.340 *): Non-significant at 0|1 (MCC = 0.208), then strongly significant and increasingly dominant at 1|2 (MCC = 0.539 ****, ΔMCC = 0.266) and 2|3 (MCC = 0.672 ****, ΔMCC = 0.340 **). The 2|3 boundary is the only one where a feature set significantly improved upon matched Base Clinical predictions. CSF biomarkers appear uniquely capable of identifying the most severely affected individuals.

White Matter Hyperintensities (Rₛ = 0.232 ***): Non-significant at 0|1 (MCC = −0.007) and 1|2 (MCC = 0.147), then significant at 2|3 only (MCC = 0.239 ***, ΔMCC = −0.012). WMH signal appears exclusively at the severe amyloid boundary.

Cognitive Composite Scores (Rₛ = 0.358 ****): Significant at all three boundaries - 0|1 (MCC = 0.191 ****), 1|2 (MCC = 0.270 ****), 2|3 (MCC = 0.346 ****). No boundary significantly exceeded matched Base Clinical predictions (maximum ΔMCC = 0.014 at 2|3).

DTI – Fractional Anisotropy (Rₛ = 0.046): Non-significant globally and at all boundaries.

DTI – Axial Diffusivity (Rₛ = 0.223): Non-significant globally and at all boundaries, despite nominally rising values toward the severe end.

DTI – Mean Diffusivity (Rₛ = 0.173): Significantly negative at 0|1 (MCC = −0.122 **), then non-significant at higher boundaries. Anti-correlated prediction at the earliest threshold.

DTI – Radial Diffusivity (Rₛ = 0.129): Non-significant globally and at all boundaries.

Summary. A Score showed a clear two-tier structure: only clinical and cognitive data capture the none-vs-any boundary; CSF Biomarkers become decisive at moderate-to-severe thresholds, providing the only statistically significant improvement over Base Clinical in this outcome. sMRI and DTI contributed no reliable signal at any boundary. WMH showed a narrow signal at the severe boundary only.

**5.1.3 CERAD Score  (Supplementary Figure 3)**

Staging. The CERAD score is a 4-class measure of neuritic plaque density in the neocortex: 0 = none; 1 = sparse/possible; 2 = moderate/probable; 3 = frequent/definite. Boundaries: no plaques vs. any (0|1); sparse vs. moderate-or-definite (1|2); possible-to-probable vs. definite (2|3).

Global performance. CSF Biomarkers achieved the highest global Rₛ of any outcome in the entire study (Rₛ = 0.746 ****) and significantly improved upon matched Base Clinical (ΔMCC = 0.191 **). All four sMRI subtypes, WMH, and Cognitive Composite Scores were globally significant. DTI-AD was the only DTI metric to reach global significance (Rₛ = 0.160 *). Cognitive Composite Scores also significantly improved upon matched Base Clinical globally (ΔMCC = 0.020 *).

Base Clinical (Rₛ = 0.463 ****): Significant at all three boundaries: 0|1 (MCC = 0.321 ****), 1|2 (MCC = 0.366 ****), 2|3 (MCC = 0.427 ****). A strong and consistent reference throughout.

sMRI – Volumes (Rₛ = 0.243 ****): Non-significant at 0|1 (MCC = 0.078), marginally significant at 1|2 (MCC = 0.143 *), and clearly significant at 2|3 (MCC = 0.270 ****). Signal emerges only at the highest boundary.

sMRI – Thickness (Rₛ = 0.242 ****): Non-significant at 0|1 (MCC = 0.055) and 1|2 (MCC = 0.089), then significant at 2|3 (MCC = 0.225 ****, ΔMCC = −0.184). Like Volumes, meaningful only at the highest boundary.

sMRI – rICV (Rₛ = 0.295 ****): The strongest sMRI subtype for this outcome. Non-significant at 0|1 (MCC = 0.065), marginally significant at 1|2 (MCC = 0.136 *, ΔMCC = −0.214), and strongly significant at 2|3 (MCC = 0.332 ****, ΔMCC = −0.077).

All sMRI Features (Rₛ = 0.259 ****): Non-significant at 0|1 (MCC = 0.068) and 1|2 (MCC = 0.078), then significant at 2|3 (MCC = 0.266 ****, ΔMCC = −0.143).

CSF Biomarkers (Rₛ = 0.746 ****, ΔMCC = 0.191 **): Strongly significant at all three boundaries — 0|1 (MCC = 0.672 ****, ΔMCC = 0.258 *), 1|2 (MCC = 0.739 ****, ΔMCC = 0.184 *), 2|3 (MCC = 0.629 ****). The only feature set to significantly improve upon matched Base Clinical at the 0|1 and 1|2 boundaries. Unlike in Thal Phase or A Score, CSF Biomarkers here show strong signal even at the earliest threshold.

White Matter Hyperintensities (Rₛ = 0.339 ****): Significant at all three boundaries — 0|1 (MCC = 0.206 ****), 1|2 (MCC = 0.262 ****), 2|3 (MCC = 0.290 ****) — providing unusually uniform coverage without significantly exceeding Base Clinical at any boundary.

Cognitive Composite Scores (Rₛ = 0.479 ****, ΔMCC = 0.020 *): Significant at all three boundaries — 0|1 (MCC = 0.325 ****), 1|2 (MCC = 0.388 ****), 2|3 (MCC = 0.436 ****) — with no individually significant improvement over matched Base Clinical at any boundary despite the significant global increment.

DTI – Fractional Anisotropy (Rₛ = 0.150): Non-significant at 0|1 (MCC = 0.043) and 1|2 (MCC = 0.101), then marginally significant at 2|3 (MCC = 0.176 *, ΔMCC = −0.131).

DTI – Axial Diffusivity (Rₛ = 0.160 *): Non-significant at 0|1 (MCC = 0.009) and 1|2 (MCC = 0.108), then marginally significant at 2|3 (MCC = 0.197 *, ΔMCC = −0.110).

DTI – Mean Diffusivity (Rₛ = 0.134): Non-significant at all boundaries, including 2|3 (MCC = 0.135).

DTI – Radial Diffusivity (Rₛ = 0.121): Non-significant at 0|1 (MCC = −0.035) and 1|2 (MCC = 0.072), marginally significant at 2|3 (MCC = 0.175 *, ΔMCC = −0.132).

Summary. CERAD showed the broadest feature-set coverage of any amyloid outcome. CSF Biomarkers led at every boundary and were the only feature set to significantly exceed Base Clinical at the two lower thresholds, making them uniquely valuable for detecting any degree of neuritic plaque pathology. sMRI subtypes (particularly rICV) contributed meaningful signal but only at the highest boundary. WMH and Cognitive Composite Scores provided consistent, uniform coverage across all three boundaries. DTI metrics offered only marginal signal at the highest boundary.

**5.1.4 Braak Stage  (Supplementary Figure 4)**

Staging. The AD Braak Stage is a 7-class system (0–VI) characterizing the stereotypical spread of neurofibrillary tau pathology: 0 = none; I–II = transentorhinal (entorhinal cortex and hippocampus); III–IV = limbic (amygdala, cingulate); V–VI = neocortical. The six class boundaries are: no tau vs. transentorhinal onset (0|1); Braak I vs. II–VI (1|2); transentorhinal vs. limbic-or-greater (2|3); up to Stage III vs. IV–VI (3|4); up to Stage IV vs. V–VI (4|5); Braak V vs. maximal neocortical (5|6).

Global performance. CSF Biomarkers led (Rₛ = 0.742 ****) and significantly improved upon Base Clinical globally (ΔMCC = 0.158 **). Cognitive Composite Scores were second (Rₛ = 0.620 ****) and also significantly improved upon Base Clinical globally (ΔMCC = 0.047 ****). Base Clinical (Rₛ = 0.577 ****) was third. All sMRI subtypes and WMH were globally significant. Only DTI-MD reached global significance among DTI metrics (Rₛ = 0.160 *).

Base Clinical (Rₛ = 0.577 ****): Significant at all six boundaries, steadily rising: 0|1 (MCC = 0.189 ****), 1|2 (MCC = 0.274 ****), 2|3 (MCC = 0.314 ****), 3|4 (MCC = 0.410 ****), 4|5 (MCC = 0.516 ****), 5|6 (MCC = 0.517 ****).

sMRI – Volumes (Rₛ = 0.435 ****): Non-significant at 0|1 through 2|3; significant from 3|4 onward: 3|4 (MCC = 0.199 ***, ΔMCC = −0.236), 4|5 (MCC = 0.386 ****, ΔMCC = −0.128), 5|6 (MCC = 0.441 ****, ΔMCC = −0.151). Reliable signal only at limbic and neocortical stages.

sMRI – Thickness (Rₛ = 0.429 ****): Paradoxically negative at 0|1 (MCC = −0.016 **) and 1|2 (MCC = −0.030 ***), then non-significant at 2|3, recovering to significance at 3|4 (MCC = 0.182 **), 4|5 (MCC = 0.438 ****), and 5|6 (MCC = 0.501 ****). The anti-correlated predictions at early tau stages are a notable artifact of attempting to classify very early tau accumulation from macrostructural features in a small sub-cohort.

sMRI – rICV (Rₛ = 0.502 ****): Non-significant at 0|1 through 2|3; significant at 3|4 (MCC = 0.200 ***, ΔMCC = −0.235), 4|5 (MCC = 0.467 ****, ΔMCC = −0.046), and 5|6 (MCC = 0.506 ****, ΔMCC = −0.086). The strongest sMRI subtype for Braak Stage globally, concentrated in the highest boundaries.

All sMRI Features (Rₛ = 0.450 ****): Significantly negative at 0|1 (MCC = −0.013 *) and 1|2 (MCC = −0.027 ***); non-significant at 2|3; then significant at 3|4 (MCC = 0.236 ****, ΔMCC = −0.198), 4|5 (MCC = 0.434 ****, ΔMCC = −0.079), 5|6 (MCC = 0.460 ****, ΔMCC = −0.132). Mirrors the pattern of Thickness with early-stage artifacts.

CSF Biomarkers (Rₛ = 0.742 ****, ΔMCC = 0.158 **): Insufficient class representation at 0|1 (NA); non-significant at 1|2 (MCC = 0.123); then strongly significant from 2|3 onward: 2|3 (MCC = 0.500 ****), 3|4 (MCC = 0.642 ****), 4|5 (MCC = 0.658 ****), 5|6 (MCC = 0.670 ****, ΔMCC = 0.246 **). Significantly exceeds matched Base Clinical predictions at the highest boundary only.

White Matter Hyperintensities (Rₛ = 0.424 ****): Non-significant at 0|1 through 2|3; marginally significant at 3|4 (MCC = 0.112 *, ΔMCC = −0.240); strongly significant at 4|5 (MCC = 0.427 ****, ΔMCC = −0.073) and 5|6 (MCC = 0.476 ****, ΔMCC = −0.054). Signal confined to the highest tau stages.

Cognitive Composite Scores (Rₛ = 0.620 ****, ΔMCC = 0.047 ****): The only non-CSF feature set significant at all six boundaries: 0|1 (MCC = 0.180 ****), 1|2 (MCC = 0.290 ****), 2|3 (MCC = 0.350 ****), 3|4 (MCC = 0.424 ****), 4|5 (MCC = 0.562 ****, ΔMCC = 0.054 ****), 5|6 (MCC = 0.573 ****, ΔMCC = 0.054 ***). Significantly improved upon matched Base Clinical predictions at the two highest boundaries — the most consistent incremental contribution of any feature set for this outcome.

DTI – Fractional Anisotropy (Rₛ = 0.092): Insufficient class representation at the two lowest boundaries (0|1, 1|2: NA); non-significant at all remaining boundaries. No useful signal for Braak Stage.

DTI – Axial Diffusivity (Rₛ = 0.132): Insufficient class representation at 0|1 (NA); nominally positive but non-significant at all remaining boundaries.

DTI – Mean Diffusivity (Rₛ = 0.160 *): Insufficient class representation at 0|1 (NA); non-significant at 1|2 through 3|4 and 5|6; marginally significant at 4|5 only (MCC = 0.227 **, ΔMCC = −0.160).

DTI – Radial Diffusivity (Rₛ = 0.132): Insufficient class representation at early boundaries; non-significant throughout.

Summary. Braak Stage showed the clearest gradient of any outcome: early tau boundaries (0|1, 1|2) are not only poorly predicted by sMRI but actively anti-correlated for Thickness and All sMRI Features. From the limbic boundary (2|3 and 3|4) onward, sMRI, WMH, and CSF Biomarkers all become reliable. Cognitive Composite Scores was uniquely consistent — the only feature set with significant predictions at every boundary — and the only one to significantly exceed Base Clinical predictions at the highest two boundaries. CSF Biomarkers provided the largest absolute MCCs from 2|3 onward and significantly beat Base Clinical at the 5|6 boundary. DTI metrics were largely unreliable throughout.

**5.1.5 B Score  (Supplementary Figure 5)**

Staging. The B Score is a 4-class condensed summary of tau neurofibrillary pathology derived from AD Braak Stage: 0 = no tau (Braak 0); 1 = low burden (Braak I–II); 2 = intermediate burden (Braak III–IV); 3 = high burden (Braak V–VI). Boundaries: no tau vs. any (0|1); low vs. intermediate-to-high (1|2); low-to-intermediate vs. high (2|3).

Global performance. CSF Biomarkers led (Rₛ = 0.678 ****), followed by Cognitive Composite Scores (Rₛ = 0.559 ****, ΔMCC = 0.050 ****), Base Clinical (Rₛ = 0.515 ****), WMH (Rₛ = 0.419 ****), and all four sMRI subtypes (0.350–0.370, all ****). All four DTI metrics reached global significance (FA 0.236 **, AD 0.246 **, MD 0.259 ***, RD 0.216 **). Cognitive Composite Scores was the only non-CSF feature set to significantly improve upon matched Base Clinical globally.

Base Clinical (Rₛ = 0.515 ****): Significant at all three boundaries, rising from 0|1 (MCC = 0.157 ***) to 1|2 (MCC = 0.306 ****) to 2|3 (MCC = 0.517 ****).

sMRI – Volumes (Rₛ = 0.360 ****): Significantly negative at 0|1 (MCC = −0.013 *); non-significant at 1|2 (MCC = 0.121); strongly significant at 2|3 (MCC = 0.419 ****, ΔMCC = −0.081). All sMRI signal concentrated at the highest tau boundary.

sMRI – Thickness (Rₛ = 0.364 ****): Non-significant at 0|1 (MCC = 0.299) and 1|2 (MCC = 0.074, note the non-monotone pattern); strongly significant at 2|3 (MCC = 0.417 ****, ΔMCC = −0.082). Note that MCC at 0|1 is nominally high but not significant, likely due to variance in the sub-cohort.

sMRI – rICV (Rₛ = 0.350 ****): Non-significant at 0|1 (MCC = 0.263) and 1|2 (MCC = 0.036); significant at 2|3 (MCC = 0.408 ****, ΔMCC = −0.091).

All sMRI Features (Rₛ = 0.370 ****): Significantly negative at 0|1 (MCC = −0.013 *); non-significant at 1|2 (MCC = 0.143); significant at 2|3 (MCC = 0.399 ****, ΔMCC = −0.101).

CSF Biomarkers (Rₛ = 0.678 ****): Insufficient class representation at 0|1 (NA); strongly significant at 1|2 (MCC = 0.528 ****) and 2|3 (MCC = 0.650 ****). No individually significant improvement over matched Base Clinical at either available boundary.

White Matter Hyperintensities (Rₛ = 0.419 ****): Non-significant at 0|1 (MCC = −0.005) and 1|2 (MCC = 0.101); strongly significant at 2|3 (MCC = 0.464 ****, ΔMCC = −0.017). Consistent with Braak Stage, WMH signal is limited to the highest tau stage.

Cognitive Composite Scores (Rₛ = 0.559 ****, ΔMCC = 0.050 ****): Significant at all three boundaries — 0|1 (MCC = 0.159 ***), 1|2 (MCC = 0.356 ****), 2|3 (MCC = 0.567 ****, ΔMCC = 0.058 ****). Significantly improved upon matched Base Clinical predictions at the highest boundary — the clearest incremental gain over Base Clinical for any tau outcome at a single class boundary.

DTI – Fractional Anisotropy (Rₛ = 0.236 **): Insufficient class representation at 0|1 (NA); non-significant at 1|2 (MCC = 0.115); significant at 2|3 only (MCC = 0.242 **, ΔMCC = −0.134).

DTI – Axial Diffusivity (Rₛ = 0.246 **): Insufficient class representation at 0|1 (NA); non-significant at 1|2 (MCC = 0.152); significant at 2|3 only (MCC = 0.220 **, ΔMCC = −0.155).

DTI – Mean Diffusivity (Rₛ = 0.259 ***): Non-significant at 0|1 (MCC = −0.011) and 1|2 (MCC = 0.161); significant at 2|3 only (MCC = 0.222 **, ΔMCC = −0.154).

DTI – Radial Diffusivity (Rₛ = 0.216 **): Insufficient class representation at 0|1 (NA); non-significant at 1|2 (MCC = 0.107); significant at 2|3 only (MCC = 0.216 *, ΔMCC = −0.159).

Summary. B Score produced the most uniformly significant results of any single boundary in the entire study at the 2|3 threshold: every available feature set reached significance, including all four DTI metrics — an occurrence not observed at any other boundary across all outcomes. This concentration of signal at the high-tau boundary reflects the cumulative structural, microstructural, and cognitive impact of advanced neurofibrillary disease. Cognitive Composite Scores was the only feature set to significantly improve upon Base Clinical at any boundary. sMRI and DTI were again absent at the earliest tau boundary.

**5.1.6 CVD — Arteriolosclerosis  (Supplementary Figure 6)**

Staging. Arteriolosclerosis is a 4-class measure of pathological thickening and hyalinization of small arterioles: 0 = none; 1 = mild; 2 = moderate; 3 = severe. Boundaries: none vs. any (0|1); mild vs. moderate-to-severe (1|2); mild-to-moderate vs. severe (2|3).

Global performance. DTI diffusivity metrics were the strongest global predictors: DTI-MD (Rₛ = 0.360 ****), DTI-AD (Rₛ = 0.302 ****), and DTI-RD (Rₛ = 0.298 ****). CSF Biomarkers (Rₛ = 0.269 ****), WMH (Rₛ = 0.228 ****), and Cognitive Composite Scores (Rₛ = 0.214 ****) followed. Cortical Thickness (Rₛ = 0.147 **), sMRI Volumes (Rₛ = 0.127 *), and All sMRI Features (Rₛ = 0.119 *) were marginally globally significant; rICV and DTI-FA were not. Cognitive Composite Scores significantly improved upon matched Base Clinical globally (ΔMCC = 0.081 ****).

Base Clinical (Rₛ = 0.135 ****): Significant at 0|1 (MCC = 0.067 ****) and 1|2 (MCC = 0.119 ****), but not at 2|3 (MCC = 0.006). Base Clinical signal collapses at the severe boundary.

sMRI – Volumes (Rₛ = 0.127 *): Non-significant at 0|1 (MCC = 0.046) and 2|3 (MCC = −0.003); marginally significant at 1|2 only (MCC = 0.137 *, ΔMCC = 0.038).

sMRI – Thickness (Rₛ = 0.147 **): Non-significant at 0|1 (MCC = 0.101); significant at 1|2 (MCC = 0.125 *, ΔMCC = 0.026); significantly negative at 2|3 (MCC = −0.027 *). The negative prediction at the severe boundary indicates anti-correlated model behavior.

sMRI – rICV (Rₛ = 0.062): Non-significant globally and at all boundaries; slightly negative at 0|1.

All sMRI Features (Rₛ = 0.119 *): Non-significant at 0|1 (MCC = −0.009); significant at 1|2 (MCC = 0.134 *, ΔMCC = 0.035); significantly negative at 2|3 (MCC = −0.046 ****). Same problematic pattern as Thickness at the severe boundary.

CSF Biomarkers (Rₛ = 0.269 ****): Dominated at the 0|1 boundary (MCC = 0.489 ****, ΔMCC = 0.391 ***) — the largest single-boundary incremental improvement over Base Clinical of any feature set for any outcome in the study. Dropped to marginal significance at 1|2 (MCC = 0.142 *) and non-significant at 2|3 (MCC = 0.024). The CSF signal is uniquely concentrated at the none-vs-any threshold.

White Matter Hyperintensities (Rₛ = 0.228 ****): Significant at 0|1 (MCC = 0.132 **) and 1|2 (MCC = 0.196 ****); non-significant at 2|3 (MCC = 0.004). Reasonably consistent across the two lower boundaries.

Cognitive Composite Scores (Rₛ = 0.214 ****, ΔMCC = 0.081 ****): Significant at 0|1 (MCC = 0.161 ****, ΔMCC = 0.097 ****) and 1|2 (MCC = 0.167 ****, ΔMCC = 0.049 *); marginally significant at 2|3 (MCC = 0.052 *, ΔMCC = 0.043). Significantly improved upon matched Base Clinical predictions at both the 0|1 and 1|2 boundaries — the most consistent incremental contribution for this outcome.

DTI – Fractional Anisotropy (Rₛ = 0.093): Non-significant at 0|1 (MCC = 0.115) and 1|2; significantly negative at 2|3 (MCC = −0.030 *).

DTI – Axial Diffusivity (Rₛ = 0.302 ****): Strongly significant at 0|1 (MCC = 0.299 ***, ΔMCC = 0.177); non-significant at 1|2 (MCC = 0.079) and 2|3. Signal concentrated entirely at the none-vs-any boundary.

DTI – Mean Diffusivity (Rₛ = 0.360 ****): Significant at both 0|1 (MCC = 0.306 ***, ΔMCC = 0.184) and 1|2 (MCC = 0.212 **, ΔMCC = 0.039); non-significant at 2|3. The broadest DTI coverage for this outcome.

DTI – Radial Diffusivity (Rₛ = 0.298 ****): Significant at 0|1 (MCC = 0.262 **, ΔMCC = 0.140) and 1|2 (MCC = 0.191 *, ΔMCC = 0.018); non-significant at 2|3.

Summary. Arteriolosclerosis had a distinct profile: DTI diffusivity and CSF Biomarkers led at the none-vs-mild boundary, with CSF providing the single largest incremental gain over Base Clinical of any feature set and any outcome in the study. DTI-MD and DTI-RD also covered the 1|2 boundary. sMRI subtypes contributed only modestly at 1|2, and were unreliable or negative at the severe end. The severe boundary (2|3) was essentially unpredictable by all feature sets, consistent with its rarity and pathological heterogeneity.

**5.1.7 CVD — Atherosclerosis  (Supplementary Figure 7)**

Staging. Atherosclerosis is a 4-class measure of large-vessel arterial disease: 0 = none; 1 = mild; 2 = moderate; 3 = severe. Boundaries: none vs. any (0|1); mild vs. moderate-to-severe (1|2); mild-to-moderate vs. severe (2|3).

Global performance. DTI-AD was the strongest single predictor globally (Rₛ = 0.262 ***). Nearly all feature sets reached global significance: Cognitive Composite Scores (Rₛ = 0.235 ****, ΔMCC = 0.036 *), rICV (Rₛ = 0.223 ****), Thickness (Rₛ = 0.207 ****), Base Clinical (Rₛ = 0.206 ****), DTI-MD (Rₛ = 0.208 **), CSF Biomarkers (Rₛ = 0.178 **), WMH (Rₛ = 0.147 ***), All sMRI Features (Rₛ = 0.162 **), DTI-RD (Rₛ = 0.181 *), Volumes (Rₛ = 0.132 *). DTI-FA was non-significant globally. Cognitive Composite Scores was the only feature set to significantly improve upon matched Base Clinical globally.

Base Clinical (Rₛ = 0.206 ****): Significant at 0|1 (MCC = 0.217 ****) and 1|2 (MCC = 0.117 ****); non-significant at 2|3 (MCC = 0.025).

sMRI – Volumes (Rₛ = 0.132 *): Non-significant at 0|1 (MCC = 0.039); marginally significant at 1|2 (MCC = 0.138 *); non-significant at 2|3 (MCC = 0.190, note NA for base comparison indicating sparse class representation).

sMRI – Thickness (Rₛ = 0.207 ****): Significant at 0|1 (MCC = 0.205 **) and 1|2 (MCC = 0.137 *); significantly negative at 2|3 (MCC = −0.018 *).

sMRI – rICV (Rₛ = 0.223 ****): Significant at 0|1 (MCC = 0.166 *) and 1|2 (MCC = 0.150 *); NA at 2|3 due to absent minority class predictions.

All sMRI Features (Rₛ = 0.162 **): Marginally significant at 0|1 (MCC = 0.144 *); non-significant at 1|2 and 2|3.

CSF Biomarkers (Rₛ = 0.178 **): Significantly negative at 0|1 (MCC = −0.068 ****), indicating an anti-correlated prediction for the none-vs-any boundary — the CSF amyloid-tau profile appears paradoxically inversely related to large-vessel atherosclerosis onset. Significant and positive at 1|2 (MCC = 0.232 ***); non-significant at 2|3.

White Matter Hyperintensities (Rₛ = 0.147 ***): Significant at 0|1 (MCC = 0.146 **) and 1|2 (MCC = 0.124 **); non-significant at 2|3 (MCC = 0.167).

Cognitive Composite Scores (Rₛ = 0.235 ****, ΔMCC = 0.036 *): Significant at 0|1 (MCC = 0.201 ****) and 1|2 (MCC = 0.165 ****); non-significant at 2|3. No individually significant boundary improvement over matched Base Clinical.

DTI – Fractional Anisotropy (Rₛ = 0.123): Non-significant at 0|1 and 1|2; significantly negative at 2|3 (MCC = −0.033 ***).

DTI – Axial Diffusivity (Rₛ = 0.262 ***): Strongly significant at 0|1 (MCC = 0.291 **, ΔMCC = 0.242); non-significant at 1|2 and 2|3 (negative at 2|3). Signal concentrated entirely at the none-vs-any boundary.

DTI – Mean Diffusivity (Rₛ = 0.208 **): Non-significant at 0|1; significant at 1|2 (MCC = 0.182 *); non-significant at 2|3.

DTI – Radial Diffusivity (Rₛ = 0.181 *): Non-significant at all boundaries individually.

Summary. Atherosclerosis was distinctive for the paradoxically negative CSF prediction at the none-vs-any boundary, likely reflecting divergent risk factor profiles for AD-type and large-vessel pathologies. DTI-AD and most sMRI subtypes showed moderate signals at the lower boundaries. The severe boundary was universally unpredictable. Cognitive Composite Scores provided the most consistent coverage across both accessible boundaries.

**5.1.8 CVD — Cerebral Amyloid Angiopathy  (Supplementary Figure 8)**

Staging. CAA is a 4-class measure of β-amyloid deposition in cerebral vessel walls: 0 = none; 1 = mild; 2 = moderate; 3 = severe. Boundaries: none vs. any (0|1); mild vs. moderate-to-severe (1|2); mild-to-moderate vs. severe (2|3).

Global performance. Only CSF Biomarkers (Rₛ = 0.382 ****), Cognitive Composite Scores (Rₛ = 0.264 ****, ΔMCC = 0.078 ****), and Base Clinical (Rₛ = 0.186 ****) reached global significance. All sMRI subtypes, WMH, and all DTI metrics were globally non-significant. Cognitive Composite Scores significantly improved upon matched Base Clinical globally.

Base Clinical (Rₛ = 0.186 ****): Significant at 0|1 (MCC = 0.180 ****) and 1|2 (MCC = 0.044 ***); non-significant at 2|3 (MCC = 0.017).

sMRI – Volumes (Rₛ = 0.063): Non-significant at all boundaries.

sMRI – Thickness (Rₛ = 0.083): Non-significant at 0|1 and 1|2; significantly negative at 2|3 (MCC = −0.093 **).

sMRI – rICV (Rₛ = 0.080): Non-significant at all boundaries.

All sMRI Features (Rₛ = −0.028): Non-significant globally; slightly negative at all boundaries.

CSF Biomarkers (Rₛ = 0.382 ****): Significant at 0|1 (MCC = 0.253 ***) and strongly significant at 1|2 (MCC = 0.333 ****); non-significant at 2|3 (MCC = 0.123). Consistent positive signal at the two lower boundaries.

White Matter Hyperintensities (Rₛ = 0.043): Non-significant globally and at all boundaries.

Cognitive Composite Scores (Rₛ = 0.264 ****, ΔMCC = 0.078 ****): Significant at 0|1 (MCC = 0.255 ****, ΔMCC = 0.077 ****) and 1|2 (MCC = 0.108 ****, ΔMCC = 0.060 *); non-significant at 2|3. Significantly improved upon matched Base Clinical predictions at both the 0|1 and 1|2 boundaries — the most consistent incremental contribution for CAA.

DTI – Fractional Anisotropy (Rₛ = −0.061): Non-significant globally; significantly negative at 2|3 (MCC = −0.069 ****).

DTI – Axial Diffusivity (Rₛ = 0.078): Non-significant at all boundaries.

DTI – Mean Diffusivity (Rₛ = 0.114): Non-significant at all boundaries.

DTI – Radial Diffusivity (Rₛ = 0.062): Non-significant globally; significantly negative at 2|3 (MCC = −0.045 **).

Summary. CAA was predicted almost exclusively by CSF Biomarkers and Cognitive Composite Scores at the two lower boundaries; all structural and diffusion imaging feature sets failed to reach significance at any boundary. Cognitive Composite Scores significantly exceeded matched Base Clinical at both accessible boundaries. The severe boundary (2|3) was unpredictable, with several imaging feature sets producing significantly negative predictions.

**5.1.9 PD Braak Stage (5-class Lewy Body)  (Supplementary Figure 9)**

Staging. The PD Braak Stage is a 5-class system characterizing α-synuclein/Lewy body spread: 0 = none; 1 = brainstem-predominant; 2 = limbic/amygdala-predominant; 3 = neocortical; 4 = olfactory-bulb-predominant. Boundaries: none vs. any (0|1); brainstem vs. limbic-or-greater (1|2); brainstem-to-limbic vs. neocortical-or-olfactory (2|3); neocortical vs. olfactory-predominant (3|4).

Global performance. Only CSF Biomarkers (Rₛ = 0.142 *), Cognitive Composite Scores (Rₛ = 0.102 ****), and Base Clinical (Rₛ = 0.067 ****) reached global significance. All sMRI subtypes, WMH, and all DTI metrics were non-significant globally. No feature set significantly improved upon matched Base Clinical globally.

Base Clinical (Rₛ = 0.067 ****): Significant at 0|1 (MCC = 0.071 ****) and 1|2 (MCC = 0.080 ****); non-significant at 2|3 (MCC = 0.024) and 3|4 (MCC = 0.004).

sMRI – Volumes (Rₛ = −0.005): Non-significant globally and at all boundaries; NA at 3|4. No useful signal.

sMRI – Thickness (Rₛ = 0.055): Non-significant at all boundaries; NA at 3|4.

sMRI – rICV (Rₛ = 0.027): Non-significant at all boundaries; NA at 3|4.

All sMRI Features (Rₛ = 0.036): Non-significant at all boundaries; NA at 3|4.

CSF Biomarkers (Rₛ = 0.142 *): Significant at 0|1 (MCC = 0.155 *) and 1|2 (MCC = 0.202 **); non-significant at 2|3 (MCC = 0.024); NA at 3|4. The only imaging-adjacent feature set with any class-level signal.

White Matter Hyperintensities (Rₛ = 0.027): Non-significant at all boundaries; NA at 3|4.

Cognitive Composite Scores (Rₛ = 0.102 ****): Significant at 0|1 (MCC = 0.102 ****), 1|2 (MCC = 0.115 ****), and 2|3 (MCC = 0.082 ****); marginally significant and effectively at chance at 3|4 (MCC = −0.007 ****). The only feature set with any positive signal at the 2|3 boundary; also the only feature set with data at 3|4.

DTI – Fractional Anisotropy (Rₛ = 0.046): Non-significant at 0|1 through 2|3; NA at 3|4.

DTI – Axial, Mean, Radial Diffusivity: All non-significant at all available boundaries; NA at 3|4.

Summary. Fine-grained PD Braak staging was nearly entirely unpredictable across all feature sets. Only CSF Biomarkers and Cognitive Composite Scores showed any meaningful class-level signal, and only at the two lowest boundaries. The 3|4 boundary was effectively inaccessible to most feature sets due to sparse class representation. DTI and sMRI contributed nothing. The progressive collapse of predictability across boundaries illustrates the fundamental difficulty of staging Lewy body pathology from antemortem data with currently available feature sets.

**5.1.10 L Score (3-class Lewy Body)  (Supplementary Figure 10)**

Staging. The L Score is a 3-class condensed summary of Lewy body pathology derived from PD Braak staging: 0 = none (PD Braak 0); 1 = brainstem-predominant (PD Braak Stage 1); 2 = limbic or neocortical involvement (PD Braak Stages 2–4). Boundaries: no Lewy bodies vs. any (0|1); brainstem-predominant vs. limbic-or-neocortical (1|2).

Global performance. CSF Biomarkers led (Rₛ = 0.220 ***), followed by Cognitive Composite Scores (Rₛ = 0.152 ****), WMH (Rₛ = 0.128 **), Base Clinical (Rₛ = 0.121 ****), rICV (Rₛ = 0.126 *), All sMRI Features (Rₛ = 0.124 *), and DTI-FA (Rₛ = 0.175 *). sMRI Volumes and Thickness were non-significant globally. No feature set significantly improved upon matched Base Clinical globally.

Base Clinical (Rₛ = 0.121 ****): Significant at both boundaries — 0|1 (MCC = 0.114 ****) and 1|2 (MCC = 0.128 ****) — with modest and comparable MCCs.

sMRI – Volumes (Rₛ = 0.064): Non-significant at both boundaries.

sMRI – Thickness (Rₛ = 0.088): Non-significant at both boundaries.

sMRI – rICV (Rₛ = 0.126 *): Marginally significant at 0|1 (MCC = 0.130 *); non-significant at 1|2 (MCC = 0.108).

All sMRI Features (Rₛ = 0.124 *): Marginally significant at 0|1 (MCC = 0.135 *); non-significant at 1|2 (MCC = 0.092).

CSF Biomarkers (Rₛ = 0.220 ***): Significant at 0|1 (MCC = 0.203 **) and 1|2 (MCC = 0.243 ***). The strongest feature set for both boundaries.

White Matter Hyperintensities (Rₛ = 0.128 **): Significant at 0|1 (MCC = 0.131 **) and 1|2 (MCC = 0.114 *). Consistent modest coverage at both boundaries.

Cognitive Composite Scores (Rₛ = 0.152 ****): Significant at 0|1 (MCC = 0.147 ****) and 1|2 (MCC = 0.155 ****). Consistent modest coverage.

DTI – Fractional Anisotropy (Rₛ = 0.175 *): Non-significant at either boundary individually (0|1: MCC = 0.169; 1|2: MCC = 0.188), despite reaching global significance — a case where the global Rₛ aggregates a consistent but sub-threshold signal.

DTI – Axial Diffusivity (Rₛ = 0.067): Non-significant at either boundary.

DTI – Mean Diffusivity (Rₛ = 0.139): Non-significant at either boundary.

DTI – Radial Diffusivity (Rₛ = 0.149): Non-significant at either boundary.

Summary. L Score showed modest but broadly distributed predictability across feature sets. CSF Biomarkers provided the strongest and most consistent signal at both boundaries. WMH, Cognitive Composite Scores, and rICV contributed at the lower boundary. No feature set significantly improved upon Base Clinical at either boundary, consistent with the overall difficulty of Lewy body prediction.

**5.1.11 TDP-43 5-Stage  (Supplementary Figure 11)**

Staging. The TDP-43 5-Stage is a 6-class system tracking the anatomical progression of TDP-43 inclusions: 0 = none; 1 = brainstem/spinal cord; 2 = amygdala; 3 = hippocampus; 4 = entorhinal cortex; 5 = neocortex. Boundaries: none vs. any (0|1); Stage 1 vs. 2–5 (1|2); Stages 0–2 vs. 3–5 (2|3); Stages 0–3 vs. 4–5 (3|4); Stage 4 vs. 5 (4|5).

Global performance. Only Cognitive Composite Scores reached global significance (Rₛ = 0.140 ****, ΔMCC = 0.094 *) and significantly improved upon matched Base Clinical globally. Base Clinical was only marginally significant (Rₛ = 0.046 *). All sMRI subtypes, CSF Biomarkers, WMH, and all DTI metrics were globally non-significant.

Base Clinical (Rₛ = 0.046 *): Non-significant at 0|1 (MCC = 0.034) and 1|2 (MCC = 0.039); marginally significant at 2|3 (MCC = 0.054 *) and 3|4 (MCC = 0.073 **); non-significant at 4|5 (MCC = 0.077).

sMRI – Volumes (Rₛ = −0.129): Non-significant globally; significantly negative at 3|4 (MCC = −0.150 ****). Consistently negative values throughout.

sMRI – Thickness (Rₛ = −0.009): Non-significant at all boundaries; NA at 4|5.

sMRI – rICV (Rₛ = 0.097): Non-significant globally and at all boundaries, though nominally positive throughout.

All sMRI Features (Rₛ = 0.120): Non-significant globally; significantly negative at 2|3 (MCC = −0.130 ***) and 3|4 (MCC = −0.112 ***). Anti-correlated predictions at intermediate boundaries.

CSF Biomarkers (Rₛ = 0.160): Non-significant globally; significantly negative at 4|5 only (MCC = −0.038 *). Model behavior at the highest boundary is unreliable.

White Matter Hyperintensities (Rₛ = −0.059): Non-significant globally; significantly negative at 4|5 (MCC = −0.029 **).

Cognitive Composite Scores (Rₛ = 0.140 ****, ΔMCC = 0.094 *): Significant at 0|1 (MCC = 0.130 ****), 1|2 (MCC = 0.133 ****), 2|3 (MCC = 0.144 ****), and 3|4 (MCC = 0.140 ****); non-significant at 4|5 (MCC = 0.079). The only feature set with any positive signal at any boundary, and the only one to significantly improve upon matched Base Clinical globally.

DTI – Fractional Anisotropy (Rₛ = 0.101): Non-significant at 0|1 and 1|2; NA at 2|3, 3|4, and 4|5 (insufficient class representation).

DTI – Axial Diffusivity (Rₛ = −0.277): Non-significant at 0|1 and 1|2; significantly negative at 2|3 (MCC = −0.293 ****) and 3|4 (MCC = −0.279 ****). Strongly anti-correlated predictions at intermediate stages.

DTI – Mean Diffusivity (Rₛ = −0.000): Non-significant at all boundaries; significantly negative at 4|5 (MCC = −0.086 *).

DTI – Radial Diffusivity (Rₛ = −0.007): Non-significant at all boundaries; significantly negative at 4|5 (MCC = −0.101 **).

Summary. TDP-43 5-Stage was the most uniformly poor outcome in the study. Only Cognitive Composite Scores provided any positive signal, and exclusively at the four lower boundaries. The pattern of significantly negative MCCs from sMRI (Volumes, All sMRI) and DTI-AD at intermediate boundaries reflects model instability from inadequate sample sizes and sparse class occupancy rather than true informative anti-correlation. This outcome serves as a cautionary example of what can occur when ordinal classes are too fine-grained relative to sample size.

**5.1.12 TDP-43 3-Stage  (Supplementary Figure 12)**

Staging. The TDP-43 3-Stage is a 4-class condensed summary of LATE-NC (limbic-predominant age-related TDP-43 encephalopathy): 0 = none; 1 = brainstem/spinal cord involvement; 2 = amygdala involvement; 3 = hippocampal, entorhinal cortex, and/or neocortical involvement. Boundaries: none vs. any TDP-43 (0|1); brainstem vs. amygdala-or-greater (1|2); brainstem-to-amygdala vs. widespread (2|3).

Global performance. Cognitive Composite Scores led globally (Rₛ = 0.263 ****, ΔMCC = 0.070 **) and significantly improved upon matched Base Clinical globally. Base Clinical (Rₛ = 0.193 ****), rICV (Rₛ = 0.217 ***), Volumes (Rₛ = 0.175 **), Thickness (Rₛ = 0.139 *), and CSF Biomarkers (Rₛ = 0.189 *) all reached global significance. WMH, All sMRI Features, and all DTI metrics were globally non-significant.

Base Clinical (Rₛ = 0.193 ****): Significant and stable across all three boundaries: 0|1 (MCC = 0.167 ****), 1|2 (MCC = 0.167 ****), 2|3 (MCC = 0.207 ****).

sMRI – Volumes (Rₛ = 0.175 **): Significant at 0|1 (MCC = 0.159 *) and 1|2 (MCC = 0.207 **); non-significant at 2|3 (MCC = 0.081). Signal strongest at the brainstem-to-amygdala boundary.

sMRI – Thickness (Rₛ = 0.139 *): Non-significant at 0|1 (MCC = 0.082); marginally significant at 1|2 (MCC = 0.144 *); significant at 2|3 (MCC = 0.229 **). Signal shifts to the higher boundary.

sMRI – rICV (Rₛ = 0.217 ***): Significant at all three boundaries — 0|1 (MCC = 0.158 *), 1|2 (MCC = 0.218 **), 2|3 (MCC = 0.220 **). The most broadly covered sMRI subtype for TDP-43 3-Stage.

All sMRI Features (Rₛ = 0.085): Non-significant globally and at all boundaries. Combining all sMRI measures does not help for this outcome.

CSF Biomarkers (Rₛ = 0.189 *): Non-significant at any individual boundary — 0|1 (MCC = 0.158), 1|2 (MCC = 0.209), 2|3 (MCC = 0.186) — despite global significance. Values approach significance at all boundaries but none survive FDR correction.

White Matter Hyperintensities (Rₛ = 0.049): Non-significant globally and at all boundaries.

Cognitive Composite Scores (Rₛ = 0.263 ****, ΔMCC = 0.070 **): Significant at all three boundaries — 0|1 (MCC = 0.228 ****), 1|2 (MCC = 0.237 ****, ΔMCC = 0.073 *), 2|3 (MCC = 0.275 ****). Significantly improved upon matched Base Clinical at the 1|2 boundary specifically.

DTI – Fractional Anisotropy (Rₛ = −0.084): Non-significant globally; consistently negative across all boundaries.

DTI – Axial Diffusivity (Rₛ = −0.015): Non-significant globally; consistently negative across all boundaries.

DTI – Mean Diffusivity (Rₛ = 0.128): Non-significant globally and at all boundaries.

DTI – Radial Diffusivity (Rₛ = −0.035): Non-significant globally; consistently negative across all boundaries.

Summary. TDP-43 3-Stage was notable for the consistent failure of DTI metrics (several showing negative global Rₛ) and the strong performance of Cognitive Composite Scores, which was the only feature set to significantly improve upon Base Clinical at a class boundary. rICV was the most useful sMRI subtype, reaching significance at all three boundaries. CSF Biomarkers had a suggestive but sub-threshold pattern. The outcome is one of the few where cognitive rather than imaging biomarkers carry the most discriminative weight.

**5.1.13 White Matter Rarefaction (Supplementary Figure 13)**

Staging. White matter rarefaction (WMR) is a 4-class measure of diffuse white matter pallor reflecting axonal and myelin injury from vascular or neurodegenerative causes: 0 = none; 1 = mild; 2 = moderate; 3 = severe. Boundaries: none vs. any rarefaction (0|1); mild vs. moderate-to-severe (1|2); mild-to-moderate vs. severe (2|3).

Global performance. DTI diffusivity metrics were the strongest global predictors: DTI-RD (Rₛ = 0.363 *), DTI-MD (Rₛ = 0.344 *), DTI-AD (Rₛ = 0.339 *). All sMRI Features was also strongly significant globally (Rₛ = 0.337 ***). WMH (Rₛ = 0.199 **) and Cognitive Composite Scores (Rₛ = 0.131 ****, ΔMCC = 0.111 ****) also reached global significance. Base Clinical was non-significant globally (Rₛ = 0.016) — one of only two outcomes in the main text where Base Clinical failed to reach significance. Individual sMRI subtypes (Volumes, Thickness, rICV) and DTI-FA were all globally non-significant.

Base Clinical (Rₛ = 0.016): Marginally significant at 0|1 only (MCC = 0.045 *); non-significant at 1|2 (MCC = 0.035) and 2|3 (MCC = −0.009). Essentially uninformative for this outcome.

sMRI – Volumes (Rₛ = 0.142): Non-significant at 0|1 (MCC = 0.124) and 1|2 (MCC = 0.129); significantly negative at 2|3 (MCC = −0.054 **).

sMRI – Thickness (Rₛ = 0.081): Non-significant at 0|1 (MCC = 0.093) and 1|2 (MCC = 0.088); significantly negative at 2|3 (MCC = −0.044 *).

sMRI – rICV (Rₛ = 0.189): Non-significant at all boundaries individually, despite a nominally positive pattern across 0|1 (MCC = 0.109), 1|2 (MCC = 0.188), and 2|3 (MCC = 0.092).

All sMRI Features (Rₛ = 0.337 ***): Strongly significant at 0|1 (MCC = 0.311 **, ΔMCC = 0.115) and at 1|2 (MCC = 0.286 *, ΔMCC = 0.266); NA at 2|3. This is the only individual sMRI subtype to reach significance, likely because combining all features provides complementary information about the spatial distribution of white matter injury. Notably, All sMRI provides large incremental gains over Base Clinical at both boundaries, though these do not survive FDR correction for ΔMCC.

CSF Biomarkers (Rₛ = 0.049): Non-significant at 0|1 (MCC = 0.039); significantly negative at 1|2 (MCC = −0.067 ****); NA at 2|3. The CSF amyloid-tau signature is anti-correlated with progression from mild to moderate white matter rarefaction, consistent with its vascular rather than Alzheimer-type etiology.

White Matter Hyperintensities (Rₛ = 0.199 **): Non-significant at all three individual boundaries — 0|1 (MCC = 0.138), 1|2 (MCC = 0.124), 2|3 (MCC = 0.036) — despite reaching global significance. The global Rₛ reflects a consistent sub-threshold signal that does not concentrate at any single boundary.

Cognitive Composite Scores (Rₛ = 0.131 ****, ΔMCC = 0.111 ****): Significant at 0|1 (MCC = 0.153 ****, ΔMCC = 0.106 **) and 1|2 (MCC = 0.118 ****); non-significant at 2|3. Significantly improved upon matched Base Clinical predictions at the 0|1 boundary. One of the few feature sets with consistent positive signal across the two accessible boundaries.

DTI – Fractional Anisotropy (Rₛ = 0.103): Non-significant at 0|1 (MCC = 0.112) and 1|2 (MCC = 0.030); NA at 2|3. The weakest DTI metric for this outcome.

DTI – Axial Diffusivity (Rₛ = 0.339 *): Strongly significant at 0|1 (MCC = 0.507 **, ΔMCC = 0.265) — the highest MCC observed at any single class boundary across the entire supplemental analysis. Non-significant at 1|2 (MCC = 0.187) and non-significant at 2|3 (MCC = −0.025). Signal is concentrated entirely at the none-vs-mild boundary.

DTI – Mean Diffusivity (Rₛ = 0.344 *): Significant at 0|1 (MCC = 0.443 *, ΔMCC = 0.200); non-significant at 1|2 (MCC = 0.078) and 2|3 (NA). Second-highest MCC at the 0|1 boundary.

DTI – Radial Diffusivity (Rₛ = 0.363 *): Significant at 0|1 (MCC = 0.397 *, ΔMCC = 0.155); non-significant at 1|2 (MCC = 0.301) and 2|3 (NA). Strong signal at the none-vs-mild boundary only.

Summary. White Matter Rarefaction stands out as the most distinctive outcome in the study. Base Clinical was essentially non-informative. DTI diffusivity metrics (AD, MD, RD) dominated the 0|1 boundary with the three highest class-specific MCCs observed across all outcomes and boundaries — a reflection of the unparalleled sensitivity of diffusion microstructure to the earliest emergence of white matter injury. All sMRI Features provided meaningful signal at both the 0|1 and 1|2 boundaries. Cognitive Composite Scores contributed consistent modest signal at both lower boundaries and significantly exceeded Base Clinical at 0|1. CSF Biomarkers were paradoxically negative at 1|2. The severe boundary (2|3) was inaccessible to most feature sets due to sparse class representation. Overall, WMR illustrates a case where DTI microstructure, not conventional clinical or volumetric measures, drives prediction — and only for the earliest stage of injury.

**Supplemental Discussion**

**S5.2 Class-Specific Prediction: Insights from Ordinal Boundary Decomposition**

The class-specific binary decomposition of ordinal outcomes, reported in Supplemental Results 5.2, reveals a level of granularity not visible in global Spearman rank correlations alone. For several outcomes, feature sets that appeared broadly informative at the global level showed a predictive signal that was entirely concentrated at one or two class boundaries, while performing at or below chance at others. This pattern has direct implications for how biomarker utility should be interpreted and communicated to the field. Below we discuss the class-specific findings for each neuropathological domain in the context of current research.

**S5.2.1 Amyloid Pathology: The Gradient from Earliest to Most Advanced Deposition**

For Thal Phase, the class-specific analysis revealed that Cognitive Composite Scores was the only non-CSF feature set to achieve significant predictions across all five ordinal boundaries. CSF biomarkers, despite producing the highest global Rₛ (0.666), failed entirely at the earliest boundary (Thal 0|1, MCC = 0.065, ns), recovering strongly from the 1|2 boundary onward. This pattern is consistent with the proposed temporal ordering of biomarker changes in the AD cascade, in which CSF Aβ42 becomes detectable before cognitive impairment but only after sufficient neocortical deposition has occurred [Jack et al., 2013]. It also aligns with recent data from the A4 trial and related pre-symptomatic cohorts, where CSF Aβ42 suppression is more reliably detected at moderate-to-high amyloid burden than at the threshold of earliest deposition [Sperling et al., 2023]. The failure of all structural and diffusion MRI feature sets at the earliest Thal boundaries is consistent with the late position of atrophy in the hypothetical biomarker model [Jack et al., 2010; Jack et al., 2013], in which volumetric MRI changes trail CSF and amyloid PET by many years.

For A Score, the critical finding was that CSF biomarkers provided the only significant improvement over Base Clinical at the severe amyloid boundary (2|3: ΔMCC = 0.340 *), while no feature set discriminated none-vs-any (0|1) amyloid with meaningful accuracy. This dissociation — poor sensitivity to the earliest amyloid accumulation but strong performance at advanced burden — has been noted in comparable studies using PET-validated cohorts and is likely to reflect the lower CSF Aβ42/Aβ40 ratio sensitivity at the Thal 1 threshold [Hansson et al., 2006; Ossenkoppele et al., 2022].

The CERAD outcome showed the broadest and most uniform feature set coverage of any amyloid measure, with CSF biomarkers significantly exceeding Base Clinical at both the 0|1 and 1|2 boundaries — a result not replicated for Thal Phase or A Score. CERAD's emphasis on plaque density in neocortical regions, rather than the spatial progression captured by Thal staging, may make it more sensitive to the combined signal of Aβ42 suppression and tau elevation at intermediate stages of disease [Serrano-Pozo et al., 2011]. WMH and Cognitive Composite Scores provided consistent coverage across all three CERAD boundaries, suggesting that neuritic plaque burden is reflected both in white matter signal changes and in domain-specific cognitive decline.

**S5.2.2 Tau Pathology: Early-Stage Failure and the Uniqueness of Cognitive Breadth**

For Braak Stage, the class-specific analysis exposed a striking asymmetry: sMRI feature sets (Thickness, All sMRI) produced significantly negative MCCs at the earliest tau boundaries (0|1 and 1|2), indicating anti-correlated predictions in the small sub-cohort. This artifact arises from the near-absence of structural atrophy at transentorhinal Braak stages I–II, such that the model learns to associate cortical volume with the absence of early tau pathology — an inversion that reflects appropriate biology but a counterproductive cross-validation artifact at very low sample size [Varoquaux et al., 2018]. Importantly, Cognitive Composite Scores avoided this failure entirely and was the only feature set significantly predictive at all six Braak boundaries, with statistically significant incremental improvements over Base Clinical at the two highest boundaries (4|5 and 5|6). This is consistent with growing evidence that domain-specific memory and executive function scores capture neocortical tau spread at limbic and isocortical stages before frank dementia is clinically apparent [Brier et al., 2016; Schultz et al., 2017].

For B Score, the class-specific results produced the most uniformly significant set of results across the entire study: at the 2|3 boundary (low-to-intermediate vs. high tau), all available feature sets — including all four DTI metrics — reached significance simultaneously. This is the only outcome-by-boundary pairing where DTI contributed meaningful signal jointly with clinical, cognitive, CSF, and sMRI measures. This convergence at the high-tau boundary is biologically interpretable: advanced neocortical neurofibrillary pathology (Braak V–VI) is accompanied by widespread axonal loss and myelin breakdown that reduces white matter integrity across multiple tracts [Acosta-Cabronero et al, 2012; Weston et al., 2015]. The Cognitive Composite Scores feature set was the only one to significantly exceed matched Base Clinical predictions at this boundary, reinforcing its value as a globally sensitive predictor even for tau pathology.

**S5.2.3 Cardiovascular Disease: Boundary-Specific Dissociations and the CSF Paradox**

For arteriolosclerosis, CSF biomarkers produced the single largest class-specific incremental improvement over Base Clinical observed across all outcomes and all boundaries in the study (0|1: ΔMCC = 0.391 ***), while DTI diffusivity metrics (MD, RD, AD) jointly significant at both the 0|1 and 1|2 boundaries. Yet at the severe boundary (2|3), no feature set performed above chance. This floor effect at the severe end of the arteriolosclerosis scale likely reflects the rarity, pathological heterogeneity, and potential confounding by concurrent severe AD pathology that characterizes grade 3 small-vessel disease [Pasi et al., 2015; Kapasi et al., 2017]. The strong CSF signal at the none-vs-any boundary — concentrated in early arteriolosclerosis — is a somewhat unexpected finding, as CSF Aβ42 and tau do not directly reflect small-vessel disease severity. It may instead reflect the well-documented co-occurrence of early amyloid accumulation and arteriolar disease in aging brains, as shared vascular risk factors (hypertension, diabetes, APOE ε4) promote both pathological processes simultaneously [Toledo et al., 2013; Iadecola, 2013; Iadecola et al., 2016]. The paradoxically negative CSF prediction at the none-vs-any atherosclerosis boundary (MCC = −0.068 ****) reinforces this interpretation: the amyloid-tau CSF profile may actually be inversely associated with large-vessel disease burden at the earliest threshold, consistent with known differences in the demographic and genetic risk factor profiles of small- versus large-vessel disease.

For CAA, the class-specific findings reconfirm the unique specificity of CSF amyloid-tau analytes to this outcome — providing significant predictions at the 0|1 and 1|2 boundaries — but the near-total failure of structural and diffusion MRI, WMH, and DTI at all CAA boundaries is notable. WMH features are frequently used as imaging surrogates of CAA in clinical and research settings [Charidimou et al., 2017], but here they were non-significant at all three CAA boundaries. This null result may reflect the broad heterogeneity of the WMH measures included in our harmonized feature set, which capture white matter signal abnormality from both small-vessel disease and perivascular amyloid accumulation without molecular specificity. More targeted CAA biomarkers — such as cortical microbleeds on susceptibility-weighted imaging, convexal subarachnoid hemorrhage, or cortical superficial siderosis — were not available in the harmonized dataset but would be expected to provide boundary-specific signal here [Greenberg et al., 2018].

**S5.2.4 Lewy Body Pathology: Irreducible Difficulty of Fine-Grained Staging**

For PD Braak Stage, the class-specific results confirm the near-total failure of all feature sets to predict fine-grained Lewy body staging. CSF biomarkers and Cognitive Composite Scores provided the only positive class-level signal, and exclusively at the two lowest boundaries (0|1 and 1|2). The 3|4 boundary was inaccessible to most feature sets due to insufficient class representation, and the 2|3 boundary was predicted only by Cognitive Composite Scores with marginal MCCs (0.082 **** — significant but very small). This pattern is consistent with the biological reality that PD Braak staging captures a spatial propagation of α-synuclein pathology– the CSF panels used here contained no direct α-synuclein surrogate, and currently, no available antemortem imaging modality — including structural or diffusion MRI — can directly detect α-synuclein inclusion bodies with the molecular specificity needed to distinguish Braak stages 1 through 4. [Alzghool et al., 2022; Attems et al., 2021]. Future incorporation of αSyn-SAA into the CSF feature set — now demonstrating sensitivity of approximately 86% and specificity of 92% in Parkinson's disease diagnosis [Zheng et al., 2023; Espay et al., 2025] — could substantially alter the class-specific prediction landscape for Lewy body outcomes.

The L Score findings presented a more accessible target: CSF Biomarkers provided significant predictions at both boundaries, and WMH, Cognitive Composite Scores, rICV, and All sMRI Features also reached significance at the lower boundary. The condensation of 5-class PD Braak staging into a 3-class L Score improved discriminability, consistent with a general principle that binary and lower-granularity ordinal outcomes are inherently more predictable from non-specific biomarker features than fine-grained staging systems designed for pathological precision [Sperling et al., 2011].  But from these results we can see that the drop in performance across the feature sets for the PD Braak staging is at the later stages, specifically PD Braak 3 & PD Braak 4, both of which are condensed in the L-Score with PD Braak 2.  Nearly every feature set failed to even predict a single instance of PD Braak Stage 4 (and the subsequent binarized MCC calculation on that confusion matrix will always be NA.)  Only the base clinical and Cognitive Composite Scores on PD Braak Stage 4 produced scorable confusion matrices, but both of them performed badly on this measure.  Only Cognitive Composite Scores predictions for PD Braak Stage 3 were significant if very weak (MCC = 0.082 ****).

**S5.2.5 TDP-43 Staging: Cognitive Composite Scores as the Sole Informative Feature Set**

The class-specific analysis for TDP-43 5-Stage revealed the most alarming pattern of prediction failure in the study. DTI-AD (Rₛ globally = −0.277) showed significantly negative MCCs at the 2|3 and 3|4 boundaries (MCC = −0.293 **** and −0.279 ****), indicating strong anti-correlation between predicted and true class labels at intermediate stages. sMRI Volumes and All sMRI Features were similarly negative at intermediate boundaries. These are not informative anti-correlations but model instability artifacts arising from insufficient sample sizes and sparse class occupancy across the six TDP-43 stages — a well-described failure mode of gradient boosting and ensemble methods in highly imbalanced, small-sample regimes [Blagus & Lusa, 2010; Johnson & Khoshgoftaar, 2019]. Only Cognitive Composite Scores provided any positive signal, and exclusively at the four lower boundaries. This outcome underscores the need for larger harmonized datasets for TDP-43 neuropathology prediction and the development of TDP-43-specific fluid biomarkers [Nelson et al., 2019; Wolk et al., 2025].

For TDP-43 3-Stage, the condensed staging system substantially improved predictability: rICV reached significance at all three boundaries, and Cognitive Composite Scores — the only feature set to significantly improve upon Base Clinical — showed consistent signal across all three. The class-specific incremental improvement at the 1|2 boundary (ΔMCC = 0.073 **) is consistent with the hypothesis that domain-specific cognitive profiles reflect limbic TDP-43 staging as it progresses beyond the amygdala [Hiya et al., 2024; Sajjadi et al., 2025]. The subthreshold but numerically suggestive CSF pattern (approaching significance at all three boundaries) warrants further investigation in larger cohorts, as some emerging evidence suggests that CSF neurofilament light and glial markers may co-vary with LATE-NC stage independently of Alzheimer co-pathology [Wolk et al., 2025].

**S5.2.6 White Matter Rarefaction: DTI Microstructure at the Earliest Boundary**

White Matter Rarefaction produced the most distinctive class-specific profile in the supplemental analysis. At the none-vs-mild boundary (0|1), DTI-AD achieved an MCC of 0.507 (*) — the highest class-specific MCC observed at any single boundary across the entire study — followed by DTI-MD (0.443 *) and DTI-RD (0.397 *). This concentration of DTI signal at the earliest WMR threshold, with near-complete collapse at the 1|2 boundary and inaccessibility at 2|3, indicates that white matter microstructure measured by diffusion MRI is most sensitive to the transition from intact to minimally injured white matter, rather than to the progression through successive degrees of established injury. This finding aligns with the known sensitivity of diffusivity metrics to early periarteriolar rarefaction and subtle myelin water content changes that precede the macroscopic white matter signal changes visible on FLAIR [Maillard et al., 2012; Wardlaw et al., 2013]. The complementary performance of All sMRI Features at both the 0|1 and 1|2 boundaries — providing the only structural feature set with significant predictions across two WMR thresholds — suggests that multi-regional volumetric information partially compensates for what diffusivity measures lose at the second boundary. The paradoxically negative CSF prediction at the 1|2 boundary (MCC = −0.067 ****) further reinforces that the amyloid-tau CSF profile is orthogonal or mildly inversely related to the vascular aetiology of progressive white matter rarefaction, which is driven by hypoperfusion and arteriolar hyalinization rather than amyloid accumulation [Beach et al., 2023].

The near-total uninformativeness of Base Clinical at the WMR boundaries — the only outcome, along with CVD Any Binary, where Base Clinical was globally non-significant — deserves emphasis. Sex, age at death, clinical diagnosis, and the interval between assessment and death, which collectively explained 24 of 26 outcomes at a global level, carried essentially no information about WMR stage. This dissociation suggests that WMR is clinically underascertained during life, and that imaging rather than clinical phenotyping is required for its inference. The strong performance of Cognitive Composite Scores at the 0|1 boundary, and particularly its statistically significant improvement over Base Clinical, provides an interim path: domain-specific cognitive profiling captures white matter pathology more sensitively than clinical staging alone, consistent with evidence that subcortical white matter injury produces characteristic executive and processing speed deficits even before clinical diagnostic thresholds are reached [Sachdev et al., 2014; Kearney-Ramos et al., 2022].

**S5.2.7 General Principles Arising from Class-Specific Analysis**

Several cross-cutting principles emerge from the class-specific decomposition. First, boundary-specific concentration of signal is the rule rather than the exception: most feature sets that appear moderately informative at the global Rₛ level carry their predictive information in only one or two class boundaries, typically the most severe end of each staging scale. This limits the clinical applicability of single-feature-set models for early-stage detection and motivates the development of sequential or staged diagnostic frameworks that use different feature sets at different points in the disease continuum. Second, the severe boundary (class K-1 | K for K-class outcomes) is systematically the most predictable by all feature sets — but also the most subject to ascertainment bias, as individuals who reach autopsy with advanced neuropathology are more likely to have been clinically diagnosed and longitudinally followed by specialized research centres. The high MCC values at the most advanced boundaries should therefore be interpreted with this selection pressure in mind [Iadecola et al., 2016; Jack et al., 2024].

Third, the DTI anti-correlations observed at early Thal and Braak boundaries, and the TDP-43 5-Stage intermediate boundaries, are methodologically important: they illustrate how multi-model ensemble methods such as AutoGluon can produce spuriously negative predictions when class representation is insufficient for stable model learning. These artefacts are particularly hazardous in clinical translation contexts, as a model returning a negative MCC at a specific threshold performs worse than a random classifier. Future work applying these models to prospective clinical populations should implement minimum class representation checks before deploying ordinal boundary predictions. Finally, Cognitive Composite Scores stands out as the only feature set to provide both broad boundary coverage and statistically significant incremental improvement over Base Clinical at individual class boundaries for multiple outcomes — including Braak Stage, B Score, CAA, TDP-43 3-Stage, and WMR. These results collectively argue for prioritizing the harmonized cognitive domain framework developed by the ADSP-PHC Cognition Core [Mukherjee et al., 2023; Kang et al., 2025] as a broadly applicable antemortem proxy for multiple concurrent neuropathologies, particularly in cohorts where imaging and CSF data are unavailable.

**Supplemental References**

1. Jack CR Jr, Knopman DS, Jagust WJ, Petersen RC, Weiner MW, Aisen PS, et al. Tracking pathophysiological processes in Alzheimer's disease: an updated hypothetical model of dynamic biomarkers. Lancet Neurol. 2013 Feb;12(2):207–16. doi: 10.1016/S1474-4422(12)70291-0. PMID: 23332364.
2. Sperling RA, Donohue MC, Raman R, Rafii MS, Johnson K, Masters CL, et al. Trial of solanezumab in preclinical Alzheimer's disease. N Engl J Med. 2023 Sep 21;389(12):1096–107. doi: 10.1056/NEJMoa2305032. PMID: 37458272.
3. Ossenkoppele R, Pichet Binette A, Groot C, Smith R, Strandberg O, Palmqvist S, et al. Amyloid and tau PET-positive cognitively unimpaired individuals are at high risk for future cognitive decline. Nat Med. 2022 Nov;28(11):2381–7. doi: 10.1038/s41591-022-02049-x. PMID: 36357681.
4. Jack CR Jr, Knopman DS, Jagust WJ, Shaw LM, Aisen PS, Weiner MW, et al. Hypothetical model of dynamic biomarkers of the Alzheimer's pathological cascade. Lancet Neurol. 2010 Jan;9(1):119–28. doi: 10.1016/S1474-4422(09)70299-6. PMID: 20083042.
5. Hansson O, Zetterberg H, Buchhave P, Londos E, Blennow K, Minthon L. Association between CSF biomarkers and incipient Alzheimer's disease in patients with mild cognitive impairment: a follow-up study. Lancet Neurol. 2006 Mar;5(3):228–34. doi: 10.1016/S1474-4422(06)70355-6. PMID: 16488378.
6. Serrano-Pozo A, Frosch MP, Masliah E, Hyman BT. Neuropathological alterations in Alzheimer disease. Cold Spring Harb Perspect Med. 2011 Sep;1(1):a006189. doi: 10.1101/cshperspect.a006189. PMID: 22229116.
7. Varoquaux G. Cross-validation failure: small sample sizes lead to large error bars. Neuroimage. 2018 Jul 15;180(Pt A):68–77. doi: 10.1016/j.neuroimage.2017.06.061. PMID: 28655633.
8. Brier MR, Gordon B, Friedrichsen K, McCarthy J, Stern A, Christensen J, et al. Tau and Aβ imaging, CSF measures, and cognition in Alzheimer's disease. Sci Transl Med. 2016 May 11;8(338):338ra66. doi: 10.1126/scitranslmed.aaf2362. PMID: 27169802.
9. Schultz AP, Chhatwal JP, Hedden T, Mormino EC, Hanseeuw BJ, Sepulcre J, et al. Phases of hyperconnectivity and hypoconnectivity in the default mode and salience networks track with amyloid and tau in clinically normal individuals. J Neurosci. 2017 Apr 19;37(16):4323–31. doi: 10.1523/JNEUROSCI.3263-16.2017. PMID: 28314821.
10. Acosta-Cabronero J, Alley S, Williams GB, Pengas G, Nestor PJ. Diffusion tensor metrics as biomarkers in Alzheimer's disease. PLoS One. 2012;7(11):e49072. doi: 10.1371/journal.pone.0049072. PMID: 23145093; PMCID: PMC3493572.
11. Weston PSJ, Simpson IJA, Ryan NS, Ourselin S, Fox NC. Diffusion imaging changes in grey matter in Alzheimer's disease: a potential marker of early neurodegeneration. Alzheimers Res Ther. 2015 Oct 13;7(1):77. doi: 10.1186/s13195-015-0161-y. PMID: 26464218; PMCID: PMC4604104.
12. Wardlaw JM, Smith EE, Biessels GJ, Cordonnier C, Fazekas F, Frayne R, et al. Neuroimaging standards for research into small vessel disease and its contribution to ageing and neurodegeneration. Lancet Neurol. 2013 Aug;12(8):822–38. doi: 10.1016/S1474-4422(13)70124-8. PMID: 23867200.
13. Kapasi A, DeCarli C, Schneider JA. Impact of multiple pathologies on the threshold for clinically overt dementia. Acta Neuropathol. 2017 Aug;134(2):171–86. doi: 10.1007/s00401-017-1717-7. Epub 2017 May 9. PMID: 28488154.
14. Toledo JB, Arnold SE, Raible K, Brettschneider J, Xie SX, Grossman M, et al. Contribution of cerebrovascular disease in autopsy confirmed neurodegenerative disease cases in the National Alzheimer's Coordinating Centre. Brain. 2013 Sep;136(Pt 9):2697–706. doi: 10.1093/brain/awt188. Epub 2013 Jul 10. PMID: 23842566.
15. Iadecola C. The pathobiology of vascular dementia. Neuron. 2013 Nov 20;80(4):844–66. doi: 10.1016/j.neuron.2013.10.008. PMID: 24267647.
16. Iadecola C, Yaffe K, Biller J, Bratzke LC, Faraci FM, Gorelick PB, et al. Impact of hypertension on cognitive function: a scientific statement from the American Heart Association. Hypertension. 2016 Sep;68(6):e67–e94. doi: 10.1161/HYP.0000000000000053. PMID: 27977393; PMCID: PMC5065663.
17. Charidimou A, Boulouis G, Gurol ME, Ayata C, Bacskai BJ, Frosch MP, et al. Emerging concepts in sporadic cerebral amyloid angiopathy. Brain. 2017 Jul 1;140(7):1829–50. doi: 10.1093/brain/awx047. PMID: 28334869.
18. Greenberg SM, Charidimou A. Diagnosis of cerebral amyloid angiopathy: evolution of the Boston criteria. Stroke. 2018 Feb;49(2):491–7. doi: 10.1161/STROKEAHA.117.016990. Epub 2018 Jan 15. PMID: 29335334.
19. Alzghool OM, van Dongen G, van de Giessen E, Schoonmade L, Beaino W. α-Synuclein radiotracer development and in vivo imaging: recent advancements and new perspectives. Mov Disord. 2022 May;37(5):936–948. doi: 10.1002/mds.28984. Epub 2022 Mar 15. PMID: 35289424; PMCID: PMC9310945.
20. Attems J, Toledo JB, Walker L, Gelpi E, Gentleman S, Halliday G, et al. Neuropathological consensus criteria for the evaluation of Lewy pathology in post-mortem brains: a multi-centre study. Acta Neuropathol. 2021 Feb;141(2):159–172. doi: 10.1007/s00401-020-02258-7. PMID: 33392656; PMCID: PMC7847458.
21. Zheng Y, Li S, Yang C, Yu Z, Jiang Y, Feng T. Comparison of biospecimens for α-synuclein seed amplification assays in Parkinson's disease: a systematic review and network meta-analysis. Eur J Neurol. 2023 Dec;30(12):3949–67. doi: 10.1111/ene.16041. Epub 2023 Aug 28. PMID: 37573472.
22. Maillard P, Carmichael O, Fletcher E, Reed B, Mungas D, DeCarli C. Coevolution of white matter hyperintensities and cognition in the elderly. Neurology. 2012 Jul 31;79(5):442–8. doi: 10.1212/WNL.0b013e3182617136. Epub 2012 Jul 18. PMID: 22815562.

**Supplemental Figure Captions**

**Supplemental Figure 1: Thal Phase Class-Specific Predictive Capacity.** Black stars indicate a statistically significant prediction better than random chance (padj != 0).  Red stars indicate a statistical improvement of the participants in the given feature set over that of the predictions of the base clinical data alone.

**Supplemental Figure 2: A-Score Class-Specific Predictive Capacity.** Black stars indicate a statistically significant prediction better than random chance (padj != 0).  Red stars indicate a statistical improvement of the participants in the given feature set over that of the predictions of the base clinical data alone.

**Supplemental Figure 3: CERAD Class-Specific Predictive Capacity**. Black stars indicate a statistically significant prediction better than random chance (padj != 0).  Red stars indicate a statistical improvement of the participants in the given feature set over that of the predictions of the base clinical data alone.

**Supplemental Figure 4: AD Braak Stage Class-Specific Predictive Capacity.** Black stars indicate a statistically significant prediction better than random chance (padj != 0).  Red stars indicate a statistical improvement of the participants in the given feature set over that of the predictions of the base clinical data alone.

**Supplemental Figure 5: B-Score Class-Specific Predictive Capacity.** Black stars indicate a statistically significant prediction better than random chance (padj != 0).  Red stars indicate a statistical improvement of the participants in the given feature set over that of the predictions of the base clinical data alone.

**Supplemental Figure 6: Arteriosclerosis Class-Specific Predictive Capacity**. Black stars indicate a statistically significant prediction better than random chance (padj != 0).  Red stars indicate a statistical improvement of the participants in the given feature set over that of the predictions of the base clinical data alone.

**Supplemental Figure 7: Atherosclerosis Class-Specific Predictive Capacity.** Black stars indicate a statistically significant prediction better than random chance (padj != 0).  Red stars indicate a statistical improvement of the participants in the given feature set over that of the predictions of the base clinical data alone.

**Supplemental Figure 8: Cerebral Amyloid Angiopathy Class-Specific Predictive Capacity**. Black stars indicate a statistically significant prediction better than random chance (padj != 0).  Red stars indicate a statistical improvement of the participants in the given feature set over that of the predictions of the base clinical data alone.

**Supplemental Figure 9: PD Braak Stage (Lewy-Bodies) Class-Specific Predictive Capacity.** Black stars indicate a statistically significant prediction better than random chance (padj != 0).  Red stars indicate a statistical improvement of the participants in the given feature set over that of the predictions of the base clinical data alone.

**Supplemental Figure 10: L-Score Class-Specific Predictive Capacity**. Black stars indicate a statistically significant prediction better than random chance (padj != 0).  Red stars indicate a statistical improvement of the participants in the given feature set over that of the predictions of the base clinical data alone.

**Supplemental Figure 11: TDP-43 (5-Stage Progression) Class-Specific Predictive Capacity.** Black stars indicate a statistically significant prediction better than random chance (padj != 0).  Red stars indicate a statistical improvement of the participants in the given feature set over that of the predictions of the base clinical data alone.

**Supplemental Figure 12: TDP43 (3 Stage Progression) Class-Specific Predictive Capacity**. Black stars indicate a statistically significant prediction better than random chance (padj != 0).  Red stars indicate a statistical improvement of the participants in the given feature set over that of the predictions of the base clinical data alone.

**Supplemental Figure 13: White Matter Rarifaction Class-Specific Predictive Capacity.** Black stars indicate a statistically significant prediction better than random chance (padj != 0).  Red stars indicate a statistical improvement of the participants in the given feature set over that of the predictions of the base clinical data alone.
