## Supplementary figures and images for "Predicting Autopsy-Confirmed Neuropathology across Clinical, Neuroimaging, and CSF Biomarkers using Machine Learning"

### Sffigure 7

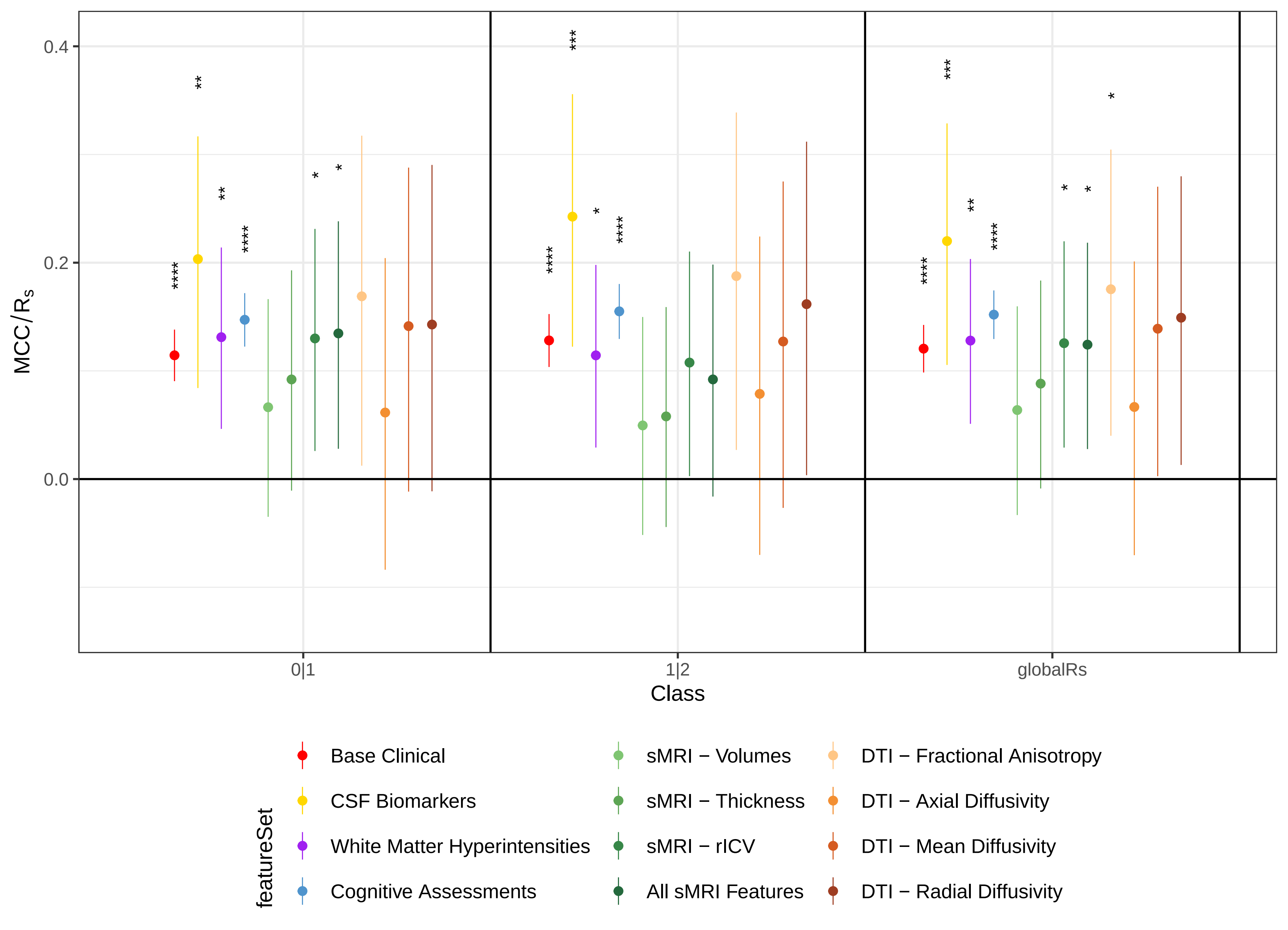

### SFigure 1

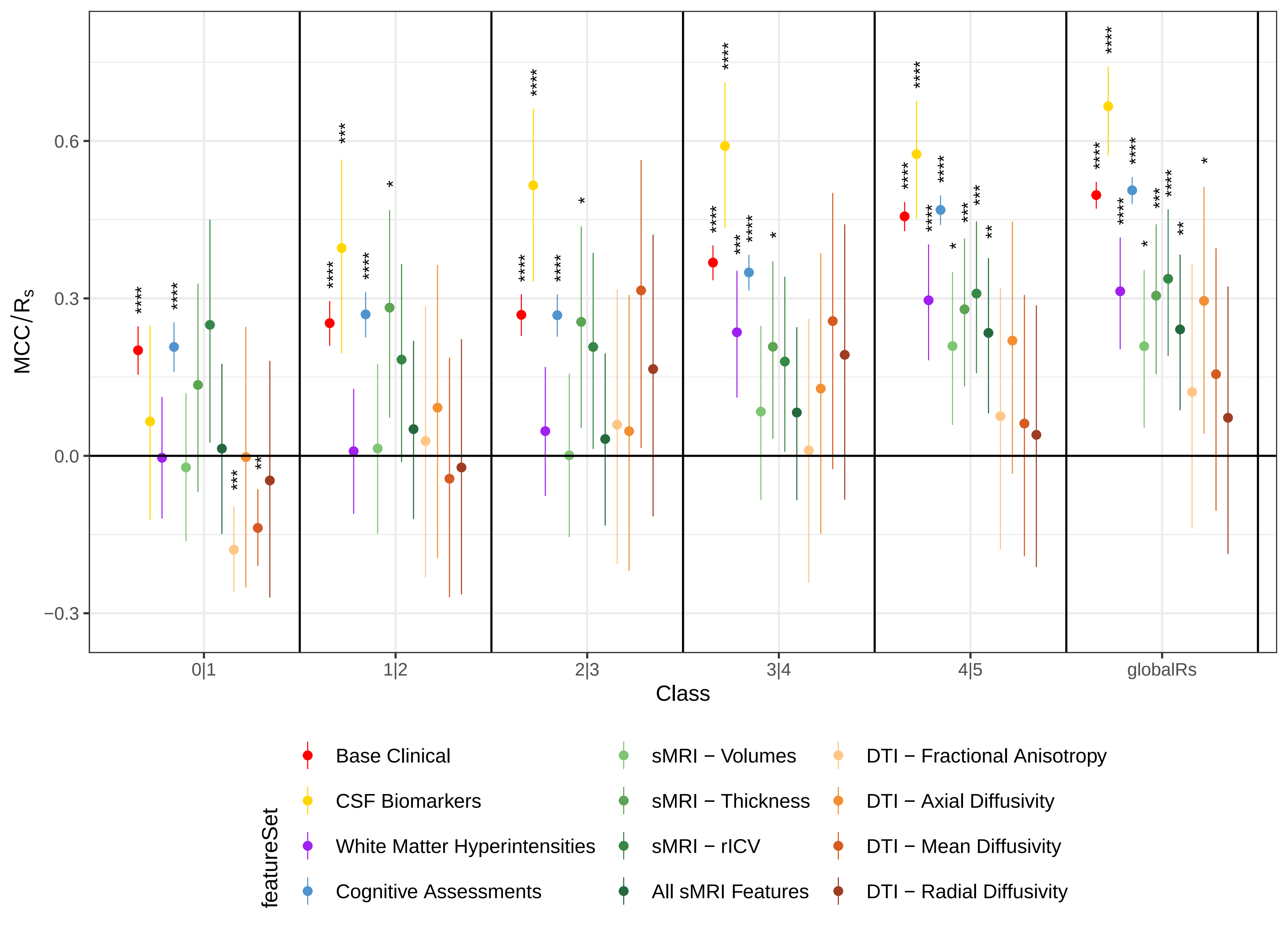

### SFigure 2

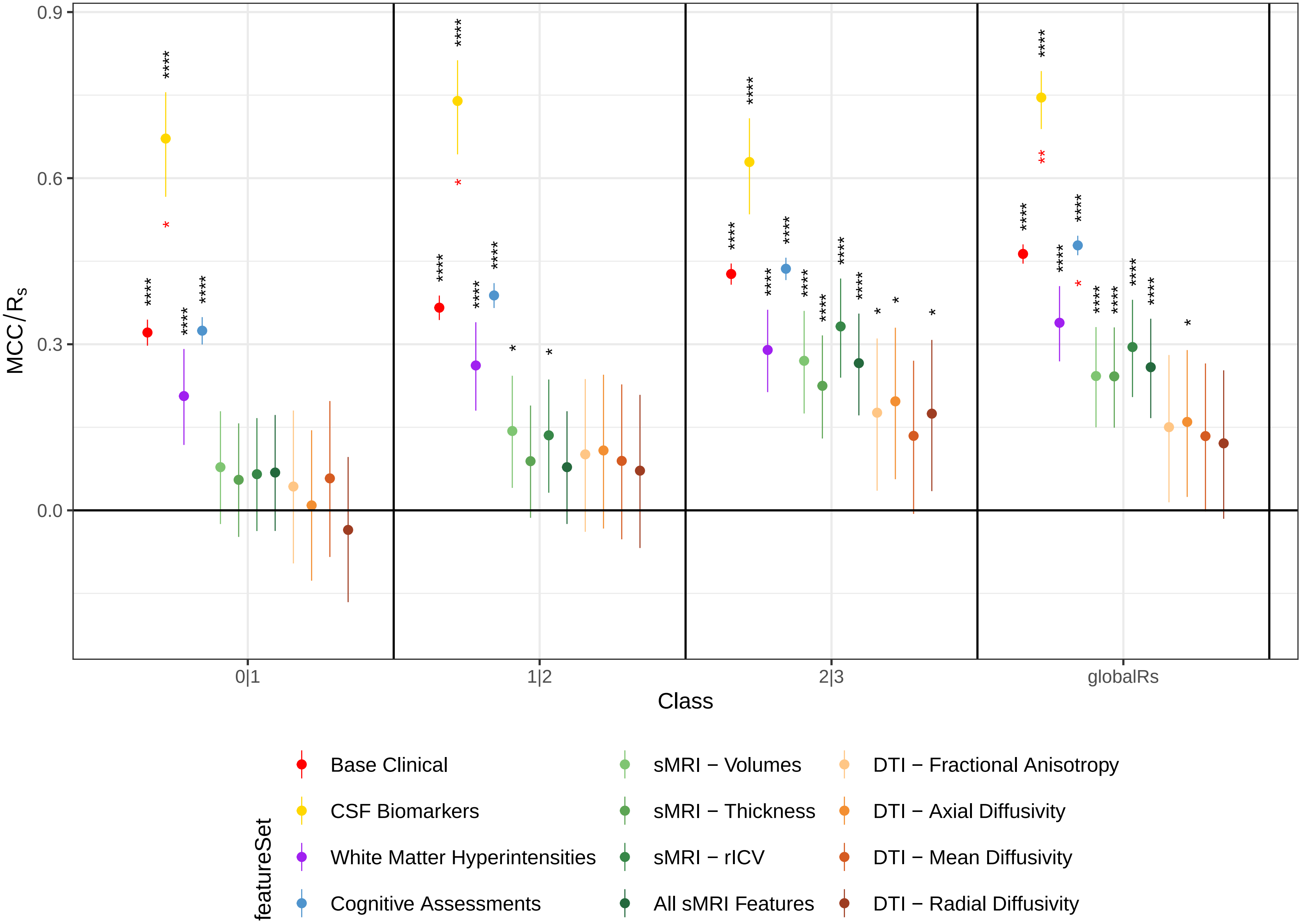

### SFigure 3

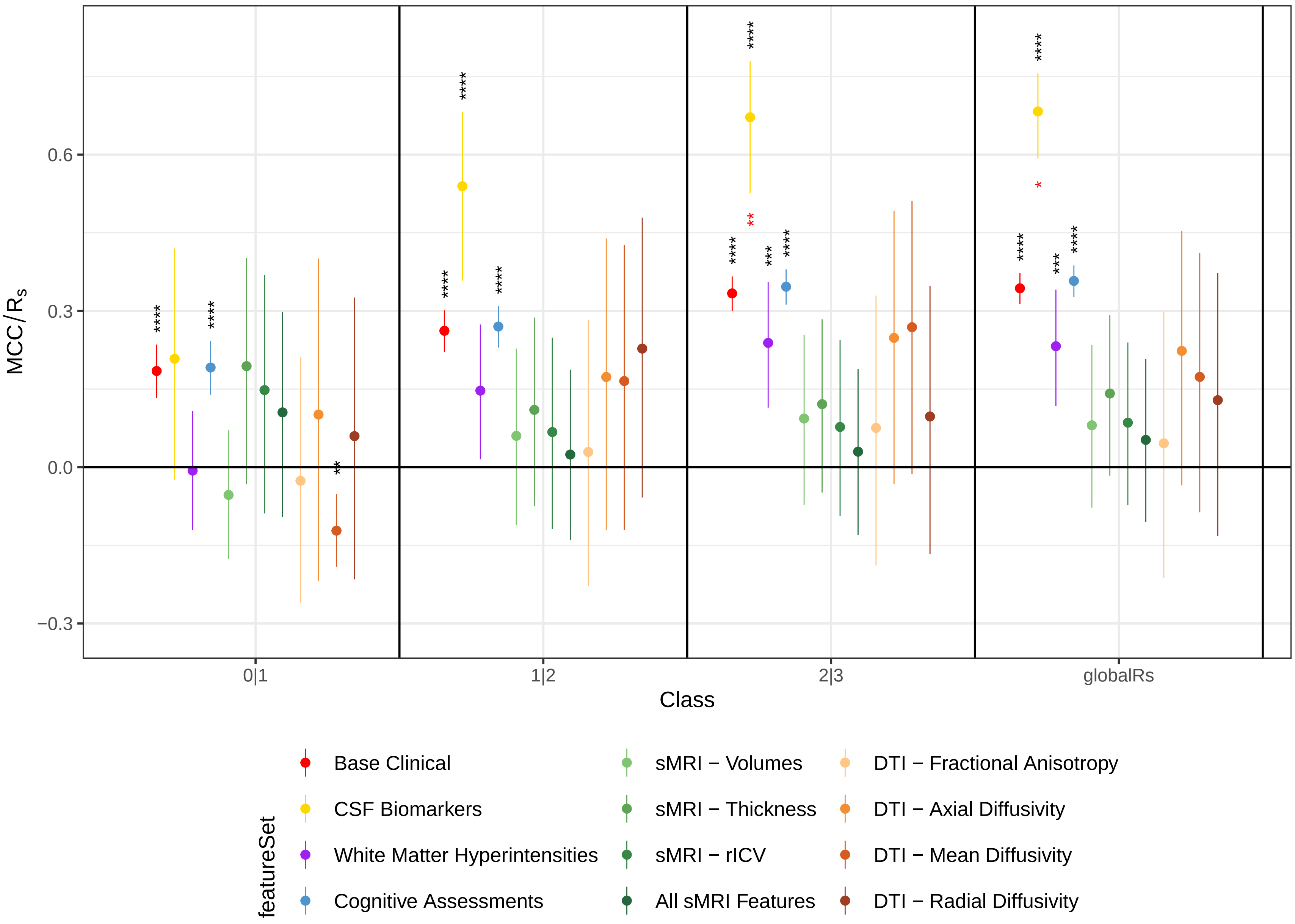

### SFigure 4

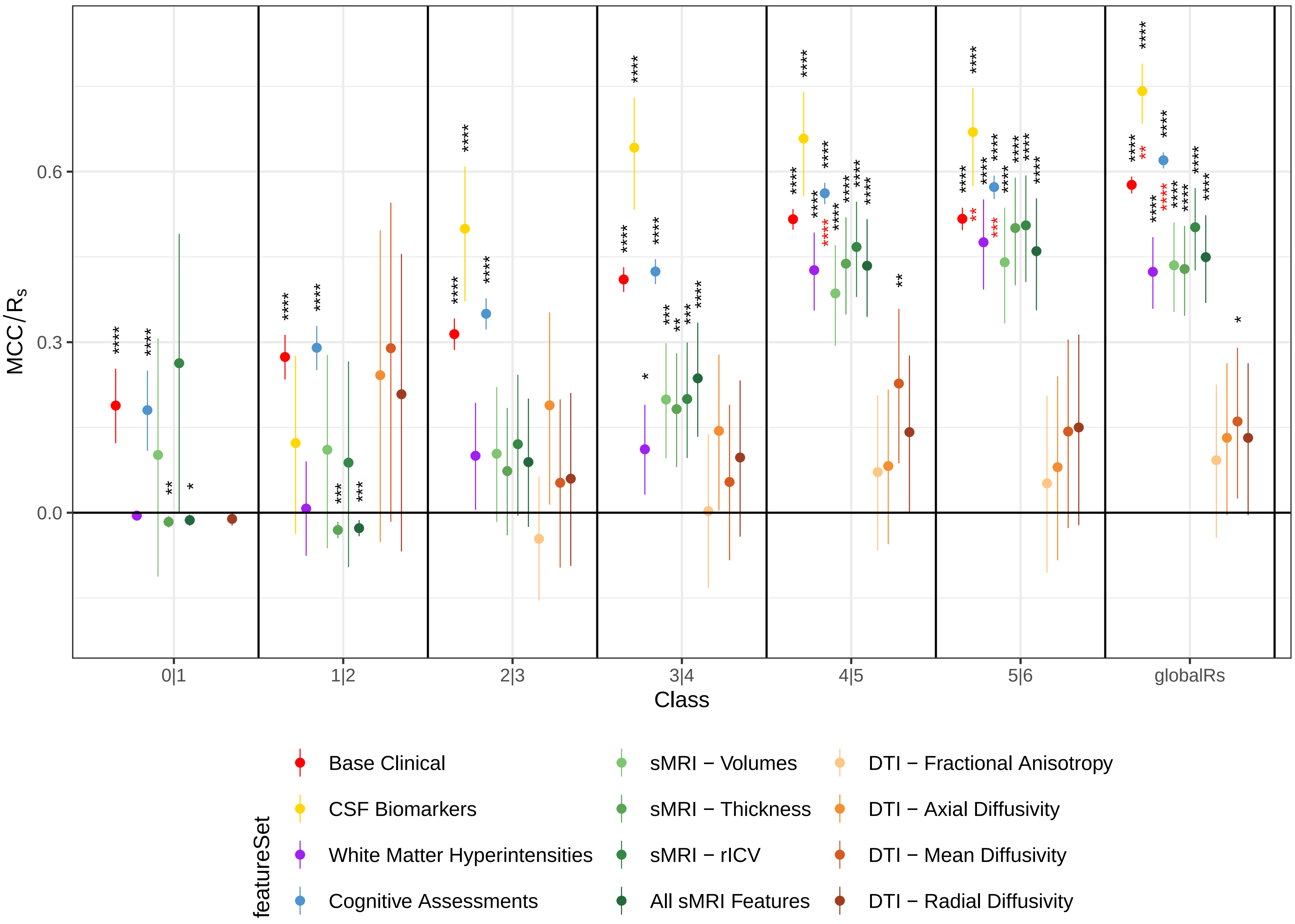

### SFigure 5

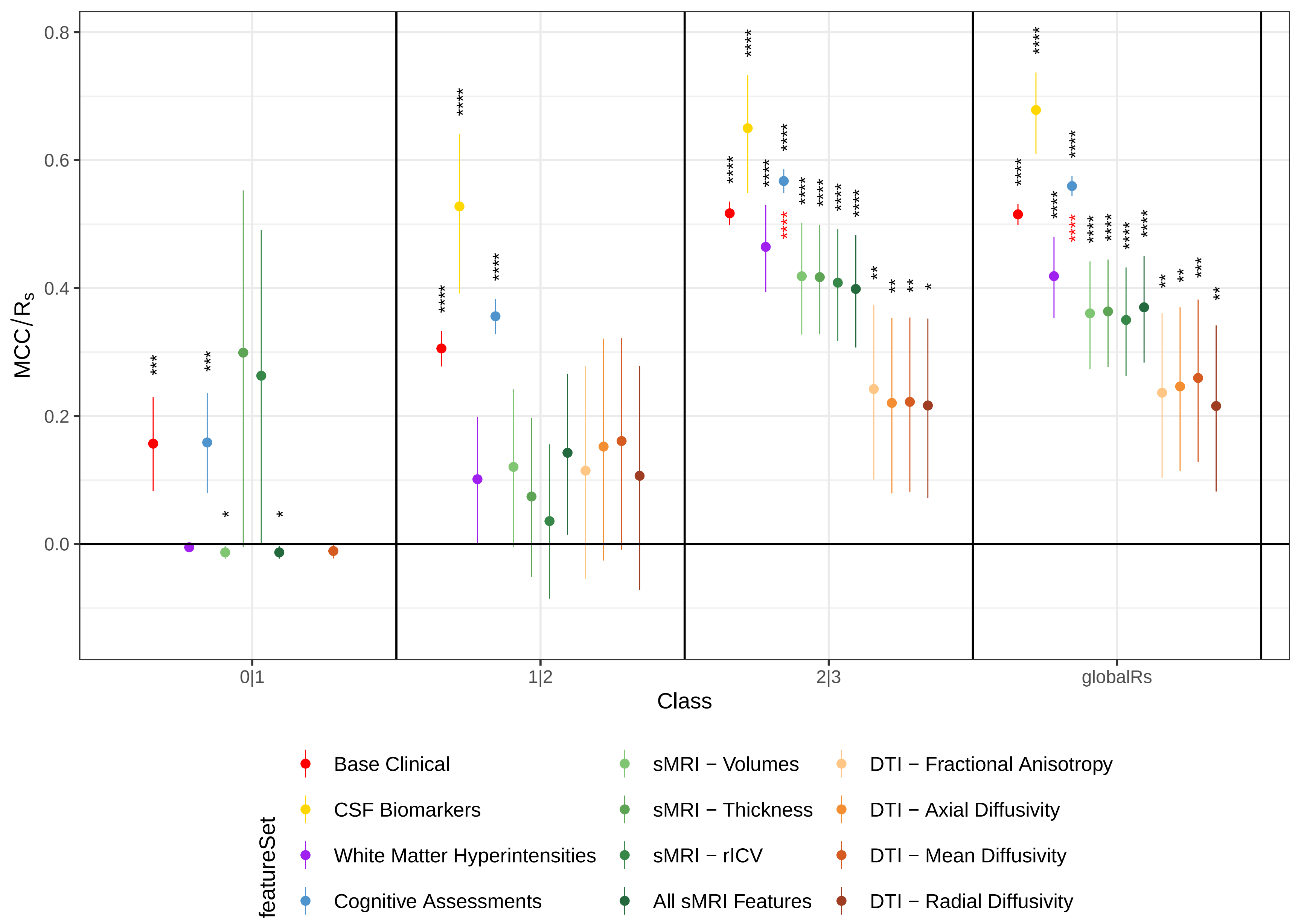

### SFigure 6

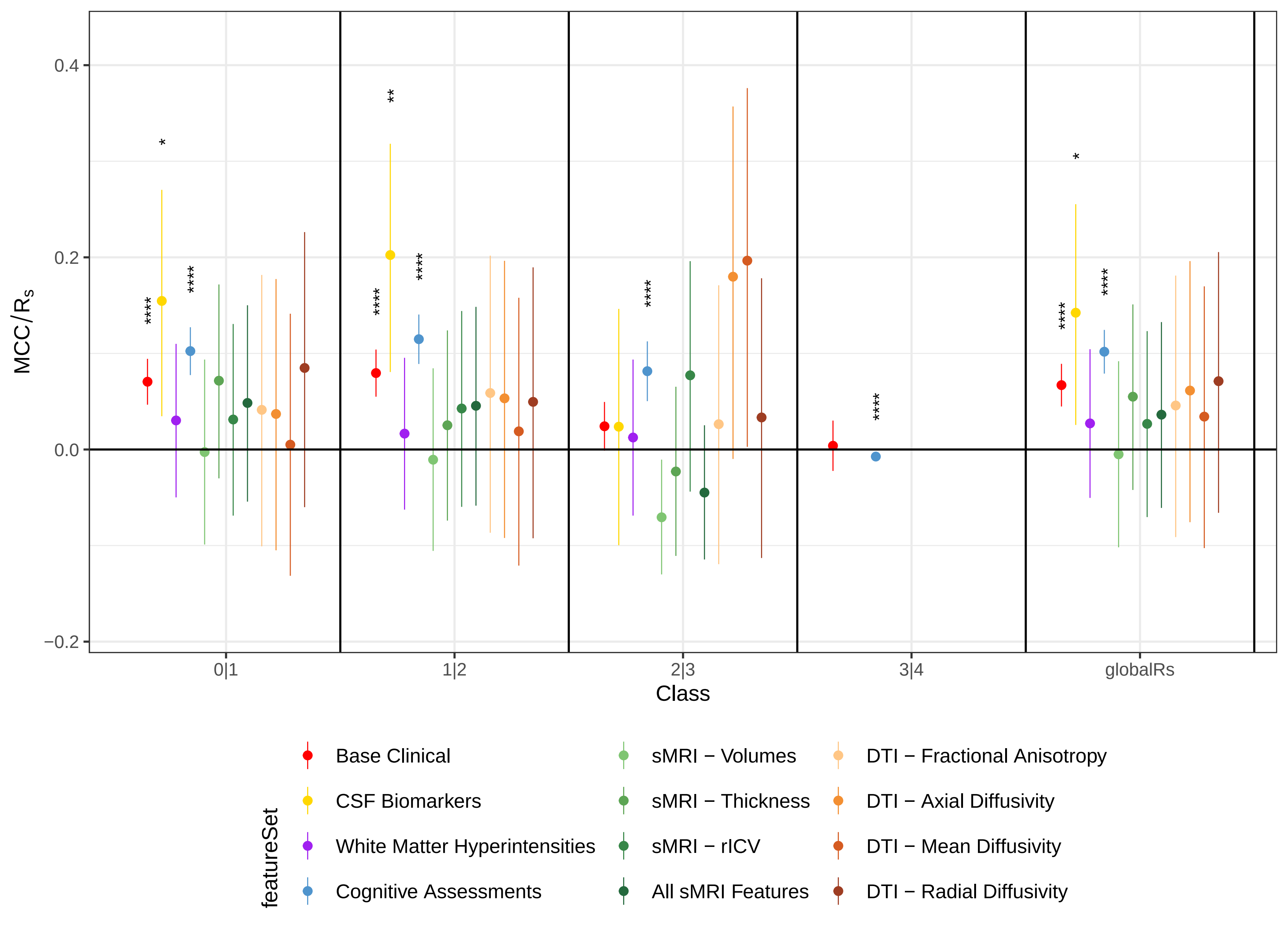

### SFigure 8

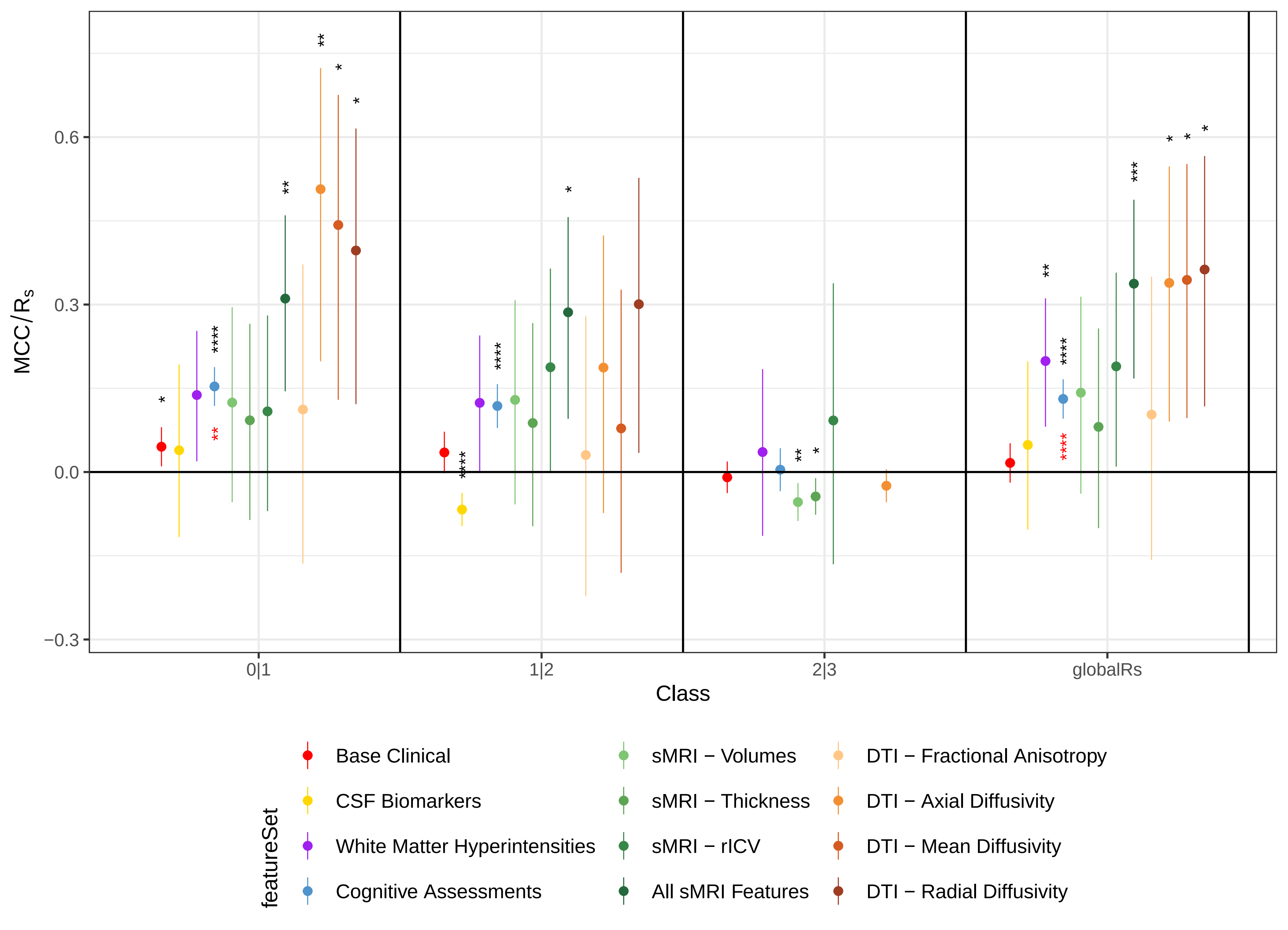

### SFigure 9

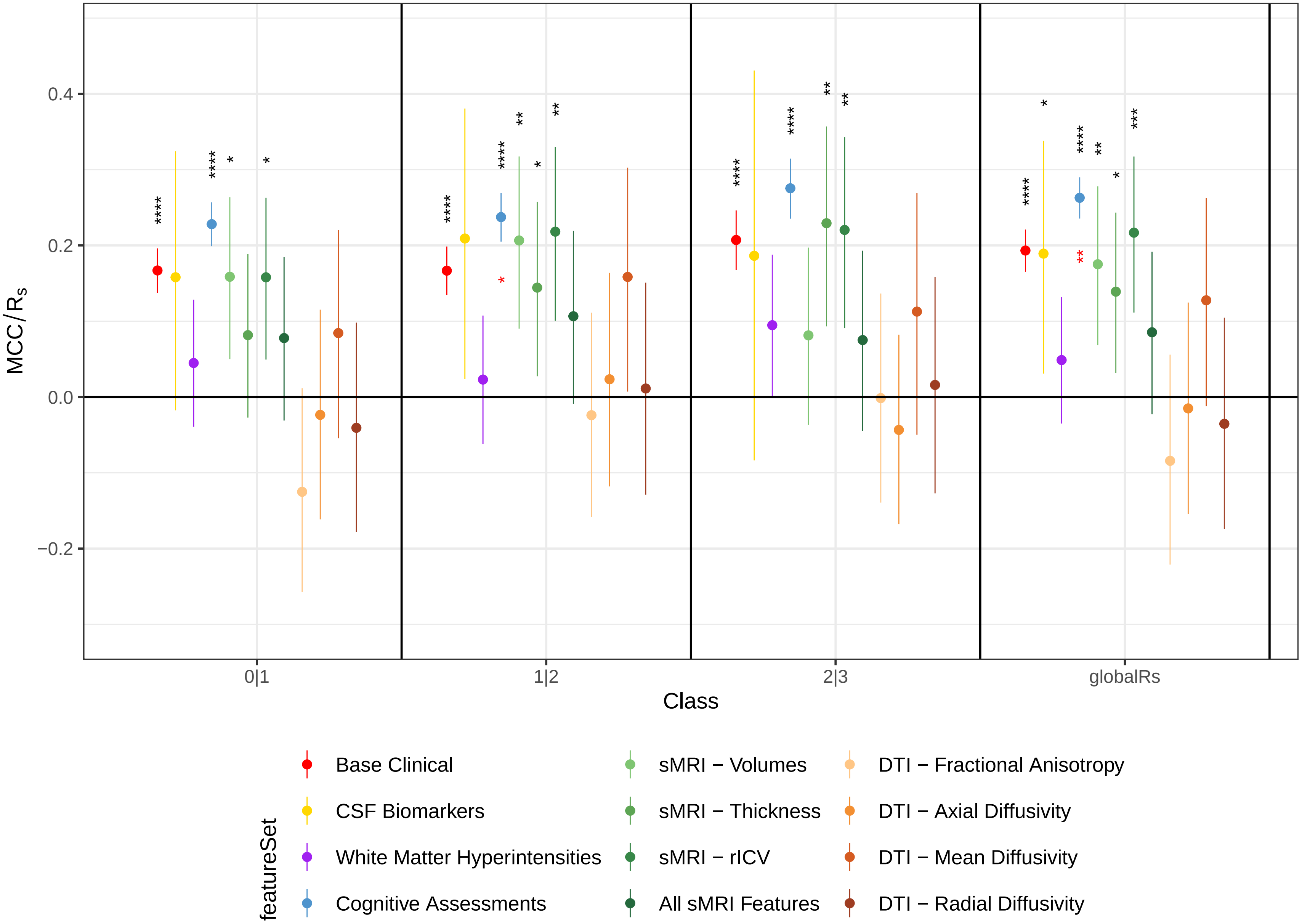

### SFigure 10

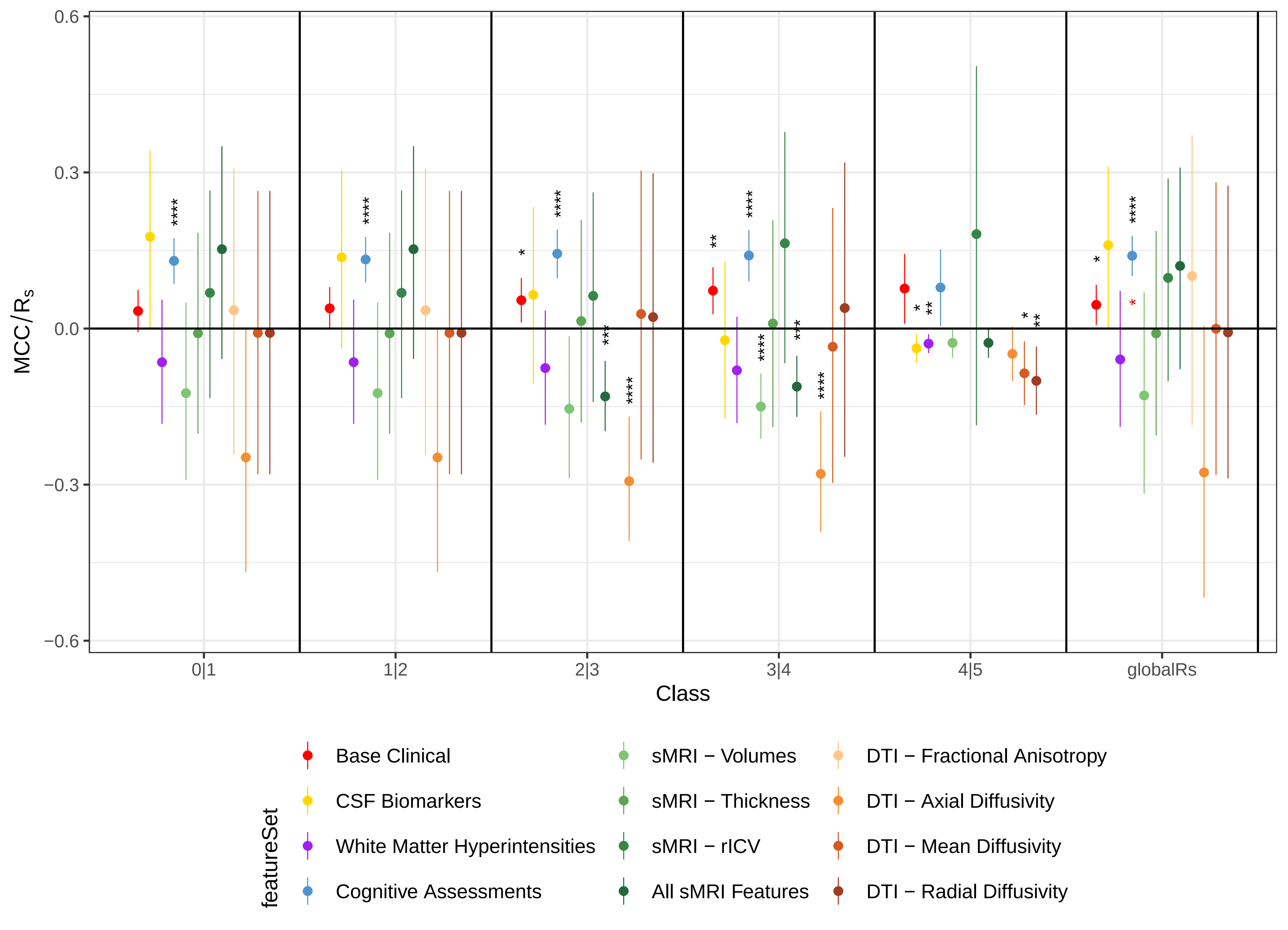

### SFigure 11

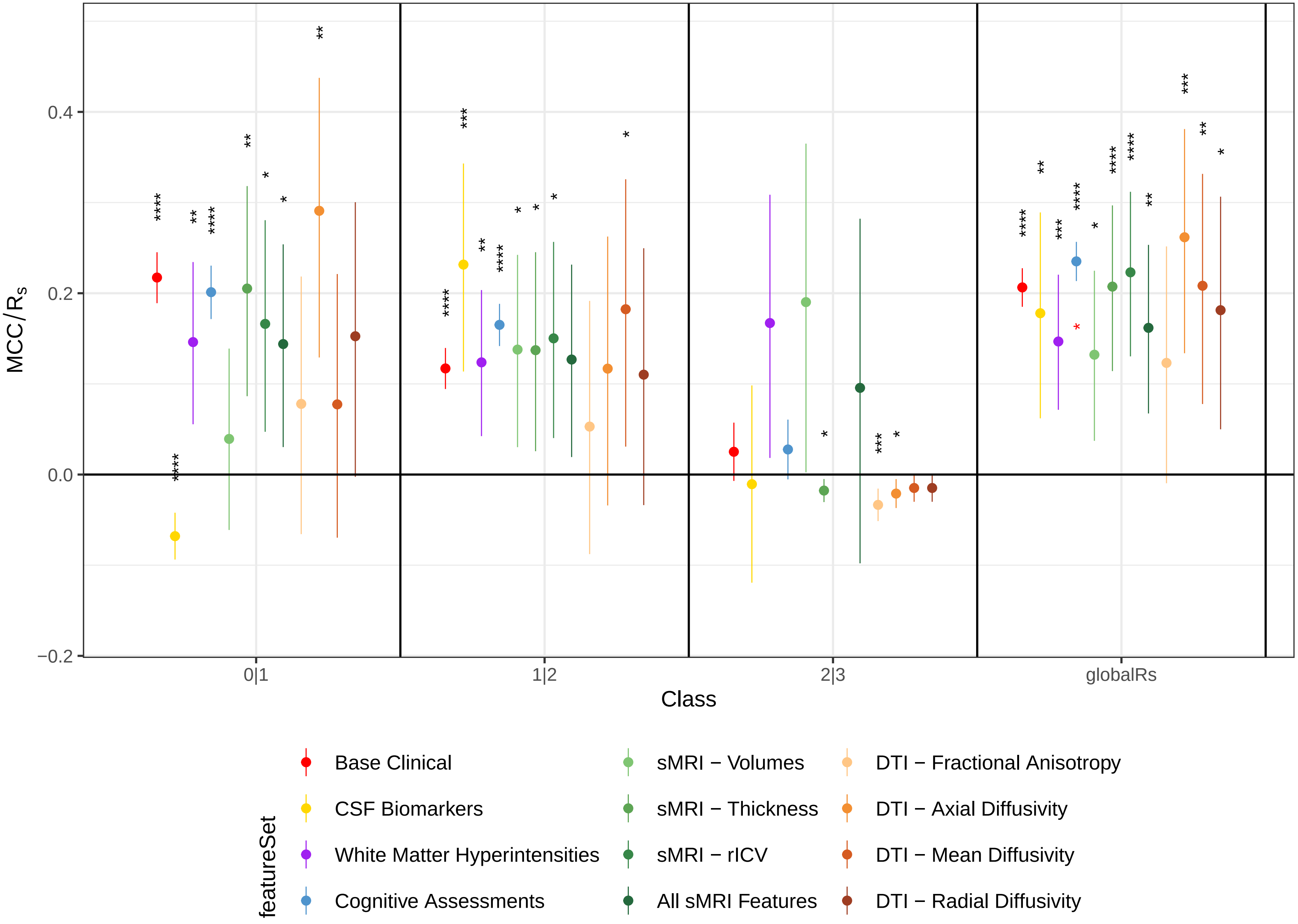

### SFigure 12

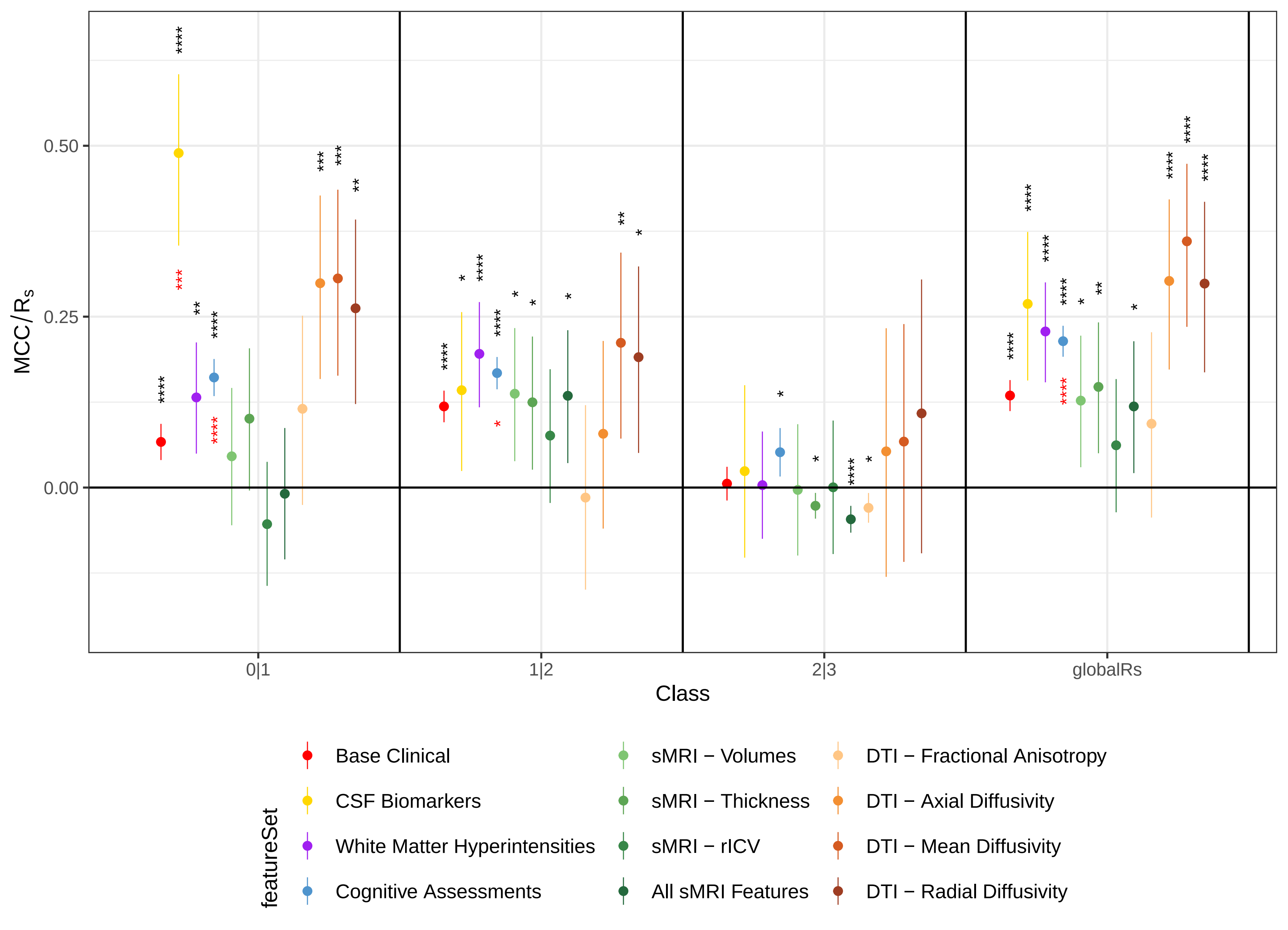

### Sfigure 13

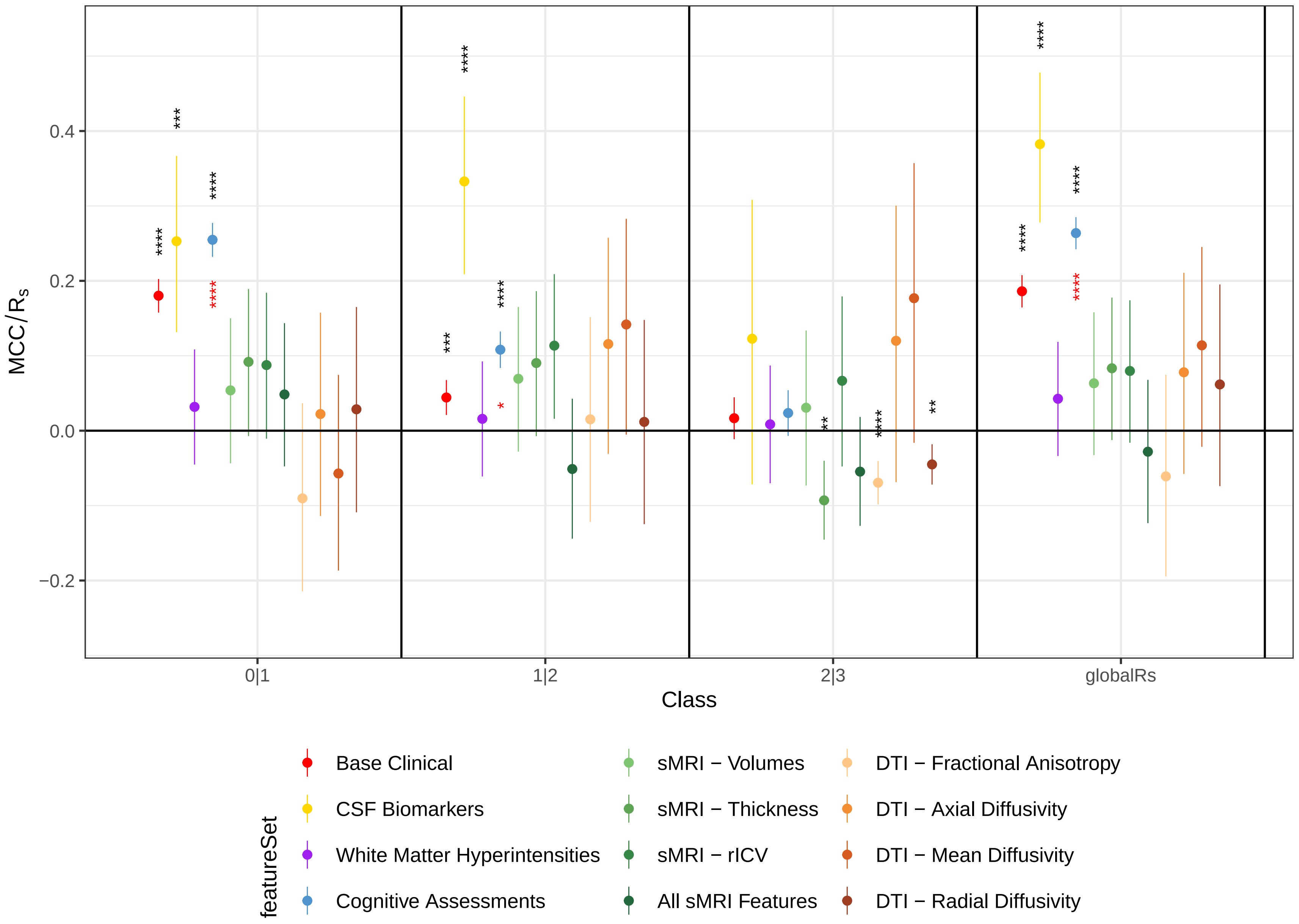
